# An alternative patterning mechanism for vertebrate tooth complexity

**DOI:** 10.64898/2026.03.12.711251

**Authors:** Daria Razmadze, Juuso Ikola, Ida-Maria Aalto, Nicolas Di-Poï

## Abstract

How development reliably generates complex patterns remains a central question in biology. Teeth provide an ideal system for addressing this problem, linking cusp-patterning mechanisms to diversification and the fossil record, yet the developmental basis of multicuspid crown patterning beyond mammals remains unresolved. In mammals, cusp initiation and spacing are controlled by transient epithelial signaling centers called enamel knots (EKs). Here, we combine comparative developmental analyses, 3D morphometrics, mutant perturbation, and Bayesian inverse modeling (BITES; Bayesian Inference for Tooth Emergence Simulation) to examine tooth morphogenesis across six squamate species spanning independent origins of multicuspid dentitions. We find that cusp patterning in these taxa does not rely on discrete EK organizers. Instead, cusps arise from a broad, persistent epithelial signaling field that deploys canonical tooth-patterning pathways but lacks the spatial restriction, apoptosis, and secondary organizers characteristic of the mammalian system. Elevated epithelial proliferation drives disproportionate expansion of this field during bud growth, enabling cusp addition. Quantitative variation in field extent and dynamics is sufficient to generate crown morphologies within and across species, providing a developmental basis for positional heterodonty and repeated gains and losses of multicuspidity in squamates. Our results identify an alternative epithelial patterning architecture for vertebrate cusp formation and suggest that the mammalian EK represents a derived, constrained implementation. Together, these findings revise how developmental complexity is encoded and show how shifts in patterning architecture can relax constraints and enable evolutionary convergence.

## INTRODUCTION

Explaining the evolution of complex morphologies requires linking developmental mechanisms to a comparative evolutionary framework. Teeth are a paradigmatic vertebrate organ whose exceptional adaptive diversity, tightly linked to diet, has driven the ecological and evolutionary success of numerous lineages. Their morphology ranges from simple conical crowns to elaborate architectures bearing multiple cusps and accessory structures that enhance functional performance. Although the developmental basis of mammalian tooth complexity is well characterized, comparable mechanisms in non-mammalian vertebrates remain largely unresolved, despite repeated independent origins of complex dentitions in fishes (1–5), amphibians (6, 7), and reptiles (8–10). Consequently, we lack a unifying framework explaining how crown elaborations such as cusps arise outside mammals, how patterning mechanisms vary across lineages, and how similar morphologies evolve convergently.

A major barrier to resolving these questions is the assumption, derived largely from a small number of mammalian models (11), that cusp formation relies on a highly stereotyped and tightly regulated signaling center, the EK. In mammals, cusp patterning is initiated by a primary EK at the cap stage, followed by secondary EKs that mark future cusp sites during the bell stage. These compact epithelial domains are characterized by cell-cycle arrest, localized apoptosis, and expression of a conserved suite of signaling molecules including Sonic hedgehog (*Shh*), bone morphogenetic protein 4 (*Bmp4*), fibroblast growth factor 4 (*Fgf4*), and *Fgf20* (11–13). Acting as transient organizers, they coordinate cusp initiation, spacing, and growth, and experimental perturbations demonstrate that even subtle modifications in their activity can dramatically alter crown morphology (14). Whether comparable epithelial organizers operate outside mammals remains unresolved. In squamates (lizards and snakes), regions of reduced proliferation and localized signaling have been described in unicuspid snakes (15–19) and in lizards bearing unicuspid teeth with or without crests (17–20). In multicuspid chameleons, EK-like structures have been proposed (20, 21). However, across these taxa, such domains generally lack the spatial restriction, consistent apoptosis, and tightly defined molecular signatures that characterize the mammalian primary EK, and are therefore typically considered non-homologous. Additional mechanisms, including asymmetric enamel deposition in crest-bearing taxa, further contribute to crown shaping (8, 18). Importantly, the presence of secondary EKs (i.e., organizers of additional cusps) remains largely unexplored in squamates. Evidence remains similarly equivocal in other vertebrate groups. Studies in unicuspid crocodilians have reached conflicting conclusions regarding the presence of EK-like organizers (20, 22, 23), while investigations in fishes have proposed alternative epithelial signaling domains (24) or more discrete EK-like structures resembling mammalian primary and even secondary knots, often without comprehensive assessment of apoptosis or functional validation (3, 25–27). These latter findings have raised the possibility that EK-like epithelial organizers represent a deep vertebrate innovation (28) predating the origin of true teeth (29). Despite decades of investigation, however, the developmental basis of cusp formation outside mammals remains poorly resolved, particularly regarding how epithelial patterning scales across tooth positions, multicuspid dentitions, and evolutionary transitions in crown complexity.

This gap is especially striking in squamates, a group that exhibits remarkable dental diversity closely linked to dietary specialization (9, 10). Multicuspid crowns with two, three, or more cusps have evolved repeatedly and occur in more than half of extant lizard families (10), often accompanied by pronounced anterior-posterior gradients in tooth complexity along the jaw (8, 30). Despite this diversity, only a single multicuspid squamate taxon has been examined in molecular detail (20, 21), and no comparative framework has tested whether convergent crown morphologies arise through shared developmental parameters, altered tissue dynamics, or lineage-specific mechanisms. Here, we address this gap by integrating comparative developmental analyses, quantitative morphometrics, and computational modeling across multiple lizard lineages that independently evolved increased tooth complexity and exhibit strong positional variation along the jaw. We introduce BITES (Bayesian Inference for Tooth Emergence Simulation), a computational framework that couples developmental simulations with phenotype-driven parameter optimization to determine how shifts in developmental landscapes generate observed crown morphologies. Our results reveal that squamate tooth complexity does not depend on a canalized EK system, but instead emerges from a flexible epithelial signaling module in which quantitative modulation of signaling-domain size and proliferative dynamics is sufficient to drive cusp elaboration. This labile patterning system operates both across species and along the jaw axis, consistent with an ancestral epithelial signaling logic that predates the rigid morphological and regulatory constraints characteristic of mammalian dentitions.

## RESULTS

### Phenotypic diversity and convergent evolution of increased tooth complexity

Our previous comparative analyses identified six major clade-level increases in tooth complexity, corresponding to independent gains of multicuspid teeth across both fossil and extant squamates (10). Here, we mapped these events onto a simplified phylogeny restricted to extant clades (Fig. 1A), highlighting their distribution across the corresponding living lineages: Gerrhosauridae, Teiioidea (Teiidae + Gymnophthalmidae), Lacertidae, Chamaeleonidae, Agamidae (excluding Uromastycinae), and Pleurodonta. X-ray computed microtomography scans (microCT-scans) of representative species from each clade (Fig. 1B-G) reveal substantial variation in tooth number, size (from <30 μm to >1.2 mm in width) and crown morphology, ranging from unicuspid to five-cusped forms across species and along the jaw. In contrast to the highly integrated cusp architectures typical of mammals, squamate multicuspid teeth generally exhibit simple transverse cusps arranged laterally across the crown, with little or no precise occlusion. All six species display heterodont dentitions and vary in tooth implantation mode, including pleurodont teeth attached to the medial side of the jaw bone (Fig. 1B-D,G), acrodont teeth attached to the apical margin (Fig. 1E), and mixed acrodont-pleurodont arrangements within a single jaw (Fig. 1F). Across species, tooth complexity consistently increases along the anterior-posterior axis. These positional gradients include transitions from unicuspid to bicuspid teeth (Fig. 1F), to tricuspid teeth (Fig. 1C-E,G), and, in some taxa, to four- or five-cusped crowns (Fig. 1B). Crown morphology varies predictably along these gradients: unicuspid teeth are triangular or conical and may be straight or slightly recurved; bicuspid teeth are typically asymmetrical; tricuspid teeth may be symmetrical or asymmetrical; and tetra- and pentacuspid crowns bear cusps arranged in a fan-shaped row and are generally asymmetrical. Notably, increased tooth complexity is typically accompanied by increased tooth size, suggesting that size-dependent positional scaling underlies variation in cusp number along the jaw and may represent a general mechanism facilitating repeated evolutionary gains in tooth complexity among squamates.

**Figure 1.**
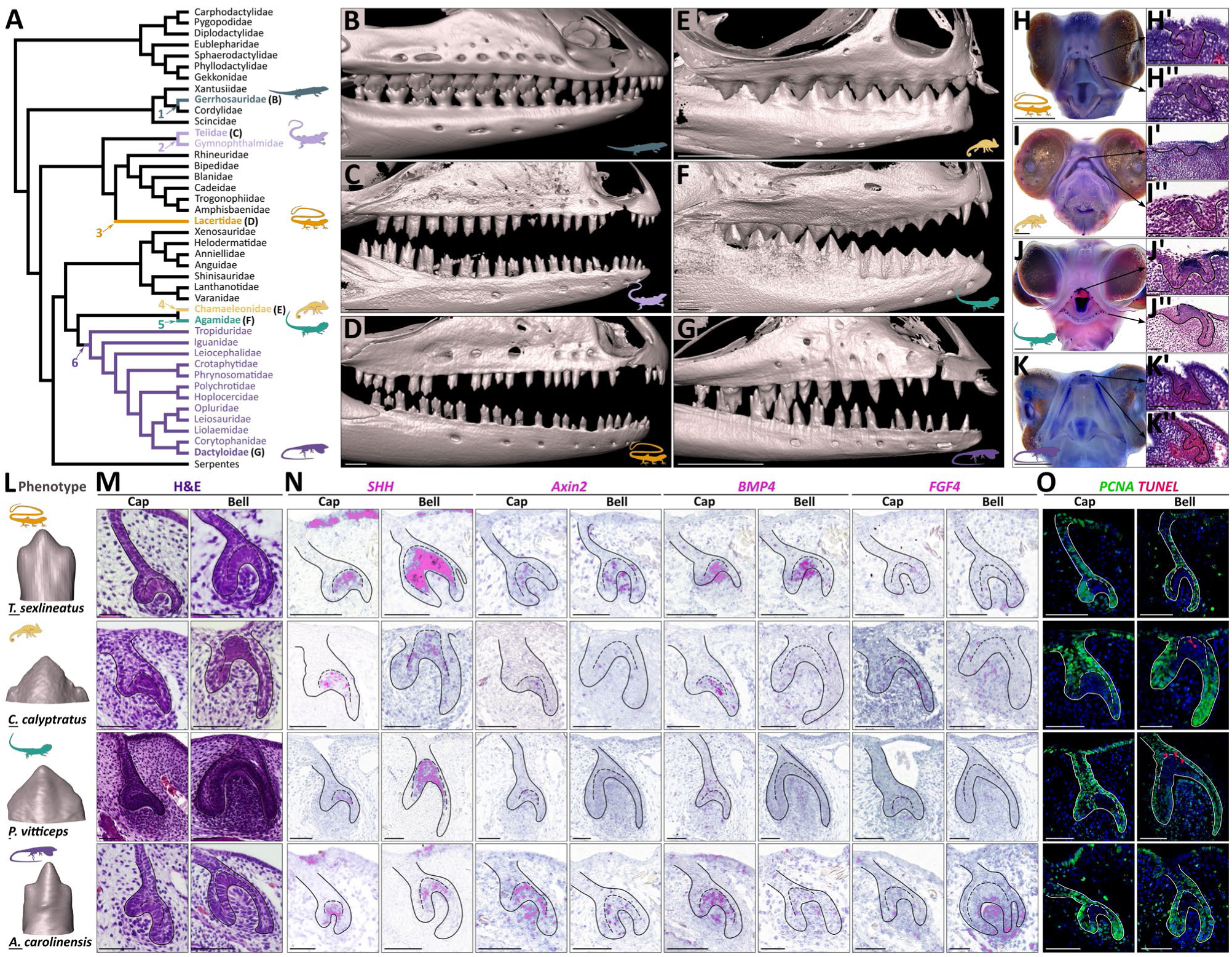
Convergent tooth complexity and epithelial signaling in squamates. (A) Simplified phylogenetic tree of extant lizard families and subfamilies, adapted from (51). Colored branches and associated numbers indicate the six major gains in tooth complexity mapped onto extant lineages. Within Agamidae, this increase excludes Uromastycinae. Silhouettes show representative species included in this study. (B-G) 3D microCT reconstructions of adult skulls in lateral view from representative species exhibiting convergent multicuspid dentitions: *Gerrhosaurus skoogi* (B), *Ameiva Ameiva* (C), *Takydromus sexlineatus* (D), *Chamaeleo calyptratus* (E), *Pogona vitticeps* (F), and *Anolis carolinensis* (G). (H-K) WMISH of embryonic whole heads showing *SHH* expression in the developing early dentition of *T. sexlineatus* (H), *C. calyptratus* (I), *P. vitticeps* (J), and *A. carolinensis* (K). (H’-K’’) Histological sections at the indicated positions within the embryonic upper jaws, confirming tooth bud development. (L) High-magnification 3D renderings of representative adult multicuspid teeth in lateral view from the same four species shown in panels (H-K), with species identities indicated. (M-O) Histological sections stained with H&E (M), ISH expression patterns of conserved EK-associated genes (*SHH*, *Axin2*, *BMP4*, *FGF4*) (N), and double IHC detection of cell proliferation (PCNA) and apoptosis (TUNEL assay) (O) in the same four species at cap and bell stages of tooth development, arranged by horizontal lane corresponding to panel (L). Scale bars: 1 mm (B-K), 50 μm (H’-K’, H’’-K’’, L-O).

### Convergent increases in tooth complexity arise from a variable signaling center

To test whether conserved developmental modules underlie convergent multicuspid tooth morphologies, we examined four representative lizard species with accessible embryonic stages: *Takydromus sexlineatus*, *Chamaeleo calyptratus*, *Pogona vitticeps*, and *Anolis carolinensis* (Fig. 1D-G). Whole-mount in situ hybridization (WMISH) for the early dental marker *SHH* was used to map the timing and spatial pattern of early tooth morphogenesis along the developing jaw (Fig. 1H-K). In squamates, functional teeth are preceded by vestigial null-generation teeth, whose number, position, and extent vary markedly among species (30). In some species, vestigial teeth extend posteriorly before any functional teeth emerge (Fig. 1H-J’’), whereas in others they are restricted to one or two teeth at the extreme anterior tip (Fig. 1K-K’’). After aligning comparable embryonic stages across species, we identified tooth-specific cap and bell stages histologically (Fig. 1L,M), corresponding to the developmental window during which cusp-patterning centers are expected to arise (30, 31). Because squamate tooth germs initiate differentiation earlier than those of mammals, stage assignment relied on tooth morphology and epithelial maturation rather than discrete morphological boundaries. Mammalian EKs serve as a conceptual reference: compact, nonproliferative epithelial clusters expressing conserved signaling molecules, precisely specifying cusp position and spacing before being eliminated by apoptosis. Across all four species, histology reveals a localized thickening of the inner enamel epithelium (IEE) at the cap stage (Fig. 1M). This domain expresses canonical EK-associated markers of Hedgehog (*SHH*), Wnt (*AXIN2*, *WNT10b, WNT6*), BMP (*BMP4*, *BMP2*), and FGF (*FGF4*) signaling (Fig. 1N and Fig. S1). However, unlike the sharply delimited mammalian primary EK, the squamate signaling domain is consistently broader and less spatially restricted, with marker expression extending into adjacent regions of the enamel organ (Fig. 1N and Fig. S1). Three-dimensional (3D) reconstructions of tooth germs further reveal that *SHH* forms a continuous ring-like domain rather than discrete cusp-specific centers (Fig. S2). This epithelial domain persists into the late bell stage and remains active during epithelial folding associated with additional cusp formation and throughout early mineralization (Fig. S2). Consistent with this organization, no secondary EK-like organizers were detected (Fig. 1M,N), and canonical secondary EK markers such as *FGF4* and *FGF20* were absent (Fig. 1N and Fig. S1). Cellular behavior further distinguishes the squamate domain from the mammalian EK. Immunohistochemistry (IHC) for the proliferation marker proliferating cell nuclear antigen (PCNA) reveals a broad and variably defined zone of reduced proliferation within the IEE that persists during cusp addition (Fig. 1O), rather than a sharply bounded proliferation arrest. Similarly, apoptosis is weak or absent in most species based on TUNEL staining, with only the two acrodont taxa (*C. calyptratus* and *P. vitticeps*) showing sparse TUNEL-positive cells at the bell stage (Fig. 1O), indicating that cusp morphogenesis in squamates is not associated with extensive apoptotic elimination of epithelial cells. Together, the spatial breadth, persistence, and interspecific variability of molecular expression, proliferation, and apoptosis demonstrate that this epithelial signaling domain differs fundamentally from the stereotyped mammalian EK system. Rather than acting as a discrete, temporally resolved organizer, it functions as a labile growth-and-patterning field whose properties vary among species. Despite this variability, all species examined reliably generate distinct cusps, indicating that this domain represents a conserved yet highly adaptable organizer of crown morphology rather than a strict homolog of mammalian primary or secondary EKs.

### Positional scaling and dysregulated signaling underlie altered tooth complexity in *P. vitticeps*

To determine whether the identified labile signaling domain regulates cusp patterning, we analyzed tooth development in wild-type (WT) *P. vitticeps* and in spontaneous scaleless mutants carrying a mutation in the ectodysplasin (EDA) pathway (32). These mutants exhibit positional transformations in anterior pleurodont teeth associated with *EDA*-dependent modulation of tooth size (33). Here, we focused on the posterior acrodont dentition, whose crown phenotype has not been characterized. In WT animals, as in other multicuspid lizards (Fig. 1B-G), lateral skull views and microCT analyses reveal a progressive increase in crown complexity along the anterior-posterior axis of both mandible and maxilla, including the addition of a lateral cusp at posterior positions (Fig. 2A-C). This gradient is accompanied by a strong positive correlation between tooth shape and tooth width, with crowns becoming progressively larger and more complex toward the posterior region of the jaw (Fig. 2D). In *EDA* mutants, this positional relationship is disrupted. Most mutant teeth are significantly larger and correspondingly more complex than WT teeth at equivalent jaw positions (Fig. 2B-D and Table S1). As a result, many mid-jaw mutant teeth occupy regions of morphospace normally restricted to posterior WT teeth, with some extending beyond the WT range, effectively shifting the size-complexity trajectory forward along the jaw (Fig. 2C,D). In addition, mutant acrodont teeth display pronounced shape abnormalities, including bifid (two-lobed) central cusps or duplication of the cusp apex at positions where WT crowns are uniformly simple, producing geometries outside the WT range (Fig. 2B,D). These phenotypes therefore provide a natural perturbation for dissecting the mechanisms controlling tooth complexity. To identify the molecular basis of these defects, we analyzed transcriptome profiles from early acrodont dental epithelia of WT and *EDA* mutant embryos. This revealed extensive molecular disruption, with nearly half of the differentially expressed genes associated with developmental process Gene Ontology (GO) categories and approximately one quarter corresponding to developmental genes involved in the regulation of cell proliferation (Fig. 2E and Table S2). Among these, nine keystone regulators of mammalian tooth development (34) were significantly dysregulated. Notably, the EK marker *SHH* was the most strongly upregulated developmental and proliferation-associated gene in mutants (Fig. 2E and Table S2). This pattern parallels previously reported increases in epithelial proliferation in mutant posterior dentary tooth germs, including at the cap stage (33), indicating that disruption of the *EDA* pathway alters both epithelial signaling activity and growth dynamics. Together, these results show that aberrant crown morphologies in *EDA* mutants arise from quantitative dysregulation of the developing dental epithelium, supporting a model in which positional scaling and epithelial growth jointly regulate tooth complexity in squamates.

**Figure 2.**
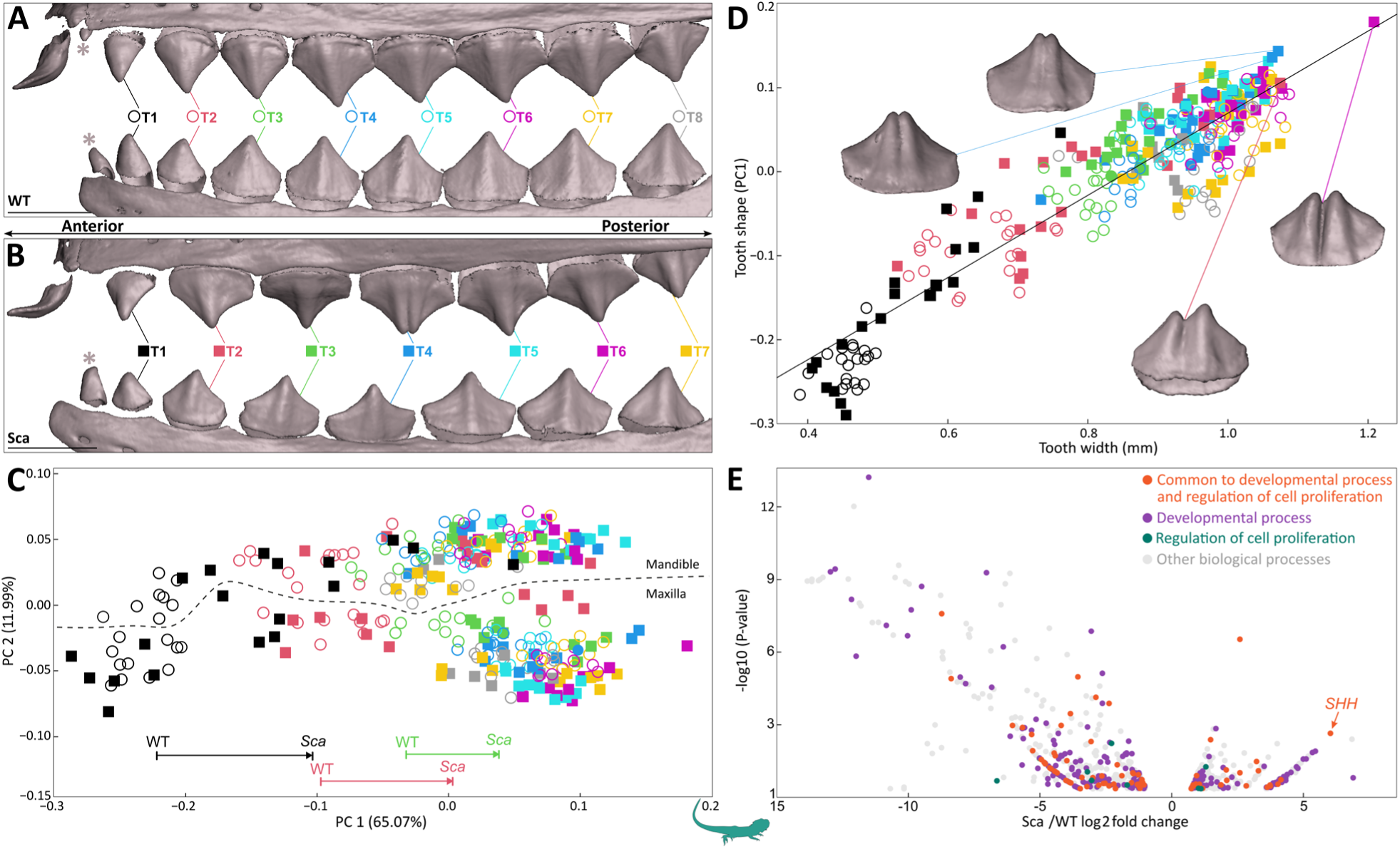
Altered tooth size and shape in mutant scaleless *P. vitticeps*. (A,B) 3D-rendered upper jaw (premaxilla and maxilla; top row) and mandible (bottom row) of WT (A) and homozygous scaleless (Sca) (B) *P. vitticeps* lizards at hatchling stage in lateral view. Individual acrodont teeth are numbered sequentially from anterior to posterior (T1-T8). Teeth are color-coded according to their position along the jaw in WT (colored open circles) and Sca mutants (colored squares). Grey asterisks indicate pleurodont teeth not included in the analyses. Scale bars: 1 mm. (C) Tooth-shape morphospace showing the distribution of all mandibular and maxillary acrodont teeth from WT and Sca *P. vitticeps* at hatchling stage (n = 5 animals per group). Symbols and colors correspond to tooth positions as indicated in panel (A). Numbers in parentheses indicate the percentage of total shape variance explained by each principal component (PC) axis. Arrows at the bottom of the graph indicate a shift of the mean values of Sca anterior teeth along PC1 in morphospace compared to WT. (D) Scatterplot showing the relationship between tooth shape, represented by PC1, and tooth width in WT and Sca *P. vitticeps* at hatchling stage (n = 5 animals per group). The regression indicates a strong correlation between tooth size and shape along the jaw. Symbols and colors correspond to tooth positions as indicated in panel (A). 3D renderings of representative *Sca* mutant bifid teeth and their positions in morphospace are shown. (E) Volcano plot of genes significantly differentially expressed (FDR-corrected *P* < 0.05) between WT and Sca dental epithelium from early acrodont tooth germs (n = 3 per group). The x-axis shows log₂ fold change (Sca relative to WT), and the y-axis shows associated *P*-values. Positive log₂ values indicate upregulation in *Sca* teeth. *SHH* is among the top upregulated genes in mutant teeth. Colors indicate selected Gene Ontology (GO) biological process categories, shown individually or in combination, for all differentially expressed genes.

### Positional modulation of epithelial signaling and growth within *A. carolinensis* dentition

To determine whether positional modulation of the squamate signaling domain operates within naturally graded heterodont dentitions, we performed high-resolution developmental and quantitative analyses along the anterior-posterior jaw axis of *A. carolinensis*. This species exhibits a shift from anterior unicuspid to wider posterior tricuspid teeth separated by a variable intermediate zone comprising both morphotypes (Fig. 3A and Fig. S4), where tooth complexity remains tightly correlated with crown width (Fig. 3B, Fig. S5 and Table S3). Direct comparison of simple and complex teeth required first establishing the embryonic sequence of tooth formation at each jaw position, as timing and positional patterning vary widely among squamates and have not been characterized in pleurodont species (see Fig. 1H-K). MicroCT and histological analyses show that the first functional teeth initiate in the embryonic jaw at a position that later becomes relatively more posterior as the jaw elongates anteriorly. Because cusp morphology corresponds to final jaw position, some of these early-forming teeth ultimately occupy the transitional or tricuspid region of the tooth row (Fig. 3A,B and Fig. S4). Subsequent teeth are added anterior to this initial position while additional posterior teeth continue to form, resulting in bidirectional expansion of the tooth row (Fig. 3A and Fig. S4). To investigate the developmental basis of the complexity gradient, we generated 3D reconstructions of cap-stage tooth germs stained by ISH for the EK marker *SHH*, whose signaling-center expression and mutant alteration are both shown in this study. Volumetric quantification of *SHH* expression normalized to tooth-germ volume reveals that anterior unicuspid and posterior tricuspid teeth exhibit similarly sized *SHH* domains at the early cap stage (Fig. 3C and Table S4). At the late cap stage, however, posterior teeth display a significantly enlarged normalized *SHH* domains, driven by lateral expansion of *SHH* expression (Fig. 3D and Table S4). Notably, IEE cells undergo changes in shape and orientation as they differentiate into columnar, polarized ameloblasts; however, cell size does not differ significantly between early and late cap stages (Fig. 3E and Table S5). Thus, rather than scaling in direct proportion to overall tooth size, the squamate signaling domain expands disproportionately at posterior positions that later develop increased cusp complexity. These findings indicate that growth and patterning are coupled nonlinearly, such that enlargement of the tooth germ is accompanied by allometric expansion of the signaling domain, potentially enabling the formation of additional cusps. In contrast to the strict scaling relationships described in mammals (14, 35), this more flexible relationship may underlie the pronounced positional variation in tooth complexity along the squamate jaw.

**Figure 3.**
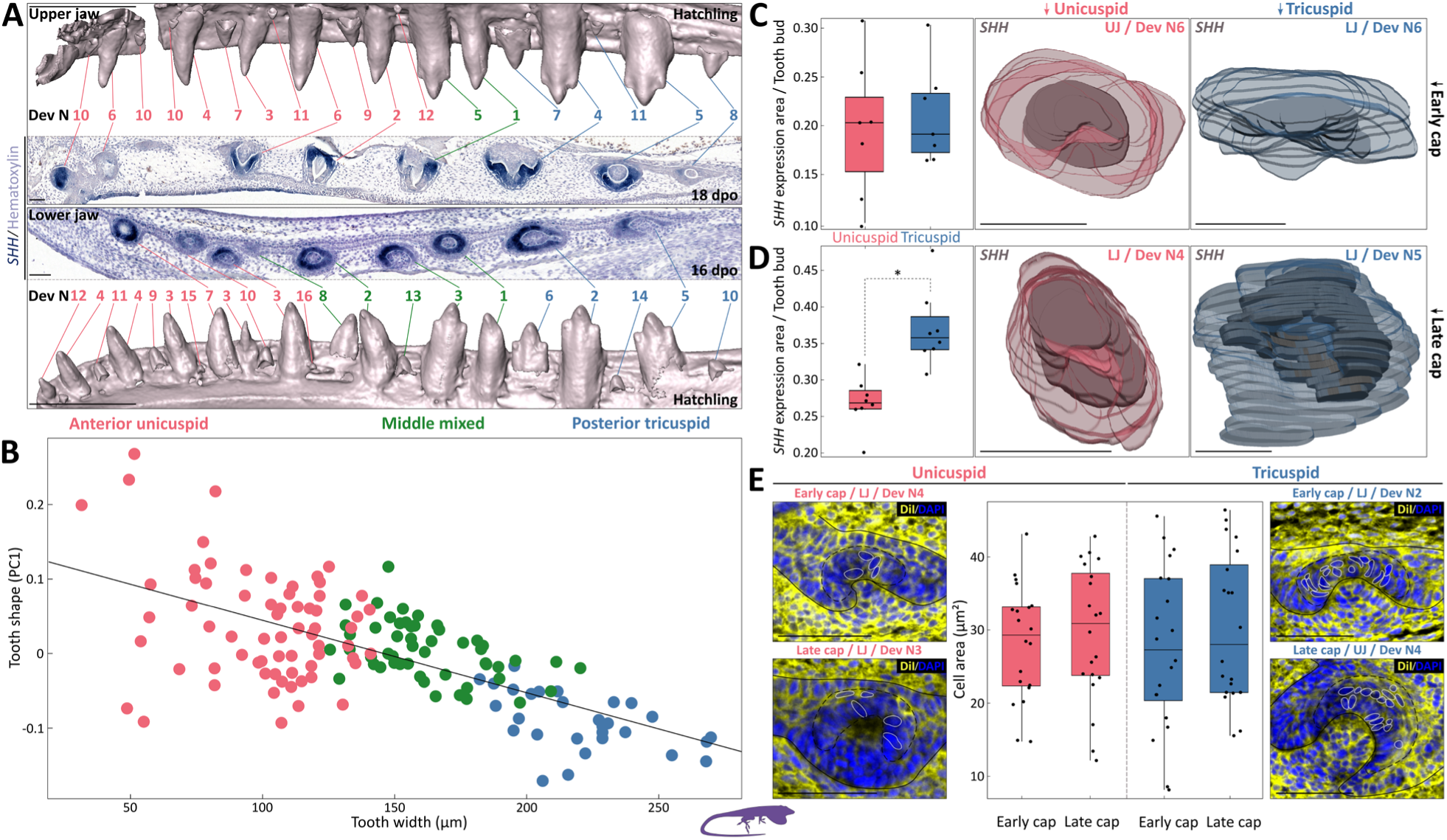
Positional scaling of tooth complexity in *A. carolinensis*. (A) Representative 3D renderings of upper (top) and lower (bottom) hatchling jaws in lateral view, together with and ISH (*SHH*) and hematoxylin-stained sagittal histological section of the upper embryonic jaw at 18 days postoviposition (dpo) (top) and lower embryonic jaw at 16 dpo (bottom). Histological sections were selected to show the maximum number of developing teeth in a single section and thereby illustrate dentition patterning during tooth row development. Developmental number (Dev N), indicating the order of tooth initiation, is conserved across jaw regions and individuals and is shown for all visible teeth. Corresponding tooth positions between embryonic and hatchling stages are connected by lines. Colors indicate tooth complexity, from anterior unicuspid teeth (brick red) to posterior tricuspid teeth (blue), with a variable intermediate zone (dark green). (B) Scatterplot showing the relationship between mandibular tooth shape (PC1) and width at the hatchling stage (n = 6 animals per group). The regression indicates a strong correlation between tooth size and shape along the jaw. Colors correspond to tooth positions in (A). (C,D) 3D reconstructions and quantification of *SHH* ISH signal volume normalized to tooth germ volume in anterior unicuspid and posterior tricuspid teeth at the early cap (C) and late cap (D) stages. Dental epithelium is colored by tooth positions as in (A), and *SHH* signal is shown in gray. Box plots show normalized *SHH* signal volumes (n = 7 teeth per group at early cap and n = 8 at late cap), with a significant difference at late cap (randomization test, *P* < 0.05). (E) Quantification of cell size and cell number overlapping the *SHH* signal. Representative sagittal histological sections of unicuspid and tricuspid tooth buds at early and late cap stages are shown. Measured cells are outlined with a solid white line, and the approximate signaling center area is indicated by a dashed black line. Box plots show measured cell areas (µm²) (n = 20 cells for each stage and tooth position). Scale bars: 1 mm (CT scans) and 50 µm (histological sections) in (A); 50 µm in (C-E).

### *In silico* modeling unifies developmental and evolutionary drivers of tooth shape

Computational models of tooth morphogenesis were initially developed to explore how developmental parameters generate phenotypic diversity in mammals (14, 36), with later extensions to other clades such as sharks (29, 37). These approaches typically adjust model parameters manually to achieve a desired tooth phenotype. However, empirical studies usually start with observed morphologies rather than known developmental parameters, making forward parameter tuning impractical for species with uncharacterized tooth shapes or divergent signaling dynamics. To overcome this limitation, we developed BITES (Fig. 4A and Fig. S6), an inverse-modeling framework in which real tooth shapes are specified first and matched to simulated outputs through automated Bayesian optimization built on a morphodynamic tooth model (36). Applying BITES to our squamate dataset successfully recapitulated all six independent increases in tooth complexity, matching crown morphologies in the four experimentally examined species as well as in two additional taxa for which embryonic material was unavailable (Fig. 4B-E and Table S6). Although squamate tooth germs lack discrete mammalian-like EKs, the broader and more labile epithelial signaling domains identified experimentally map onto the activator-inhibitor and growth parameters recovered by the model. Across independent optimization runs, similar crown morphologies could be achieved through multiple parameter combinations; however, only a limited subset of parameters consistently showed low variance across replicates and species (Fig. 4C,D, Fig. S7 and Table S7). The least variable among these span both cellular processes, such as epithelial growth rate, and genetic-patterning components, particularly inhibitor strength corresponding to *SHH*-related signaling in the model, indicating that convergent multicuspid tooth shapes occupy a restricted region of developmental parameter space. Strikingly, the same parameters vary systematically *in vivo*, differing between unicuspid and multicuspid teeth along the jaw as well as between normal and malformed crowns in *EDA*-mutant *P. vitticeps*, thereby supporting their central role in cusp patterning. Further simulations incorporating graded shifts in these parameters recapitulate the anterior-posterior complexity gradient observed in *A. carolinensis*, providing *in silico* validation of the developmental trends inferred experimentally (Fig. 4E). Together, these results show that both interspecific convergence and intraspecific positional patterning in squamate dentitions arise from quantitative modulation of a limited number of key developmental parameters acting on a flexible epithelial field, thereby directly linking developmental dynamics to evolutionary diversification.

**Figure 4.**
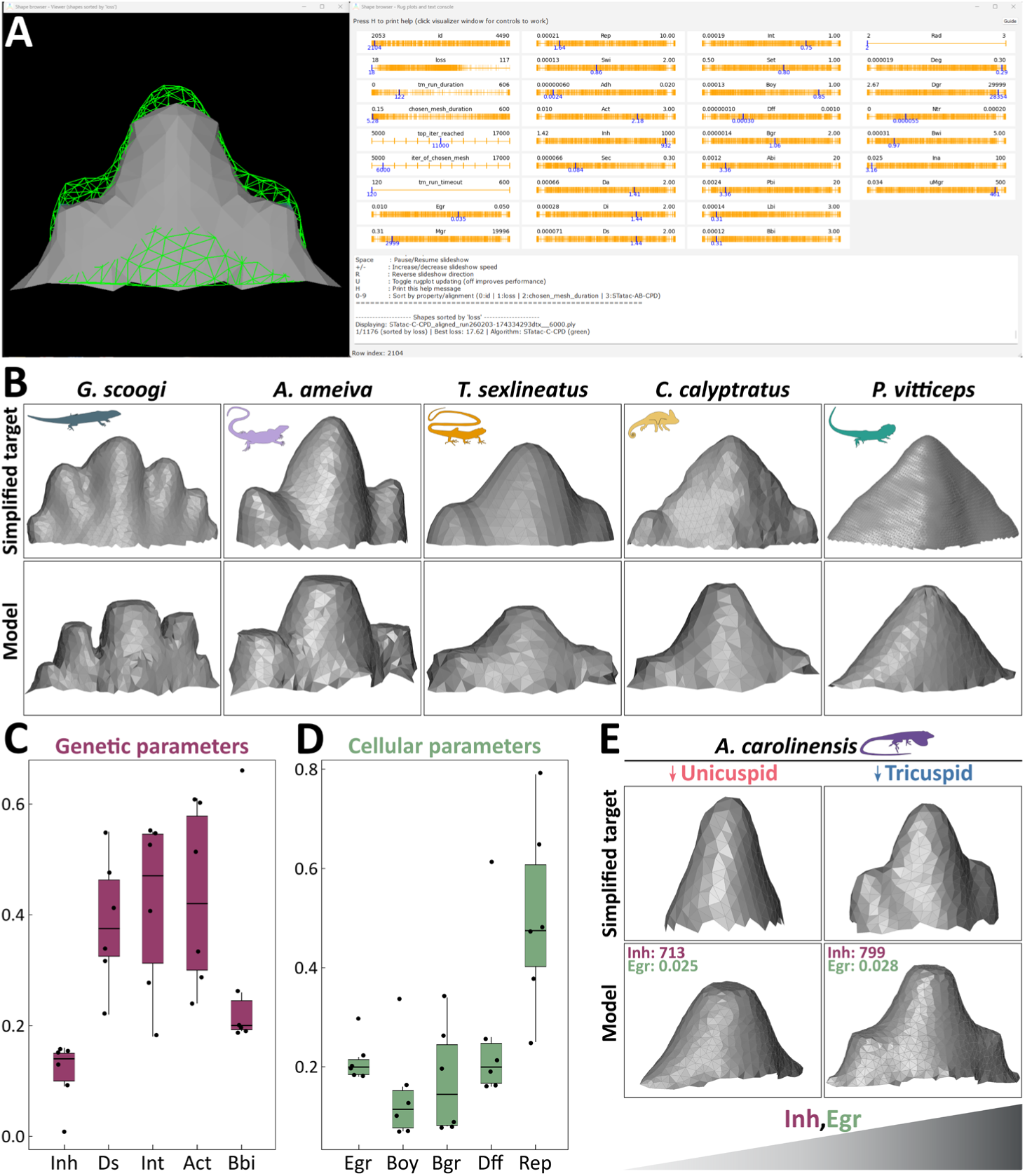
*In silico* modeling of tooth shape convergence using BITES. (A) Overview of the BITES interface showing an interactive 3D visualization of the optimized simulated tooth shape superimposed on the target source shape (green wireframe) in the left window, and rug plots illustrating the distribution of tested parameter values across their search ranges in the right window. Highlighted values in each rug plot correspond to the parameter set used to generate the shape currently displayed. Genetic and cellular parameters correspond to those defined in the original morphodynamic tooth model (36). Data are shown for a single optimization run targeting posterior tricuspid tooth shapes from *A. carolinensis*. (B) Example *in silico* modeling results for five lizard species (species indicated): the top row shows simplified target tooth meshes, and the bottom row shows examples of the best-fitting models. Model parameters are provided in Table S6. (C,D) Box plots showing variability in normalized *in silico* model parameters across three independent optimization runs for complex tooth shapes in each of the six convergent species. The five least variable genetic parameters are shown in (C) and cellular parameters in (D). Variability was calculated as the range of normalized parameter values. Box plots for all parameters are provided in Fig. S7. The least variable constrained parameters across species include epithelial growth rate (Egr) and inhibitor strength (Inh), corresponding to *SHH*-related signaling in the model. (E) BITES optimization modeling results from unicuspid (left panels) to tricuspid teeth (right panels) in *A. carolinensis*. Simplified target tooth meshes and representative best-fitting models aligned with the target meshes are shown for each tooth. Modulating Egr and Inh alone was sufficient to transform unicuspid into tricuspid tooth shapes.

## DISCUSSION

Our comparative, mechanistic, and modeling analyses reveal a developmental route to squamate tooth-shape diversification that diverges sharply from the canonical mammalian model. Instead of relying on the stepwise deployment of discrete EKs, convergent increases in cusp number across lizards arise from a single, flexible epithelial signaling region whose size, dynamics, and proliferative landscape vary across species and along the jaw. This labile organizer retains core elements of EK-associated molecular logic yet lacks the strict spatial boundaries as well as functional and structural constraints that characterize mammalian systems. Together, these findings demonstrate that cusp elaboration outside mammals can emerge from a diffuse and adaptable developmental module capable of both evolutionary and positional tuning, likely reflecting an ancestral mode of tooth-shape regulation in vertebrates.

### A flexible signaling center drives squamate tooth morphogenesis

A key finding of this study is that squamate teeth do not employ the canonical mammalian EK system. Across all examined convergent lineages, we identified a single epithelial signaling domain persisting from cap to bell stages, marked by conserved expression of Hedgehog, Wnt, BMP, and FGF pathway genes (11, 12). Yet this domain lacks the defining features of mammalian EKs: it is spatially broad rather than compact, not simply scaled with tooth size, exhibits a wide but variable zone of reduced proliferation, and never resolves into secondary organizers at prospective cusp positions. Instead of functioning as a discrete morphological module, the squamate signaling domain operates as a persistent growth-and-patterning field whose size and morphology can shift across species and even across tooth positions along the jaw. This organization is developmentally coherent in a system where cusp formation emerges from spatial gradient in epithelial signaling and proliferation, rather than from the sequential appearance of restricted signaling centers. Notably, low or absent apoptosis in squamate tooth germs at the bell stage is consistent with such prolonged signaling-field activity, whereas in mammals apoptosis marks the termination of EK activity (38), highlighting a fundamental difference in how signaling domains are regulated across amniotes. The epithelial growth differentials are likely to drive the bending and folding events that initiate cusp shape, providing a mechanistic basis for folding behaviours previously documented in multicuspid squamate odontogenesis (8). These results argue against homology with the mammalian primary EK as a structurally and functionally bounded organizer. Nonetheless, the presence of conserved signaling signatures and localized proliferation arrest indicates that squamates retain ancestral EK-like developmental logic deployed in a more diffuse and dynamic manner. This distinction clarifies long-standing uncertainty about reptilian cusp initiation (e.g., (16, 17, 19–21) and aligns with observations from other vertebrates such as sharks, where broader epithelial signaling territories do not resolve into a compact mammalian-like EK (24). Although interpretations differ among fish systems (24–29, 39), available evidence indicates that classical, sharply bounded, apoptotic EKs are not a universal vertebrate feature. Our results therefore support the view that a diffuse epithelial organizer may represent the ancestral vertebrate condition, with the mammalian EK system emerging later through spatial compartmentalization and tighter coupling between signaling and growth arrest. Alternatively, compact organizers may have arisen earlier in gnathostome evolution and were later relaxed in reptiles, or evolved independently in different lineages. Distinguishing between these scenarios will require broader comparative sampling, but both highlight that mammals occupy a uniquely canalized region of the vertebrate tooth-patterning landscape, potentially reflecting the increased evolutionary pressure for precise cusp spacing, occlusion, and tribosphenic mastication (11, 40).

### Epithelial field dynamics drive squamate tooth complexity and ecological diversification

The discovery of a broad, flexible signaling domain has major implications for understanding how squamates generate large morphological transitions. Across species, cusp number correlates strongly with tooth size, and our detailed analyses in *A. carolinensis* show that positional differences in crown complexity are accompanied by changes in the spatial extent of the signaling domain. Teeth at different jaw positions initially form signaling domains of comparable size, but later diverge through disproportionate expansion of *SHH*-positive epithelium in regions that later give rise to additional cusps. This nonlinear relative scaling contrasts with mammalian systems, in which EK size scales predictably with tooth size and cusp formation is constrained by absolute tooth-size thresholds (14, 35). Squamates deviate markedly from this pattern: although squamate teeth are generally smaller than mammalian teeth, very small teeth can be multicuspid, larger teeth may remain unicuspid, and many species exhibit gradients in cusp number at sizes below the mammalian threshold. Together, these patterns suggest that squamates do not rely on discretized secondary EKs or strict size thresholds to generate crown complexity. Instead, cusp formation arises within a broad, disproportionately expanding epithelial signaling field that permits additional cusps to form. The evolutionary potential of this system is illustrated by *EDA*-mutant *P. vitticeps*, in which dysregulated epithelial proliferation produce an unstable phenotype that frequently results in enlarged teeth with cusp duplications at atypical positions, disrupting the normal positional gradient of size and complexity. Together with our BITES modeling results, which recapitulate both interspecific convergence and intraspecific gradients through modulation of a limited set of constrained parameters, including epithelial growth rate and activator-inhibitor strength, these findings show that the squamate signaling center functions as an integrative regulator. In this system, relatively small quantitative changes generate disproportionately large shifts in crown morphology. This intrinsic developmental plasticity likely underlies the pronounced positional heterodonty characteristic of a large proportion of lizard jaws, in which tooth size and complexity increase along the jaw but do so with local variation, such that adjacent teeth can differ markedly in shape and crowns rarely converge on a single optimal form. In lizards, tooth patterning generally operates as a continuous, position-specific process, permitting both ontogenetic and evolutionary flexibility and generating variability consistent with continuous tooth replacement in many lineages and weak occlusal constraints across squamates. By contrast, mammalian dentitions are typically organized into discrete heterodont tooth classes, within which teeth approximate functionally optimized morphologies shaped by precise occlusion and strong spatial constraints (41), reflecting a more canalized patterning mode. These contrasting developmental architectures likely contributed to divergent evolutionary trajectories. Squamates show repeated evolutionary gains and losses of multicuspidity linked to dietary diversification, reflecting a non-unidirectional pattern of tooth evolution (10) consistent with a flexible, permissive patterning field that can be easily tuned across lineages. In mammals, canalized tooth patterning promoted functional specialization around precise occlusal performance, facilitating evolutionary radiations by preserving tooth function across body-size evolution (35, 42) and dietary specialization (43–46), while constraining departures from established occlusal optima. Together, our results redefine how tooth complexity can arise outside mammals, demonstrating that cusp elaboration does not require discretized EKs but can instead emerge from modulation of a broad epithelial signaling field. This flexible developmental architecture links positional heterodonty, evolutionary convergence, and ecological diversification within a single mechanistic framework. More broadly, our findings highlight how conserved signaling logic can be deployed through distinct patterning architectures across vertebrates, generating alternative evolutionary solutions to shared functional demands.

## MATERIALS AND METHODS

### Sample collection

All embryonic and postnatal stages of bearded dragon (*P. vitticeps*), green anole (*A. carolinensis*), and long-tailed grass lizard (*T. sexlineatus*) were obtained from reptile colonies at the University of Helsinki. Veiled chameleon (*C. calyptratus*) embryos were obtained from private breeders. For embryonic stages, fertilized eggs were incubated on a moistened vermiculite substrate at 29.5°C (*P. vitticeps*, *A. carolinensis*, *T. sexlineatus*) or 26.5°C (*C. calyptratus*). Embryos were collected at regular days postoviposition (dpo) to obtain developmental stages of interest and were staged based on external morphology using published developmental tables for each species (47–50). Homozygous *EDA*-mutant *P. vitticeps* lizards were obtained by crossing heterozygous mutant males with wild-type females to bypass the reduced fertility of adult mutants and increase the yield of mutant embryos (33). All genotyping was performed by polymerase chain reaction using genomic DNA extracted from tail tissue, with primers as previously described (32). Data on additional lizard species (*Ameiva Ameiva* and *Gerrhosaurus skoogi*) were obtained from publicly available datasets hosted on MorphoSource (www.MorphoSource.org) or from specialized retailers. Phylogenetic relationships used for comparative analyses follow the squamate phylogeny of (51). All reptile captive breedings and experiments were done following international standards and were approved by the Laboratory Animal Centre of the University of Helsinki and/or the National Animal Experiment Board in Finland (license numbers ESAVI/7484/04.10.07/2016, and ESAVI/13139/04.10.05/2017, and ESAVI/5416/2021).

### CT-scanning and 3D volume rendering

High-resolution microCT scans of *P. vitticeps*, *A. carolinensis*, *T. sexlineatus*, *C. calyptratus*, and *A. Ameiva* specimens were performed at the University of Helsinki using Scyscan 1272 instrument (Bruker, Belgium). Samples were initially fixed overnight or longer, depending on the sample size, in 4% paraformaldehyde (PFA) at 4°C and subsequently dehydrated through an increasing alcohol series. Scanning parameters followed previously published protocols (31, 48), with minor adjustments depending on specimen age and the specific aims of each scan. All scans were reconstructed using NRecon 1.7.0.4 software (Bruker). 3D volume rendering and manual segmentation were done using Amira 5.5.0 (Thermo Fisher Scientific, USA). CT data for an adult *G. skoogi* specimen were obtained from the MorphoSource database.

### 3D surface analyses and geometric morphometrics of tooth shape

Tooth shape variation within species was quantified using a landmark-based geometric morphometric approach combining manually placed fixed landmarks and automatically projected surface semi-landmarks (Fig. S8). Fixed landmarks were placed using 3D Slicer v5.6.1 (52), and surface semi-landmarks were generated using the SlicerMorph extension (53). The landmark configuration for *P. vitticeps* consisted of five fixed landmarks and 230 semi-landmarks, whereas that for *A. carolinensis* comprised seven fixed landmarks and 138 semi-landmarks. For *P. vitticeps*, fixed landmarks were placed as follows: (1) tip of the main cusp; (2) widest point of the crown on the posterior side; (3) widest point of the crown on the anterior side; (4) midpoint between landmarks 2 and 3 on the buccal side; and (5) midpoint between landmarks 2 and 3 on the lingual side. For *A. carolinensis*, landmarks were placed at (1) the tip of the main cusp; (2) the tip of the posterior accessory cusp; (3) the tip of the anterior accessory cusp; (4 and 5) the junction between the tooth crown and jawbone on the buccal side; and (6 and 7) corresponding positions on the lingual side. All morphometric analyses were performed in R v4.3.2 (54). A single semi-landmark template was created and projected onto all tooth surfaces, after which semi-landmarks were slid to minimize bending energy (55). Template creation, semi-landmark projection, and sliding were conducted using the Morpho package v2.13 (56). A Generalized Procrustes Analysis (GPA) was applied to remove the effects of scale, position, and orientation, followed by a Principal Component Analysis (PCA) to visualize patterns of tooth shape variation within species. GPA and PCA were performed using the geomorph package v4.0.10 (57). To assess tooth size, the maximum basal tooth width at the tooth-jaw interface was measured for each individual tooth in labial view using the measurement tools in Amira v5.5.0 (Thermo Fisher Scientific, USA). Statistical significance was calculated using independent Student’s t test, and significance was determined at *P* value <0.05.

### Histology, immunohistochemistry (IHC), and apoptosis detection on paraffin sections

Following dissection, tissues or embryos were fixed overnight in 4% PFA at 4 °C. Embryos at later developmental stages exhibiting mineralization were decalcified in EDTA solution (0.5 M, pH 7.5) for 1-12 weeks, depending on sample size. Fixed and decalcified tissues were gradually dehydrated to 100% methanol, embedded in paraffin blocks using a Leica ASP200 embedding system, and sectioned at 5 or 7 μm with a Microm HM355 microtome. Hematoxylin and eosin (H&E) staining of dental sections was performed according to standard protocols. Immunofluorescent IHC staining was carried out as previously described (32), with overnight incubation at 4 °C using primary antibodies known to recognize reptile and/or chicken epitopes: proliferating cell nuclear antigen (PCNA; 1:300, mouse monoclonal, BioLegend, cat# 307901, RRID:AB_314691). Sections were subsequently incubated with Alexa Fluor–conjugated secondary antibodies (Alexa Fluor 488: goat anti-rat IgG, Thermo Fisher Scientific, cat# A-11006, RRID:AB_2354074) for 1 h at room temperature (RT) or overnight at 4 °C, and mounted with Fluoroshield mounting medium (Sigma-Aldrich) containing 4′,6-diamidino-2-phenylindole (DAPI). Nuclear DNA fragmentation in apoptotic cells was detected in situ using a TUNEL assay (In Situ Cell Death Detection Kit, Roche) according to the manufacturer’s instructions. Prior to TUNEL staining, tissues were treated with proteinase K (10 mg/ml, 15 min at RT). Positive controls were generated by DNase treatment before staining. Images of stained sections were acquired either using 3DHISTECH Pannoramic 250 FLASH III digital slide scanner, and processed in SlideViewer v2.6 (3DHISTECH, Hungary), or using a Leica DM5000B widefield microscope and Leica Application Suite X (LAS X) 3.4.2 12.4.18 software and processed in Adobe Photoshop CC using levels adjustment.

### DiI staining and quantitative analysis of cell size

DiI staining was performed on tissue sections prepared as described above. After rehydration, slides were incubated with DiI (Biotium cat# 60010) solution (∼125 μg/ml) for 1 h at RT. The slides were then washed and mounted with Fluoroshield mounting medium (Sigma-Aldrich) containing DAPI. Images were acquired using Olympus VS200 digital slide scanner, and cell perimeters were measured in OLYMPUS OlyVIA v4.2. Cell perimeters were compared using the randomized test, with significance defined as *P* value <0.05. Statistical analysis was performed in R v4.3.2.

### In situ hybridization (ISH)

ISH on paraffin sections and whole-mount ISH (WMISH) were performed as described previously (31, 58, 59). Species-specific digoxigenin (DIG)-labeled antisense riboprobes were generated from coding sequence (CDS) regions of selected genes, based on publicly available lizard genome assemblies (60–65). For *T. sexlineatus: SHH* (793 bp; forward primer (fp), 5’-AAGAGCTCACCCCCAATTACAA-3’; reverse primer (rp), 5’-RTTGATGAGGATRGTGCCCTGAG-3’) (30), *Axin2* (718 bp; fp, 5’-ACTTCTGGTTTGCCTGCAATG-3’; rp, 5’-AGAAACTTGACCGTTGGCCTT-3’), *BMP4* (850 bp; fp, 5’-AGACACCATGACGCCTGGTAA-3’; rp, 5’-CTTGAGGTAAAGAGCGGCTAATCC-3’), *FGF4* (620 bp; fp, 5’-TATTTTTAGAGCMATCGCCTCCCTG-3’; rp, 5’- TGCCTTTCTTTGTTCGTCCGT-3’). For *P. vitticeps: SHH* (931 bp) (32), *FGF20* (592 bp; fp, 5’-ATGGCTCCCTTAGCCGAA-3’; rp, 5’-GCCKGGGCAAGAAATGRGTG-3’), *Wnt10b* (547 bp; fp, 5’-TGCTTTCACACTCTCGTTGC-3’; rp, 5’-TTTGGCCAGACGGTTTCTGT-3’). For *A. carolinensis: Axin2* (936 bp; fp, 5’-GCCTGCCAGTTTGTTTCTTGT-3’; rp, 5’-TGACCATTGGCCTTGACACT-3’), *BMP2* (773 bp; fp, 5’-GAAGCCGAGCGGAGACTGA-3’; rp, 5’-TGATGCACTAATCGGGTGT-3’), *BMP4* (761 bp; fp, 5’-CTCCCTTTGATCCGCGTTGG-3’; rp, 5’-GTTGATCCGGTGGAGGTCAC-3’), *FGF4* (748 bp; fp, 5’-CCGCCCGGGGTGTATTTTTA-3’; rp, 5’-GTGGGAGAGACTTTGTTGCC-3’), *Wnt6* (884 bp; fp, 5’-TGYCAGACRGAGCCTGAGAT-3’; rp, 5’-CYTTGCGYACVGTGCACTTC-3’). Prior to riboprobe synthesis, plasmids containing each insert were linearized and sequenced to confirm insert identity and orientation. Riboprobes synthesized from *P. vitticeps* and *A. carolinensis* cDNA were used for ISH in all three iguanian species (*P. vitticeps, C. calyptratus, and A. carolinensis*). ISH was performed using a hybridization temperature of 63-65°C. Following hybridization, tissues were washed and blocked from non-specific antibody binding with blocking solution (2% Roche blocking reagent and 5% sheep serum) before incubating overnight at 4°C (or 1 hour at RT) with anti-DIG antibodies conjugated to alkaline phosphatase (1:2000, sheep polyclonal, Sigma-Aldrich, catalog no. 11093274910, RRID: AB_2734716). A staining solution containing 5-bromo-4-chloro-3-indolyl phosphate and nitro blue tetrazolium was applied for 1 to 8 days at RT to visualize hybridization. Last, slides were mounted using Dako Faramount aqueous medium (Agilent). Images of stained sections were acquired using a 3DHISTECH Pannoramic 250 FLASH III digital slide scanner and processed with SlideViewer v2.6 (3DHISTECH, Hungary). Images were subsequently postprocessed in Adobe Photoshop CC, and ISH staining was enhanced for visualization by artificially coloring selected blue pixels using the “Select Color Range” and “Brush” tools. Whole-mount samples were fixed in 1% PFA and imaged using a Leica M205 FCA stereomicroscope; images were captured with a Leica DFC7000T camera and further processed using Leica Application Suite X software (v3.7.5.24914). Following whole-mount imaging, stained tooth buds were processed into paraffin blocks to confirm their developmental stage, sectioned, counterstained with H&E, and visualized in the same manner as other sections.

### 3D volume reconstruction of tooth buds and quantitative analysis of *SHH* expression

3D volumes of tooth buds and *SHH* expression domains were reconstructed from serial sagittal sections (5 µm thick) of entire *A. carolinensis* heads processed for *SHH* by ISH and counterstained with H&E. Raw images of stained sections were acquired using a 3DHISTECH Pannoramic 250 FLASH III digital slide scanner and processed with SlideViewer v2.6 (3DHISTECH, Hungary). Low-resolution snapshots of the entire jaw from each section were first captured and sequentially stacked in Inkscape v1.4.3 software to identify individual tooth buds and determine their developmental stages. High-resolution TIFF images of selected tooth buds were then acquired at identical size and resolution using SlideViewer v2.6. TIFF files were imported into Amira 5.5.0 as a z-stack using a script specifying image directory, pixel size, and z-positions based on section thickness. Sections were initially aligned using the “AlignSlices” tool, with manual correction when necessary. Tooth buds and *SHH* expression domains were manually segmented using the “brush” tool. 3D volumes were generated from segmented labels using the “SurfaceGen” function without smoothing. Surface area and volume measurements were obtained using the “Measure → SurfaceArea” function, which provides triangle count, surface area, and volume values. Volume measurements were used for subsequent analyses. Ratios of *SHH* domain volume to tooth bud volume were compared using the randomized test, with significance defined as *P* value <0.05. Statistical analyses were performed in R v4.3.2.

### Transcriptome analysis

Differential gene expression analyses of *EDA*-mutant *P. vitticeps* dental tissues were performed using previously published RNA-seq datasets generated in our earlier study (33). These datasets were obtained from laser-capture microdissection of acrodont dental epithelial tissue from early tooth germs (*n* = 3). Details of sample collection, RNA extraction, library preparation, and sequencing are provided in the original publication. Differentially expressed genes identified in the present study were subjected to Gene Ontology (GO) functional annotation using tools available through the Gene Ontology Consortium (http://geneontology.org/). In addition, these genes were compared with a curated list of keystone regulators of mammalian tooth development (34).

### BITES implementation and optimization procedure

BITES is an inverse-modeling framework that integrates a generative model of tooth morphogenesis with Bayesian parameter optimization to identify developmental parameter combinations capable of reproducing empirical tooth shapes. The framework couples a morphodynamic tooth model, which integrates signaling and tissue growth to simulate tooth development (36), with an automated optimization pipeline implemented in Python. Parameter optimization is performed using the Optuna library (66), a flexible framework for efficient hyperparameter optimization using Bayesian algorithms. In this approach, the morphodynamic tooth model is treated as a deterministic generative model whose final tooth morphology depends on a set of adjustable developmental parameters. For each parameter set proposed by the optimizer, the model is executed to generate a three-dimensional surface mesh representing the epithelial-mesenchymal boundary of the developing tooth. Simulated tooth shapes are quantitatively compared to predefined empirical target shapes using a landmark-free similarity metric. Empirical target shapes are provided as 3D surface meshes representing desired output morphologies, supplied in standard polygon formats (.ply or .off), allowing optimization against arbitrary tooth shapes that the model is capable of reproducing. The optimization process further supports extensive configuration beyond target selection, including control of simulation stages evaluated, alignment strategies, and optimization policies (see the simplified schematic overview of the optimization process structure in Fig. S6). Surface meshes are converted to point clouds and aligned to minimize differences due to translation, rotation, and scale using a two-step alignment procedure. First, a lightweight heuristic pre-alignment (STatac) is applied, involving scaling based on anterior-posterior and buccal-lingual crown dimensions, translation to align bounding-box centers or centroids, and alignment of crown apices along the apico-basal axis. This is followed, when enabled, by Coherent Point Drift (CPD), a probabilistic point-cloud registration algorithm (67, 68), to refine alignment. Shape similarity is then quantified using symmetric Chamfer Distance (69–71), as implemented in the Point Cloud Utils library (72). For each optimization trial, the lowest resulting distance across alignment attempts and simulation outputs is retained as the loss value. Optimization is performed in parallel across multiple CPU cores, enabling efficient exploration of parameter space. Sets of parameters yielding low loss values are interpreted as alternative developmental solutions capable of generating tooth morphologies closely matching empirical specimens. Auxiliary scripts with user-friendly graphical interfaces were developed to facilitate configuration of optimization settings and visualization of optimization results. A detailed step-by-step user guide with practical examples to facilitate application of the framework to new datasets is provided on GitHub (https://github.com/jupander/BITES).

### Normalization and variability analysis of *in silico* modeling results

For each of the six species (*A. carolinensis*, *A. ameiva*, *C. calyptratus*, *P. vitticeps*, *T. sexlineatus*, and *G. skoogi*), three independent *in silico* modeling runs were performed in BITES. From each run, the best-fitting model tooth was selected, yielding three independently derived models and their corresponding parameter sets for each species based on the same target tooth. Because model parameters varied substantially in magnitude, ranging from small decimal values to values in the thousands, parameter values were normalized to a [0,1] scale using min-max scaling to facilitate comparison and visualization. Normalization was calculated as follows: xnorm = (x − xmin)/(xmax − xmin), where x is the parameter value for a given model output and xmin and xmax are the minimum and maximum bounds of that parameter in the model. For each parameter within each species, variability across the three independently derived models was quantified as the range of normalized values (xnorm_max − xnorm_min). Normalization and initial calculations were performed in Microsoft Excel. The resulting parameter variability values were then imported into R v4.3.2, where they were ordered by variability across the six species. Box plots were generated to visualize interspecific variation in model parameter variability.

## ACKNOWLEDGEMENTS

We thank A.-C. Aho, M. Partanen, T. Ahlskog and J. Sireeni for technical assistance in captive breeding and animal care; H. Suhonen (University of Helsinki, Finland) for access to microCT scanning facilities; M. and P. Joki for access to veiled chameleon eggs; the Light Microscopy Unit (University of Helsinki, Finland) for access to imaging facility; the HiLAPS platform (University of Helsinki, Finland) for access to histology and histoscanner facilities; Genome Biology Unit, FIMM Digital Pathology Unit, and BI Histology core facility supported by HiLIFE from the University of Helsinki for scanning of histological samples; the DNA Sequencing and Genomics facility (University of Helsinki, Finland) for access to sequencing facility; J. Ollonen and F. Lafuma for some CT-scans; L. Scheinberg for providing access to *Gerrhosaurus skoogi* CT data from www.MorphoSource.org (California Academy of Sciences Herpetology collection, IP holder California Academy of Sciences); J. Eymann, L. Salomies, F. Sciuto, L. Björk, and S. Sorri for technical assistance; Teemu Häkkinen, P. Hagolani, and R. Zimm for morphodynamic model discussions; R. Linna for discussions on BITES implementation; J. Jernvall and D. Rice for helpful discussions throughout the project; and members of the Di-Poï laboratory for early discussions related to this project and assistance with morphometric analyses.

## Author contributions

D.R. and N.D.-P. designed the overall experimental approach and selected the species sampling. D.R. and I.-M.A contributed to embryonic and/or postnatal specimen collection and preparation. MicroCT scans were carried out by I.-M.A. I.-M.A. collected the 3D landmark data and performed geometric morphometric and tooth measurement analyses. J.I. developed the BITES software and prepared the detailed user guide, with guidance from D.R. and N.D.-P. D.R. performed all developmental and molecular analyses. D.R. and N.D.-P. created the figures and wrote the manuscript. All co-authors contributed in the form of discussion and critical comments. All authors approved the final version of the manuscript.

## Data and materials availability

All data needed to evaluate the conclusions in the paper are present in the paper and/or the Supporting Information. The transcriptomic analyses were performed using raw RNA-seq data generated in our previous work (33). All raw Illumina reads are deposited on the Gene Expression Omnibus (GEO) database under the accession number GSE173967. A GitHub repository has been created to host BITES, along with example target shape files and a detailed step-by-step user guide (https://github.com/jupander/BITES).

## REFERENCES

1. T. H. Frazzetta, The mechanics of cutting and the form of shark teeth (Chondrichthyes, Elasmobranchii). Zoomorphology 108, 93–107 (1988).

2. J. T. Streelman, J. F. Webb, R. C. Albertson, T. D. Kocher, The cusp of evolution and development: A model of cichlid tooth shape diversity. Evol. Dev. 5, 600–608 (2003).

3. G. J. Fraser, R. F. Bloomquist, J. T. Streelman, Common developmental pathways link tooth shape to regeneration. Dev. Biol. 377, 399–414 (2013).

4. N. Peoples, P. C. Wainwright, Multifaceted impacts of an innovation on dental diversity in an adaptive radiation of cichlid fishes. Proc. R. Soc. B Biol. Sci. 292, 20252208 (2025).

5. R. Zimm, V. Tobias Santos, N. Goudemand, Integration of multi-level dental diversity links macro-evolutionary patterns to ecological strategies across sharks. Elife 14, 1–24 (2025).

6. T. Davit-Béal, H. Chisaka, S. Delgado, J. Y. Sire, Amphibian teeth: current knowledge, unanswered questions, and some directions for future research. Biol. Rev. 82, 49–81 (2007).

7. D. J. Paluh, et al., Rampant tooth loss across 200 million years of frog evolution. Elife 10, e66926 (2021).

8. O. Zahradnicek, M. Buchtova, H. Dosedelova, A. S. Tucker, The development of complex tooth shape in reptiles. Front. Physiol. 5, 1–7 (2014).

9. K. M. Melstrom, The relationship between diet and tooth complexity in living dentigerous saurians. J. Morphol. 278, 500–522 (2017).

10. F. Lafuma, I. J. Corfe, J. Clavel, N. Di-Poï, Multiple evolutionary origins and losses of tooth complexity in squamates. Nat. Commun. 12, 1–13 (2021).

11. J. Jernvall, I. Thesleff, Tooth shape formation and tooth renewal: Evolving with the same signals. Dev. 139, 3487–3497 (2012).

12. J. Jernvall, P. Kettunen, I. Karavanova, L. B. Martin, I. Thesleff, Evidence for the role of the enamel knot as a control center in mammalian tooth cusp formation: Non-dividing cells express growth stimulating Fgf-4 gene. Int. J. Dev. Biol. 38, 463–469 (1994).

13. T. Porntaveetus, et al., Expression of fibroblast growth factors (Fgfs) in murine tooth development. J. Anat. 218, 534–543 (2011).

14. E. Harjunmaa, et al., Replaying evolutionary transitions from the dental fossil record. Nature 512, 44–48 (2014).

15. M. Buchtová, J. C. Boughner, K. Fu, V. M. Diewert, J. M. Richman, Embryonic development of Python sebae - II: Craniofacial microscopic anatomy, cell proliferation and apoptosis. Zoology 110, 231–251 (2007).

16. M. Buchtová, et al., Initiation and patterning of the snake dentition are dependent on Sonic Hedgehog signaling. Dev. Biol. 319, 132–145 (2008).

17. G. R. Handrigan, J. M. Richman, Autocrine and paracrine Shh signaling are necessary for tooth morphogenesis, but not tooth replacement in snakes and lizards (Squamata). Dev. Biol. 337, 171–186 (2010).

18. G. R. Handrigan, J. M. Richman, Unicuspid and bicuspid tooth crown formation in squamates. J. Exp. Zool. Part B Mol. Dev. Evol. 316 **B**, 598–608 (2011).

19. J. M. Richman, G. R. Handrigan, Reptilian tooth development. Genesis 49, 247–260 (2011).

20. M. Landova Sulcova, et al., Developmental mechanisms driving complex tooth shape in reptiles. Dev. Dyn. 1–24 (2019). 10.1002/dvdy.138.

21. M. Buchtová, O. Zahradníček, S. Balková, A. S. Tucker, Odontogenesis in the Veiled Chameleon (Chamaeleo calyptratus). Arch. Oral Biol. 58, 118–133 (2013).

22. B. Westergaard, M. W. J. Ferguson, Development of the dentition in Alligator mississippiensis. Later development in the lower jaws of embryos, hatchlings and young juveniles. J. Zool. 212, 191–222 (1987).

23. O. Weeks, B. A. S. Bhullar, A. Abzhanov, Molecular characterization of dental development in a toothed archosaur, the American alligator Alligator mississippiensis. Evol. Dev. 15, 393–405 (2013).

24. M. Debiais-Thibaud, et al., Tooth and scale morphogenesis in shark: An alternative process to the mammalian enamel knot system. BMC Evol. Biol. 15, 1–17 (2015).

25. G. J. Fraser, R. F. Bloomquist, J. T. Streelman, A periodic pattern generator for dental diversity. BMC Biol. 6, 32 (2008).

26. W. R. Jackman, B. W. Draper, D. W. Stock, Fgf signaling is required for zebrafish tooth development. Dev. Biol. 274, 139–157 (2004).

27. R. F. Bloomquist, Developmental basis of natural tooth shape variation in cichlid fishes. Naturwissenschaften 112, 12 (2025).

28. L. J. Rasch, et al., An ancient dental gene set governs development and continuous regeneration of teeth in sharks. Dev. Biol. 415, 347–370 (2016).

29. A. P. Thiery, A. S. I. Standing, R. L. Cooper, G. J. Fraser, An epithelial signalling centre in sharks supports homology of tooth morphogenesis in vertebrates. Elife 11, 1–29 (2022).

30. D. Razmadze, L. Salomies, N. Di-Poï, Squamates as a model to understand key dental features of vertebrates. Dev. Biol. 516, 1–19 (2024).

31. L. Salomies, J. Eymann, I. Khan, N. Di-Po, The alternative regenerative strategy of bearded dragon unveils the key processes underlying vertebrate tooth renewal. Elife 8 (2019).

32. N. Di-Poï, M. C. Milinkovitch, The anatomical placode in reptile scale morphogenesis indicates shared ancestry among skin appendages in amniotes. Sci. Adv. 2, e1600708 (2016).

33. L. Salomies, J. Eymann, J. Ollonen, I. Khan, N. Di-Poï, The developmental origins of heterodonty and acrodonty as revealed by reptile dentitions. Sci. Adv. 7, 1–14 (2021).

34. O. Hallikas, et al., System - level analyses of keystone genes required for mammalian tooth development. J. Exp. Zool. Part B Mol. Dev. Evol. 336, 7–17 (2021).

35. M. M. Christensen, O. Hallikas, R. Das Roy, J. Jernvall, The developmental basis for scaling of mammalian tooth size. Proc. Natl. Acad. Sci. 120, e2300374120 (2023).

36. I. Salazar-Ciudad, J. Jernvall, A computational model of teeth and the developmental origins of morphological variation. Nature 464, 583–586 (2010).

37. R. Zimm, F. Berio, M. Debiais-Thibaud, N. Goudemand, A shark-inspired general model of tooth morphogenesis unveils developmental asymmetries in phenotype transitions. PNAS 120, e2216959120 (2023).

38. J. Pispa, et al., Cusp patterning defect in Tabby mouse teeth and its partial rescue by FGF. Dev. Biol. 216, 521–534 (1999).

39. G. J. Fraser, R. F. Bloomquist, J. T. Streelman, Common developmental pathways link tooth shape to regeneration. Dev. Biol. 377, 399–414 (2013).

40. B. M. Davis, Evolution of the tribosphenic molar pattern in early mammals, with comments on the “dual-origin” hypothesis. J. Mamm. Evol. 18, 227–244 (2011).

41. A. R. Evans, G. D. Sanson, The tooth of perfection : functional and spatial constraints on mammalian tooth shape. Biol. J. Linn. Soc. 78, 173–191 (2003).

42. M. Fortelius, Ungulate cheek teeth: developmental, functional, and evolutionary interrelations. Acta Zool. Fenn. 180, 1–76 (1985).

43. J. P. Hunter, J. Jernvallt, The hypocone as a key innovation in mammalian evolution. Proc. Natl. Acad. Sci. 92, 10718–10722 (1995).

44. A. R. Evans, G. P. Wilson, M. Fortelius, J. Jernvall, High-level similarity of dentitions in carnivorans and rodents. Nature 445, 78–81 (2007).

45. P. S. Ungar, Mammal Teeth: Origin, Evolution, and Diversity (The Johns Hopkins University Press, 2010).

46. G. P. Wilson, et al., Adaptive radiation of multituberculate mammals before the extinction of dinosaurs. Nature 483, 457–460 (2012).

47. T. J. Sanger, J. B. Losos, J. J. Gibson-Brown, A developmental staging series for the lizard genus Anolis: a new system for the integration of evolution, development, and ecology. J. Morphol. 269, 129–137 (2008).

48. J. Ollonen, F. O. Da Silva, K. Mahlow, N. Di-Poï, Skull development, ossification pattern, and adult shape in the emerging lizard model organism Pogona vitticeps: a comparative analysis with other squamates. Front. Physiol. 9, 278 (2018).

49. R. E. Diaz Jr, P. A. Trainor, N. A. Shylo, D. Roellig, M. Bronner, Filling in the phylogenetic gaps : Induction, migration, and differentiation of neural crest cells in a squamate reptile, the veiled chameleon (Chamaeleo calyptratus). Dev. Dyn. 248, 709–727 (2019).

50. K. Okuyama, Y. Sakuma, T. Sasaki, Post-ovipositional developmental stages of the Japanese grass lizard, Takydromus tachydromoides (Squamata: Lacertidae). Curr. Herpetol. 40, 66–76 (2021).

51. R. A. Pyron, F. Burbrink, J. Wiens, A phylogeny and revised classification of Squamata, including 4161 species of lizards and snakes. BMC Evol. Biol. 13, 93 (2013).

52. R. Kikinis, S. D. Pieper, K. G. Vosburgh, “3D Slicer: A Platform for Subject-Specific Image Analysis, Visualization, and Clinical Support” in Intraoperative Imaging and Image-Guided Therapy, F. Jolesz, Ed. (Springer, 2014).

53. S. Rolfe, et al., SlicerMorph: An open and extensible platform to retrieve, visualize and analyse 3D morphology. Methods Ecol. Evol. 12, 1816–1825 (2021).

54. R Core Team, R: A language and environment for statistical computing. R Found. Stat. Comput. Vienna, Austria (2021). Available at: https://cir.nii.ac.jp/crid/1370013168792282134.bib?lang=en [Accessed 25 January 2026].

55. P. Gunz, P. Mitteroecker, Semilandmarks: a method for quantifying curves and surfaces Point homology Placing semilandmarks. Hystrix, Ital. J. Mammal. 24, 103–109 (2013).

56. S. Schlager, “Morpho and Rvcg – Shape Analysis in R: R-Packages for Geometric Morphometrics, Shape Analysis and Surface Manipulations” in Statistical Shape and Deformation Analysis: Methods, Implementation and Applications, G. Zheng, S. Li, G. Szekely, Eds. (Academic Press, 2017), pp. 217–256.

57. D. C. Adams, E. Otárola-Castillo, Geomorph: An r package for the collection and analysis of geometric morphometric shape data. Methods Ecol. Evol. 4, 393–399 (2013).

58. N. Di-Poï, M. C. Milinkovitch, Crocodylians evolved scattered multi-sensory micro-organs. Evodevo 4, 1–16 (2013).

59. J. Eymann, L. Salomies, S. Macrì, Variations in the proliferative activity of the peripheral retina correlate with postnatal ocular growth in squamate reptiles. 2356–2370 (2019). 10.1002/cne.24677.

60. J. Alföldi, et al., The genome of the green anole lizard and a comparative analysis with birds and mammals. Nature 477, 587–591 (2011).

61. A. J. Geneva, et al., Chromosome-scale genome assembly of the brown anole (Anolis sagrei), an emerging model species. Commun. Biol. 5, 1–13 (2022).

62. A. Georges, et al., High-coverage sequencing and annotated assembly of the genome of the Australian dragon lizard Pogona vitticeps. Gigascience 4 (2015).

63. B. J. Pinto, et al., The revised reference genome of the leopard gecko (Eublepharis macularius) provides insight into the considerations of genome phasing and assembly. J. Hered. 114, 513–520 (2023).

64. N. Koochekian, et al., A chromosome-level genome assembly and annotation of the desert horned lizard, Phrynosoma platyrhinos, provides insight into chromosomal rearrangements among reptiles. Gigascience 11, 1–14 (2022).

65. A. A. Yurchenko, H. Recknagel, K. R. Elmer, Chromosome-level assembly of the common lizard (Zootoca vivipara) genome. Genome Biol. Evol. 12, 1953–1960 (2020).

66. T. Akiba, S. Sano, T. Yanase, T. Ohta, M. Koyama, Optuna: A next-generation hyperparameter optimization framework. Proc. ACM SIGKDD Int. Conf. Knowl. Discov. Data Min. 2623–2631 (2019). 10.1145/3292500.3330701.

67. A. Myronenko, X. Song, Point Set Registration : Coherent Point Drift. IEEE Trans. Pattern Anal. Mach. Intell. 32, 2262–2275 (2010).

68. A. A. Gatti, S. Khallaghi, PyCPD : Pure NumPy Implementation of the Coherent Point Drift Algorithm. J. Open Source Softw. 7, 4681 (2022).

69. H. G. Barrow, J. M. Tenenbaum, R. C. Bolles, H. C. Wolf, Parametric correspondence and chamfer matching: Two new techniques for image matching. Proc. Image Underst. Work. 21–27 (1977).

70. B. A. Young, K. V. Kardong, Dentitional surface features in snakes (Reptilia: Serpentes). Amphib. Reptil. 17, 261–276 (1996).

71. H. Fan, H. Su, L. J. Guibas, A point set generation network for 3D object reconstruction from a single image in Proceedings of the IEEE Conference on Computer Vision and Pattern Recognition., (2017), pp. 605–613.

72. F. Williams, Point cloud utils. (2022). Available at: https://github.com/fwilliams/point-cloud-utils.

