## Supplementary Information for "An alternative patterning mechanism for vertebrate tooth complexity"

### Supporting Information

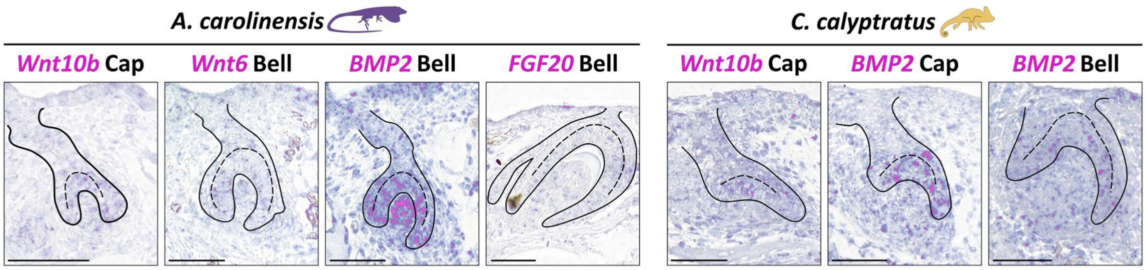


**Fig. S1**. **Epithelial signaling in *Anolis carolinensis* and *Chamaeleo calyptratus.*** ISH expression patterns of conserved EK-associated genes. From left to right: in *A. carolinensis, Wnt10b* at the cap stage, *Wnt6* at the bell stage, *BMP2* at the bell stage, and *FGF20* at the bell stage; in *C. calyptratus, Wnt10b* at the cap stage, *BMP2* at the cap and bell stages. Scale bars: 50 µm.

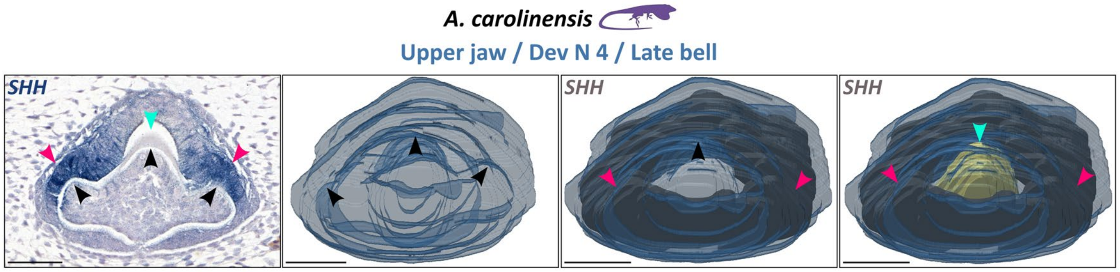


**Fig. S2. 3D visualization of *SHH* expression at the late bell stage in a developing tricuspid tooth of *A. carolinensis*.** From left to right: ISH (*SHH*) and hematoxylin-stained sagittal histological section; 3D reconstruction of the tooth bud; 3D reconstruction of the tooth bud with *SHH* expression; and 3D reconstruction of the tooth bud with *SHH* expression and mineralizing tooth tissue. The dental epithelium is shown in a color corresponding to the posterior tooth position in the jaw (as in Fig. 3), the *SHH* signal is shown in grey, and mineralized tissue is shown in yellow. Black arrowheads indicate the positions of cusps; pink arrowheads indicate *SHH* expression associated with developing additional cusps; light blue arrowheads indicate the mineralized main cusp. A 2D section passing through all three developing cusps gives the impression that *SHH* expression is localized in the region of secondary cusp formation, resembling a secondary enamel knot in 2D. However, 3D reconstruction of the entire tooth bud and the *SHH* expression domain reveals a broad ring-like expression field. Scale bars: 50 μm.

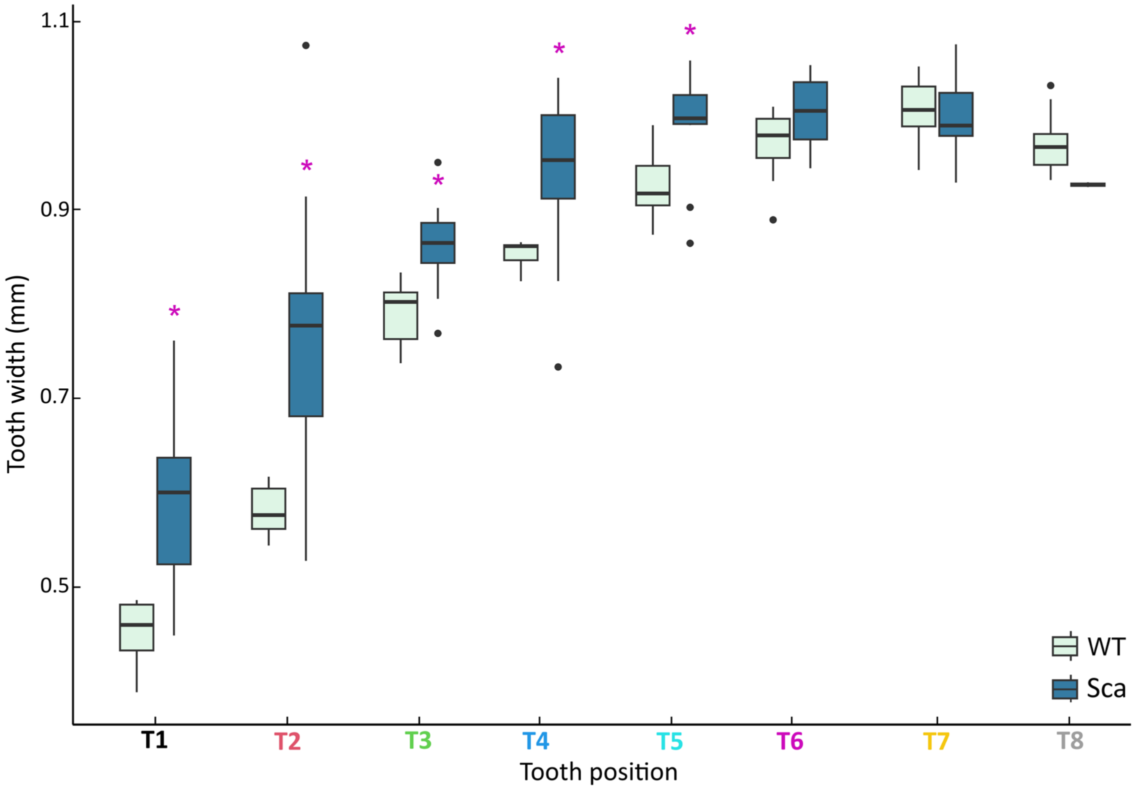


**Fig. S3**. **Tooth size by tooth position in wild-type and scaleless mutant *P. vitticeps*.** Box plots showing tooth width (mm) at each of the eight acrodont tooth positions along the jaw in hatchlings of WT and Sca mutants (n = 5 animals per group). Tooth positions are arranged along the x-axis from anterior (left) to posterior (right). For each tooth position, WT box plots are shown on the left and colored light blue, while Sca mutant box plots are shown on the right and colored dark blue. Purple asterisks indicate tooth positions showing a significant difference ($P < 0.05$) in tooth width between WT and Sca mutants.

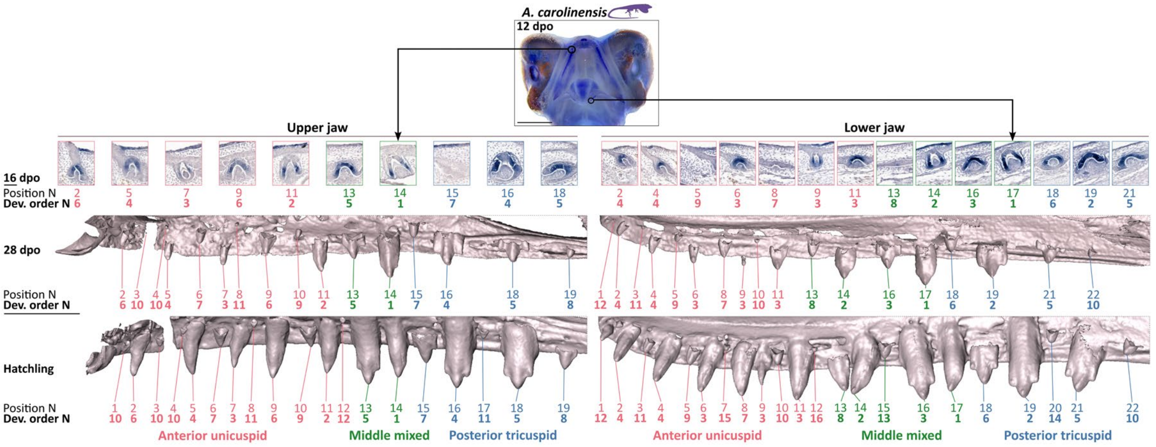


**Fig. S4. Tooth row patterning in *A. carolinensis*.** Top: WMISH (*SHH*) in the embryonic head of *A. carolinensis* at 12 days post-oviposition (dpo), showing the first functional tooth developing in the upper and lower jaws, together with two more anteriorly located vestigial teeth (as shown for the upper jaw in Fig. 1). Top row: ISH (*SHH*) and hematoxylin-stained sagittal histological sections of each developing tooth from the upper (left) and lower (right) jaws in an embryo at 16 dpo. Middle row: 3D microCT reconstructions of embryonic upper (left) and lower (right) jaws with the developing tooth row shown in lingual view at 28 dpo. Bottom row: 3D microCT reconstructions of hatchling upper (left) and lower (right) jaws shown in lingual view. Position N denotes the tooth position number from anterior (left) to posterior (right). The hatchling tooth row is considered complete; teeth are numbered from 1 to 19 in the upper jaw and from 1 to 22 in the lower jaw. The same positional numbering is applied to homologous tooth positions identified in embryonic developing tooth rows. Dev. order N indicates the order of tooth initiation along the jaw (as in Fig. 3). Colors indicating tooth position and developmental order follow the location and shape gradient shown in Fig. 3. The position of the first developing tooth at the beginning of tooth-row formation and at the end of tooth-row development indicates intensive anterior jaw growth. The order of tooth initiation along the jaw indicates intercalary tooth raw formation, with new teeth appearing between previously initiated teeth. Scale bars: 1 mm (WMISH image), 50 µm (histological sections), 0.5 mm (microCT reconstructions).

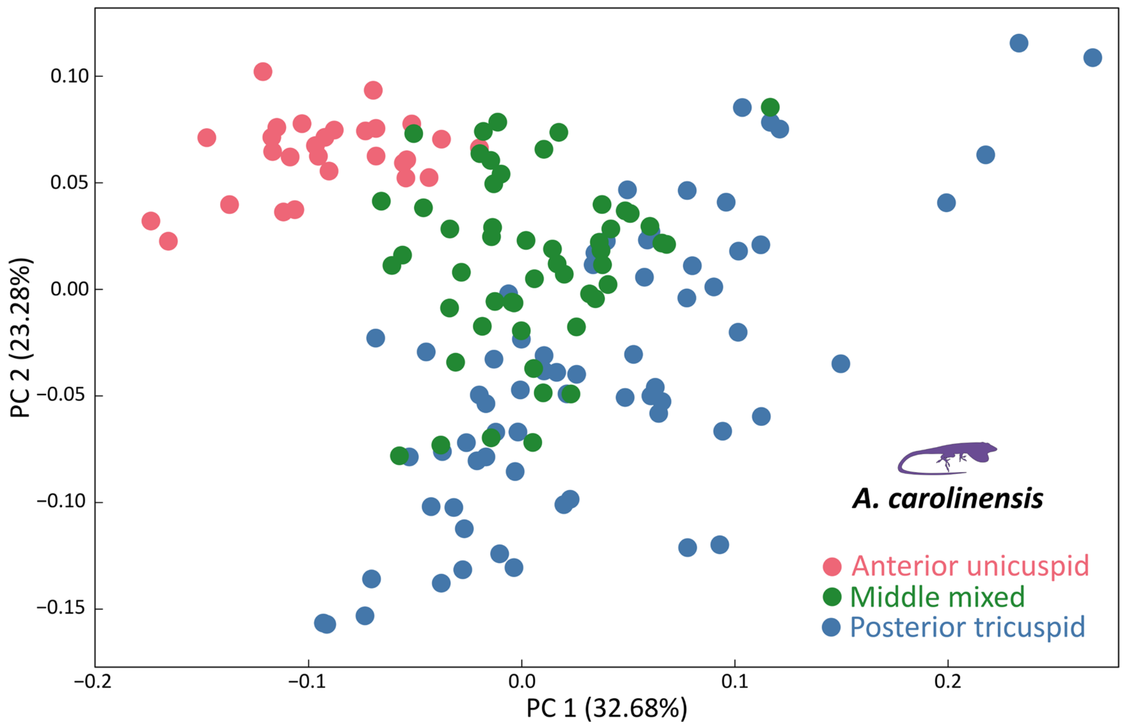


**Fig. S5**. **Tooth-shape morphospace in *A. carolinensis*.** Tooth-shape morphospace showing the distribution of anterior unicuspid teeth (brick red), teeth from middle variable intermediate zone (dark green) and broader posterior tricuspid teeth (blue) (n = 6 animals per group). Numbers in parentheses indicate the percentage of total shape variance explained by each principal component (PC) axis.

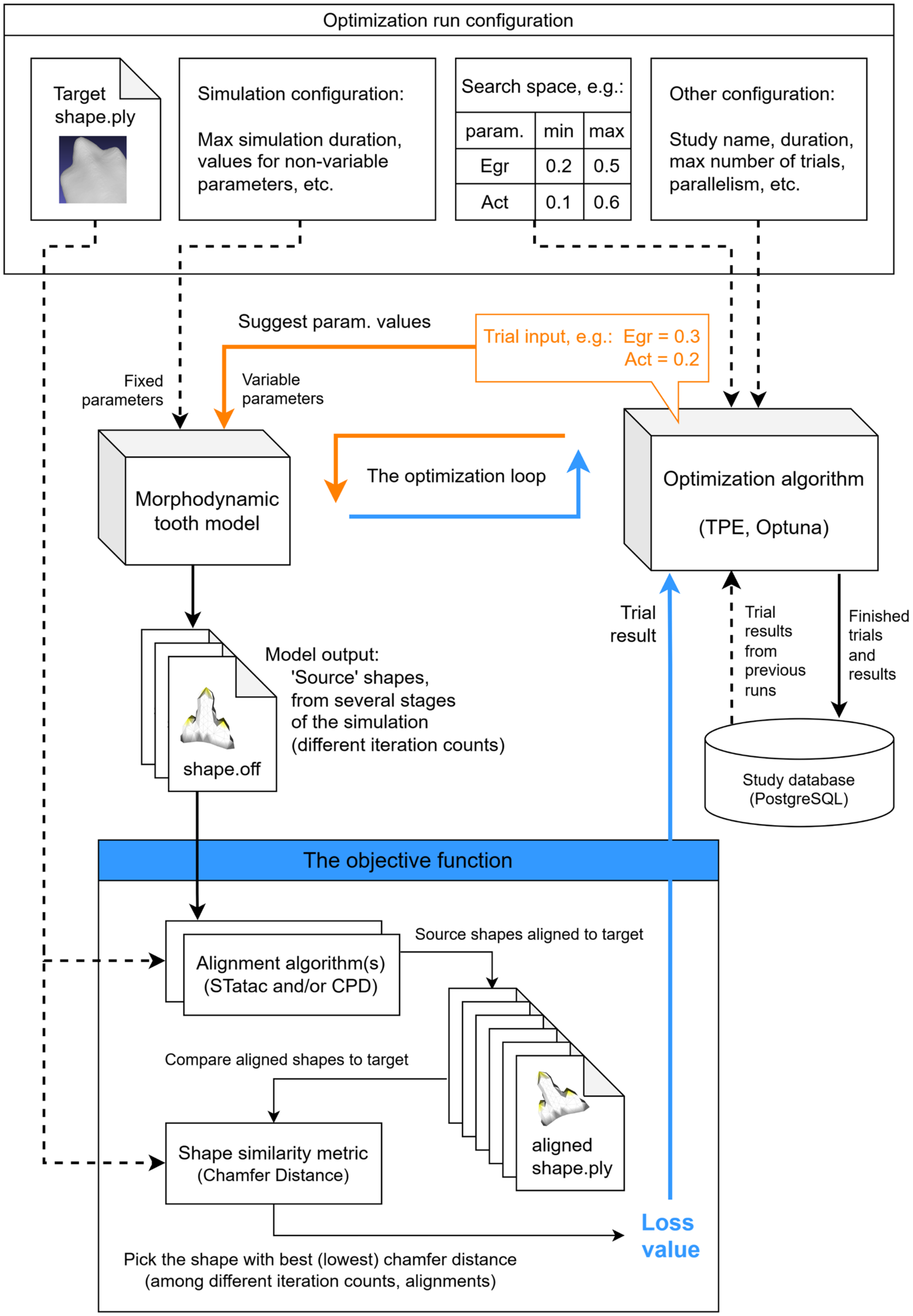


**Fig. S6. BITES: simplified overview of the optimization process structure**. Each run of the tooth model can output multiple shapes, each from a different stage of the simulation. The objective function will then evaluate each shape separately (optionally with multiple different alignment methods) and finally selects the one with the lowest loss as the canonical result of the trial.

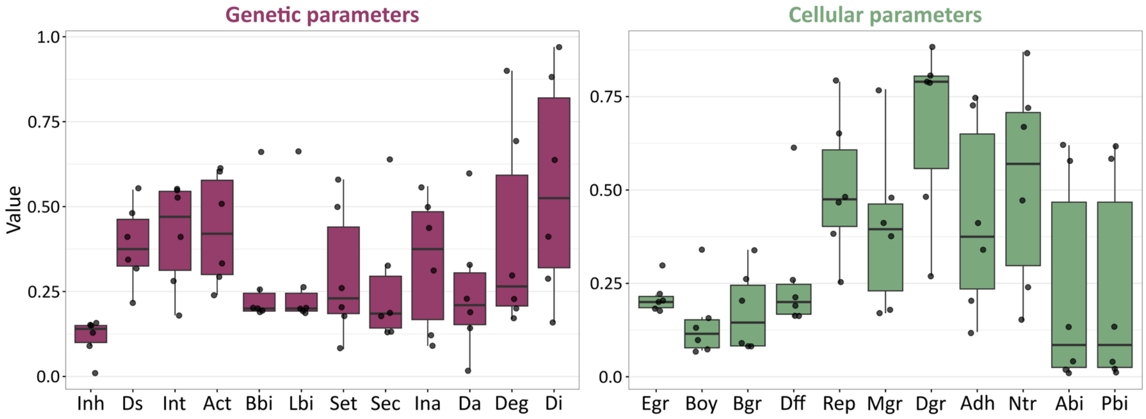


**Fig. S7. Variation in parameters during *in silico* optimization of convergent complex tooth shapes in six lizard species**. Variation index plots showing the percentage variation of all genetic parameters (left) and cellular parameters (right) across three independent optimization runs for complex tooth shapes in each of the six convergent species (*Gerrhosaurus skoogi, Ameiva ameiva, Takydromus sexlineatus, Chamaeleo calyptratus, Pogona vitticeps, and Anolis carolinensis*). Plots are organized from lowest to highest variation index. Parameter abbreviations as defined in (1): Inh, inhibition of activator; Ds, growth factor diffusion rate; Int, initial inhibitor threshold; Act, activator auto-activation; Bbi, buccal bias; Lbi, lingual bias; Set, growth factor threshold; Sec, growth factor secretion rate; Ina, initial activator concentration; Da, activator diffusion rate; Deg, protein degradation rate; Di, inhibitor diffusion rate; Egr, epithelial proliferation rate; Boy, mesenchymal mechanical resistance; Bgr, border growth (amount of mesenchyme in anterior-posterior direction); Dff, differentiation rate; Rep, Young's modulus (stiffness); Mgr, mesenchymal proliferation rate; Dgr, downward vector of growth; Adh, traction between neighbours; Ntr, mechanical traction from the borders to the nucleus; Abi, anterior bias; Pbi, posterior bias.

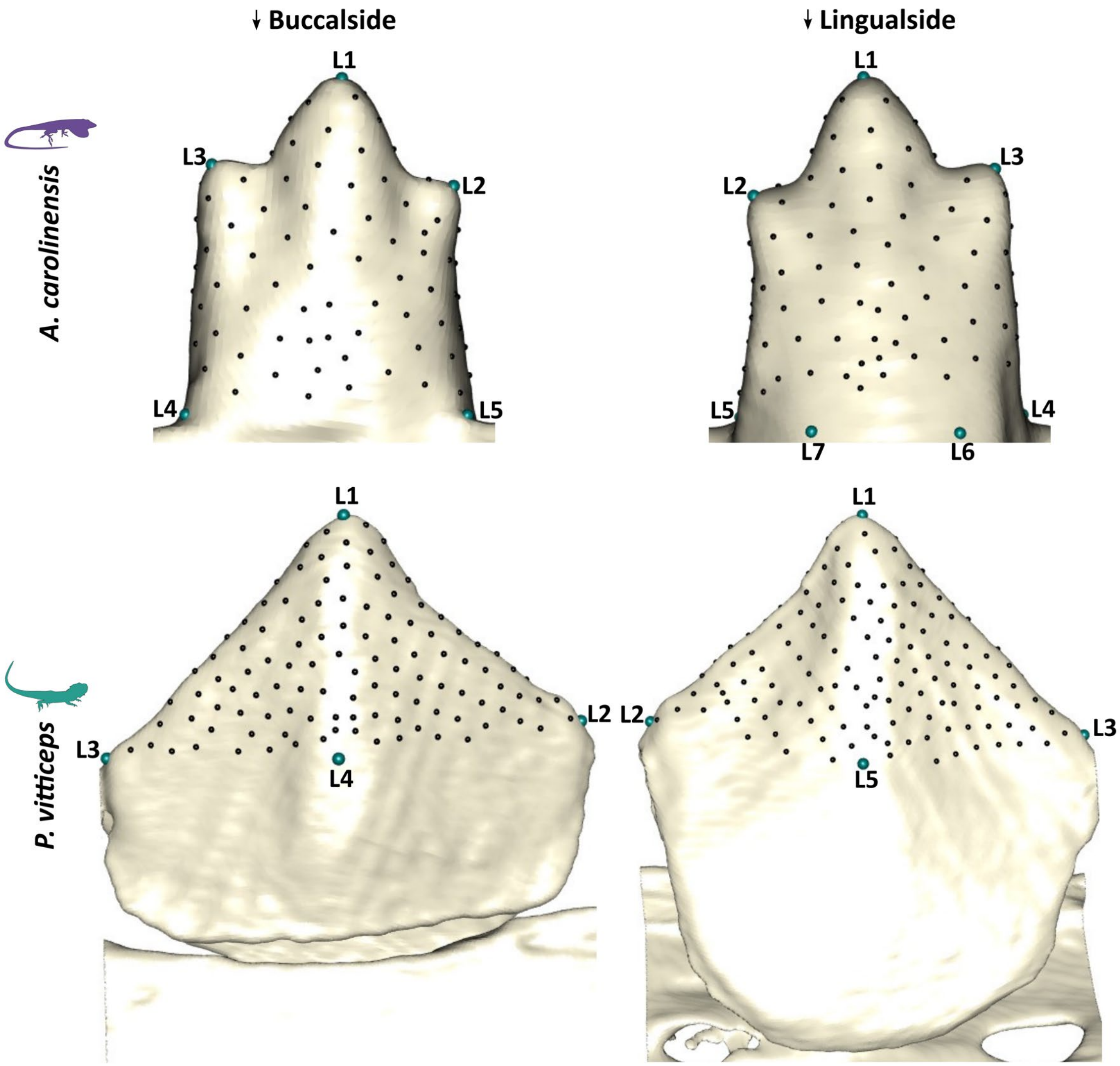


**Fig. S8**. **Landmarks placement on *A. carolinensis* and *P. vitticeps* teeth.** Top row: representative tricuspid tooth of *A. carolinensis* from buccal view (left) and lingual view (right). Bottom row: representative acrodont tooth of *P. vitticeps* from buccal view (left) and lingual view (right). L1-L7 in *A. carolinensis* and L1-L5 in *P. vitticeps* indicate fixed landmarks; black dots across the tooth surfaces indicate surface semi-landmarks.

**Table S1.** Width of acrodont teeth in WT and Sca *P. vitticeps* hatchlings (in μm). S, side; L, left; R, right.

| | Lower jaw | | | Upper jaw | | | | Lower jaw | | | Upper jaw | | |
|---|---|---|---|---|---|---|---|---|---|---|---|---|---|
| **Sp** | **Tooth** | **S** | **Width** | **Tooth** | **S** | **Width** | **Sp** | **Tooth** | **S** | **Width** | **Tooth** | **S** | **Width** |
| WT PV551 | T1 | L | 427,93 | T1 | L | 457,96 | Sca PV553 | T1 | L | 637,08 | T1 | L | 454,65 |
| | T2 | L | 581,69 | T2 | L | 753,81 | | T2Bifid | L | 1073,49 | T2 | L | 707,56 |
| | T3 | L | 813,12 | T3 | L | 827,13 | | T3 | L | 874,63 | T3 | L | 918,64 |
| | T4 | L | 862,39 | T4 | L | 953,46 | | T4 | L | 956,98 | T4 | L | 966,66 |
| | T5 | L | 968,23 | T5 | L | 965,45 | | T5 | L | 1012,55 | T5 | L | 952,13 |
| | T6 | L | 1005,36 | T6 | L | 1063,62 | | T6 | L | 1002,92 | T6 | L | 1012,72 |
| | T7 | L | 1051,22 | T7 | L | 1042,17 | | T7 | L | 928,39 | T7 | L | 1016,47 |
| | T8 | L | 931,07 | T8 | L | 948,31 | | - | - | - | T8 | L | 869,34 |
| | T1 | R | 448,44 | T1 | R | 497,04 | | T1 | R | 609,01 | T1 | R | 447,10 |
| | T2 | R | 559,65 | T2 | R | 756,15 | | T2 | R | 913,53 | T2 | R | 702,57 |
| | T3 | R | 808,66 | T3 | R | 802,13 | | T3 | R | 879,46 | T3 | R | 875,89 |
| | T4 | R | 849,67 | T4 | R | 993,27 | | T4 | R | 947,35 | T4 | R | 959,04 |
| | T5 | R | 930,75 | T5 | R | 985,53 | | T5 | R | 994,22 | T5 | R | 941,39 |
| | T6 | R | 1008,59 | T6 | R | 1077,74 | | T6 | R | 987,96 | T6 | R | 1015,09 |
| | T7 | R | 1015,34 | T7 | R | 1054,69 | | T7 | R | 982,25 | T7 | R | 1019,72 |
| | T8 | R | 981,09 | T8 | R | 892,80 | | - | - | - | T8 | R | 901,41 |
| WT PV555 | T1 | L | 470,89 | T1 | L | 480,13 | Sca PV554 | T1 | Left | 448,85 | T1 | L | 406,45 |
| | T2 | L | 616,97 | T2 | L | 702,41 | | T2 | Left | 527,87 | T2 | L | 704,62 |
| | T3 | L | 822,81 | T3 | L | 812,50 | | T3 | Left | 805,30 | T3 | L | 827,42 |
| | T4 | L | 861,29 | T4 | L | 874,29 | | T4 | Left | 824,10 | T4 | L | 932,85 |
| | T5 | L | 903,22 | T5 | L | 932,48 | | T5 | Left | 902,12 | T5 | L | 887,26 |
| | T6 | L | 978,13 | T6 | L | 988,21 | | T6 | Left | 969,31 | T6 | L | 941,89 |
| | T7 | L | 1034,62 | T7 | L | 981,84 | | T7 | Left | 984,72 | T7 | L | 965,72 |
| | T8 | L | 945,72 | T8 | L | 785,55 | | T8 | Left | 928,61 | T8 | L | 855,55 |
| | T1 | R | 481,70 | T1 | R | 454,87 | | T1 | R | 477,41 | T1 | R | 412,27 |
| | T2 | R | 613,89 | T2 | R | 688,36 | | T2 | R | 634,05 | T2 | R | 702,96 |
| | T3 | R | 807,27 | T3 | R | 833,52 | | T3 | R | 768,70 | T3 | R | 831,02 |
| | T4 | R | 838,32 | T4 | R | 917,62 | | T4 | R | 733,13 | T4 | R | 921,70 |
| | T5 | R | 873,17 | T5 | R | 916,96 | | T5 | R | 864,08 | T5 | R | 890,86 |
| | T6 | R | 978,50 | T6 | R | 986,51 | | T6 | R | 943,47 | T6 | R | 980,82 |
| | T7 | R | 986,85 | T7 | R | 940,01 | | T7 | R | 1001,04 | T7 | R | 975,42 |
| | T8 | R | 939,36 | T8 | R | 758,51 | | T8 | R | 923,66 | T8 | R | 879,15 |
| WT PV559 | T1 | L | 400,97 | T1 | L | 464,08 | Sca PV558 | T1 | L | 524,19 | T1 | L | 607,75 |
| | T2 | L | 570,94 | T2 | L | 683,40 | | T2 | L | 736,41 | T2 | L | 965,44 |
| | T3 | L | 744,14 | T3 | L | 812,62 | | T3 | L | 949,63 | T3 | L | 1012,76 |
| | T4 | L | 863,60 | T4 | L | 930,76 | | T4 | L | 1031,40 | T4 | L | 973,37 |
| | T5 | L | 873,95 | T5 | L | 846,06 | | T5 | L | 998,50 | T5 | L | 1052,43 |
| | T6 | L | 976,44 | T6 | L | 996,09 | | T6 | L | 1039,78 | T6 | L | 1077,00 |

| | | | | | | | | | | | | | |
|---|---|---|---|---|---|---|---|---|---|---|---|---|---|
| | T7 | L | 941,63 | T7 | L | 1014,43 | | T7 | L | 1030,75 | T7 | L | 849,91 |
| | T8 | L | 951,20 | T8 | L | 936,84 | | – | – | – | – | – | – |
| | T1 | R | 388,82 | T1 | R | 464,76 | | T1 | R | 524,19 | T1 | R | 583,38 |
| | T2 | R | 603,53 | T2 | R | 669,20 | | T2 | R | 776,92 | T2 | R | 973,85 |
| | T3 | R | 781,58 | T3 | R | 839,19 | | T3 | R | 901,51 | T3 | R | 1045,16 |
| | T4 | R | 844,95 | T4 | R | 945,92 | | T4 | R | 1039,24 | T4 | R | 1017,90 |
| | T5 | R | 912,43 | T5 | R | 860,31 | | T5 | R | 1030,88 | T5 | R | 1053,73 |
| | T6 | R | 929,95 | T6 | R | 1016,95 | | T6 | R | 1019,05 | T6 | R | 1054,57 |
| | T7 | R | 990,72 | T7 | R | 996,59 | | T7 | R | 992,70 | T7 | R | 864,88 |
| | T8 | R | 968,25 | T8 | R | 891,11 | | – | – | – | – | – | – |
| WT PV695 | T1 | L | 449,23 | T1 | L | 449,49 | Sca PV560 | T1 | L | 643,08 | T1 | L | 426,87 |
| | T2 | L | 560,30 | T2 | L | 689,59 | | T2 | L | 811,17 | T2 | L | 733,32 |
| | T3 | L | 756,38 | T3 | L | 837,69 | | T3 | L | 841,56 | T3 | L | 853,76 |
| | T4 | L | 865,19 | T4 | L | 937,02 | | T4 | L | 902,31 | T4 | L | 949,56 |
| | T5 | L | 907,16 | T5 | L | 884,17 | | T5 | L | 989,15 | T5 | L | 878,39 |
| | T6 | L | 947,00 | T6 | L | 1041,41 | | T6 | L | 965,74 | T6 | L | 1003,66 |
| | T7 | L | 984,76 | T7 | L | 1061,89 | | T7 | L | 1074,64 | T7 | L | 1030,92 |
| | T8 | L | 1031,06 | T8 | L | 1036,06 | | – | – | – | T8 | L | 892,07 |
| | T1 | R | 483,73 | T1 | R | 455,27 | | T1 | R | 600,33 | T1 | R | 437,88 |
| | T2 | R | 604,65 | T2 | R | 685,14 | | T2 | R | 791,56 | T2 | R | 762,30 |
| | T3 | R | 737,15 | T3 | R | 883,46 | | T3 | R | 887,81 | T3 | R | 862,16 |
| | T4 | R | 860,77 | T4 | R | 970,56 | | T4 | R | 938,45 | T4 | R | 940,15 |
| | T5 | R | 921,19 | T5 | R | 883,65 | | T5 | R | 993,84 | T5 | R | 933,62 |
| | T6 | R | 888,88 | T6 | R | 1027,12 | | T6 | R | 1005,60 | T6 | R | 969,58 |
| | T7 | R | 995,24 | T7 | R | 1059,49 | | T7 | R | 1053,69 | T7 | R | 1062,52 |
| | T8 | R | 1016,53 | T8 | R | 1032,80 | | – | – | – | T8 | R | 934,82 |
| WT PV696 | T1 | L | 486,28 | T1 | L | 470,85 | Sca PV678 | T1 | L | 761,09 | T1 | L | 504,30 |
| | T2 | L | 544,11 | T2 | L | 700,28 | | – | – | – | T2 | L | 928,13 |
| | T3 | L | 833,18 | T3 | L | 825,01 | | T3 | L | 848,01 | T3 | L | 973,16 |
| | T4 | L | 861,26 | T4 | L | 963,53 | | T4 | L | 977,21 | T4Bifid | L | 1063,20 |
| | T5 | L | 989,08 | T5 | L | 925,16 | | T5 | L | 1023,85 | T5 | L | 940,38 |
| | T6 | L | 999,20 | T6 | L | 1075,11 | | T6 | L | 1049,89 | T6Bifid | L | 1208,43 |
| | T7 | L | 1037,08 | T7 | L | 1081,79 | | T7 | L | 976,33 | T7 | L | 863,90 |
| | T8 | L | 963,64 | T8 | L | 973,90 | | – | – | – | – | – | – |
| | T1 | R | 481,07 | T1 | R | 467,12 | | – | – | – | T1 | R | 574,71 |
| | T2 | R | 565,99 | T2 | R | 690,53 | | T2 | R | 680,92 | T2 | R | 931,56 |
| | T3 | R | 796,69 | T3 | R | 833,48 | | T3 | R | 854,25 | T3 | R | 991,99 |
| | T4 | R | 823,93 | T4 | R | 972,82 | | T4 | R | 1007,24 | T4Bifid | R | 1069,57 |
| | T5 | R | 951,22 | T5 | R | 957,51 | | T5 | R | 1057,50 | T5 | R | 1005,78 |
| | T6 | R | 985,75 | T6 | R | 1087,30 | | T6 | R | 1052,66 | T6 | R | 1050,16 |
| | T7 | R | 1016,08 | T7 | R | 1022,83 | | T7 | R | 942,99 | T7 | R | 885,69 |
| | T8 | R | 975,75 | T8 | R | 974,51 | | – | – | – | – | – | – |

**Table S2. Differentially expressed genes between WT and Sca oral epithelium from early tooth buds in *P. vitticeps* revealed by RNA-seq.** Genes are classified as upregulated (UP) or downregulated (DOWN) based on log2 fold change (log2FC). Columns report gene name, log2 fold change (log2FC), –log10 transformed P-value, raw P-value, false discovery rate (FDR) from the differential expression test. Gene Ontology (GO) biological process annotations are provided only for Developmental process (Dev. Proc.) and Regulation of cell proliferation (Reg. of cell prolif.). The last column indicates genes previously identified as keystone genes according to (2).

| | Name | (log2FC) | (-log10P) | P-Value | FDR | GO (Biological Process) | Keystone gene |
|---|---|---|---|---|---|---|---|
| [UP] | TAMM41 | 0,906 | 1,304 | 0,050 | 0,686 | | |
| [DOWN] | A1CF | -4,415 | 1,813 | 0,015 | 0,469 | Dev. Proc. | |
| [DOWN] | ABAT | -7,625 | 7,146 | 0,000 | 0,000 | | |
| [UP] | ABCA4 | 4,680 | 1,905 | 0,012 | 0,420 | | |
| [DOWN] | ABCC8 | -4,937 | 2,471 | 0,003 | 0,209 | Reg. of cell prolif. | |
| [UP] | ABCG1 | 4,096 | 1,512 | 0,031 | 0,602 | | |
| [UP] | ABHD11 | 4,887 | 2,065 | 0,009 | 0,348 | | |
| [UP] | ABHD12B | 4,532 | 1,776 | 0,017 | 0,485 | | |
| [DOWN] | ABLIM2 | -10,547 | 7,222 | 0,000 | 0,000 | | |
| [DOWN] | ACAT2 | -5,874 | 7,020 | 0,000 | 0,000 | | |
| [DOWN] | ACBD4 | -7,286 | 4,797 | 0,000 | 0,002 | | |
| [DOWN] | ACTN2 | -2,727 | 1,379 | 0,042 | 0,673 | Dev. Proc. | |
| [DOWN] | ACTN3 | -5,556 | 3,237 | 0,001 | 0,053 | Dev. Proc. | |
| [UP] | ADAD1 | 4,425 | 1,769 | 0,017 | 0,485 | Dev. Proc. | |
| [DOWN] | ADAMTS12 | -1,268 | 1,521 | 0,030 | 0,597 | Dev. Proc. | |
| [DOWN] | ADAMTS14 | -1,675 | 1,741 | 0,018 | 0,502 | | |
| [DOWN] | ADAMTS8 | -2,566 | 2,119 | 0,008 | 0,336 | | |
| [UP] | ADAMTSL2 | 4,507 | 1,629 | 0,023 | 0,546 | Dev. Proc. | |
| [DOWN] | ADGRA1 | -1,447 | 1,522 | 0,030 | 0,597 | | |
| [DOWN] | ADGRF5 | -1,087 | 1,413 | 0,039 | 0,653 | Dev. Proc. | |
| [DOWN] | ADGRG6 | -1,000 | 1,542 | 0,029 | 0,589 | Dev. Proc. | |
| [UP] | ADIPOR1 | 1,031 | 1,536 | 0,029 | 0,589 | | |
| [DOWN] | ADORA2B | -4,232 | 1,868 | 0,014 | 0,435 | Reg. of cell prolif. | |
| [UP] | ADTRP | 3,678 | 2,168 | 0,007 | 0,318 | Dev. Proc. | |
| [DOWN] | AFF3 | -2,522 | 4,598 | 0,000 | 0,004 | Dev. Proc. | |
| [DOWN] | AGL | -2,887 | 1,660 | 0,022 | 0,532 | | |
| [DOWN] | AGTR1 | -1,776 | 1,810 | 0,015 | 0,470 | Dev. Proc. + Reg. of cell prolif. | |
| [DOWN] | AGXT | -13,672 | 9,300 | 0,000 | 0,000 | | |
| [UP] | AIFM3 | 4,436 | 1,714 | 0,019 | 0,516 | | |
| [UP] | ALAS2 | 4,919 | 2,152 | 0,007 | 0,325 | Dev. Proc. | |
| [DOWN] | ALDH1A3 | -1,601 | 2,633 | 0,002 | 0,161 | Dev. Proc. | |
| [UP] | ALPK2 | 3,767 | 1,365 | 0,043 | 0,673 | Dev. Proc. | |
| [DOWN] | ALX4 | -4,712 | 2,291 | 0,005 | 0,275 | Dev. Proc. | |
| [UP] | AMBN | 2,750 | 7,063 | 0,000 | 0,000 | Dev. Proc. + Reg. of cell prolif. | X |
| [DOWN] | AMBRA1 | -3,883 | 2,390 | 0,004 | 0,238 | Dev. Proc. + Reg. of cell prolif. | |

| | | | | | | |
|---|---|---|---|---|---|---|
| [DOWN] | ANGPT2 | -4,311 | 2,199 | 0,006 | 0,309 | Dev. Proc. |
| [DOWN] | ANGPTL5 | -3,998 | 2,359 | 0,004 | 0,251 | |
| [DOWN] | ANGPTL7 | -1,686 | 1,458 | 0,035 | 0,637 | Dev. Proc. |
| [DOWN] | ANKMY1 | -4,256 | 1,918 | 0,012 | 0,416 | |
| [DOWN] | ANKRD16 | -3,467 | 2,218 | 0,006 | 0,301 | |
| [DOWN] | ANKRD2 | -3,733 | 1,534 | 0,029 | 0,589 | Dev. Proc. + Reg. of cell prolif. |
| [DOWN] | ANKRD28 | -5,081 | 3,027 | 0,001 | 0,080 | |
| [DOWN] | ANKRD42 | -13,357 | 9,373 | 0,000 | 0,000 | |
| [UP] | ANKRD55 | 4,611 | 1,819 | 0,015 | 0,465 | |
| [DOWN] | ANO3 | -4,232 | 1,868 | 0,014 | 0,435 | |
| [DOWN] | ANPEP | -1,033 | 1,666 | 0,022 | 0,532 | Dev. Proc. |
| [DOWN] | AP4B1 | -1,098 | 1,532 | 0,029 | 0,590 | |
| [UP] | AP4S1 | 1,650 | 2,183 | 0,007 | 0,316 | |
| [DOWN] | APELA | -2,955 | 3,125 | 0,001 | 0,066 | Dev. Proc. + Reg. of cell prolif. |
| [UP] | APOB | 3,930 | 1,361 | 0,044 | 0,677 | Dev. Proc. |
| [DOWN] | APOBEC4 | -3,583 | 1,406 | 0,039 | 0,653 | |
| [DOWN] | APPL2 | -5,971 | 3,745 | 0,000 | 0,021 | Dev. Proc. + Reg. of cell prolif. |
| [UP] | AQP10 | 3,800 | 1,366 | 0,043 | 0,673 | |
| [DOWN] | ARHGAP28 | -6,309 | 1,745 | 0,018 | 0,502 | |
| [UP] | ARMC4 | 4,134 | 1,480 | 0,033 | 0,624 | Dev. Proc. |
| [DOWN] | ARPP21 | -10,072 | 9,290 | 0,000 | 0,000 | |
| [DOWN] | ASB5 | -11,113 | 7,723 | 0,000 | 0,000 | |
| [UP] | ASF1B | 1,264 | 1,825 | 0,015 | 0,463 | Dev. Proc. |
| [UP] | ASGR1 | 3,755 | 1,317 | 0,048 | 0,680 | |
| [DOWN] | ASPA | -8,729 | 8,761 | 0,000 | 0,000 | |
| [DOWN] | ASRGL1 | -4,773 | 4,717 | 0,000 | 0,003 | |
| [DOWN] | ATF3 | -2,266 | 1,420 | 0,038 | 0,653 | Dev. Proc. + Reg. of cell prolif. |
| [DOWN] | ATG4A | -2,771 | 2,766 | 0,002 | 0,131 | |
| [DOWN] | ATP23 | -3,359 | 3,778 | 0,000 | 0,020 | |
| [DOWN] | ATP2C1 | -2,331 | 1,586 | 0,026 | 0,577 | Dev. Proc. |
| [UP] | ATP5G1 | 0,912 | 1,312 | 0,049 | 0,680 | |
| [UP] | ATP5S | 1,580 | 1,538 | 0,029 | 0,589 | |
| [UP] | ATP6V1H | 0,963 | 1,443 | 0,036 | 0,641 | Dev. Proc. |
| [DOWN] | ATP8A1 | -1,196 | 1,418 | 0,038 | 0,653 | |
| [UP] | ATP8A2 | 4,220 | 1,598 | 0,025 | 0,568 | Dev. Proc. |
| [DOWN] | AVIL | -3,135 | 2,100 | 0,008 | 0,343 | Dev. Proc. |
| [UP] | BAHD1 | 1,257 | 1,892 | 0,013 | 0,424 | |
| [UP] | BBS12 | 1,873 | 1,567 | 0,027 | 0,587 | Dev. Proc. |
| [UP] | BGLAP | 1,142 | 1,604 | 0,025 | 0,568 | Dev. Proc. |
| [UP] | BMT2 | 1,239 | 1,938 | 0,012 | 0,416 | |
| [DOWN] | BMX | -11,784 | 8,356 | 0,000 | 0,000 | |
| [DOWN] | BRCA2 | -4,371 | 2,683 | 0,002 | 0,149 | Dev. Proc. + Reg. of cell prolif. |
| [DOWN] | BRWD3 | -3,356 | 2,183 | 0,007 | 0,316 | |

| | | | | | | |
|---|---|---|---|---|---|---|
| [DOWN] | C1QTNF3 | -4,491 | 2,076 | 0,008 | 0,344 | |
| [DOWN] | CA12 | -3,976 | 1,661 | 0,022 | 0,532 | |
| [DOWN] | CABYR | -3,370 | 1,310 | 0,049 | 0,680 | Dev. Proc. |
| [UP] | CACNA2D3 | 4,926 | 2,130 | 0,007 | 0,336 | |
| [DOWN] | CACNB4 | -4,711 | 2,266 | 0,005 | 0,283 | Dev. Proc. + Reg. of cell prolif. |
| [DOWN] | CACNG7 | -3,512 | 1,373 | 0,042 | 0,673 | |
| [DOWN] | CADM2 | -1,048 | 1,624 | 0,024 | 0,550 | |
| [UP] | CAMTA1 | 1,728 | 1,631 | 0,023 | 0,546 | |
| [DOWN] | CARD14 | -1,419 | 1,841 | 0,014 | 0,454 | |
| [DOWN] | CARMIL2 | -3,794 | 1,565 | 0,027 | 0,587 | Dev. Proc. + Reg. of cell prolif. |
| [DOWN] | CARMIL3 | -3,583 | 1,406 | 0,039 | 0,653 | |
| [UP] | CASP7 | 0,940 | 1,334 | 0,046 | 0,680 | Dev. Proc. |
| [UP] | CCDC103 | 2,656 | 1,504 | 0,031 | 0,608 | Dev. Proc. |
| [DOWN] | CCDC129 | -7,766 | 2,771 | 0,002 | 0,130 | |
| [UP] | CCDC181 | 2,579 | 1,643 | 0,023 | 0,544 | |
| [DOWN] | CCDC183 | -3,583 | 1,406 | 0,039 | 0,653 | |
| [DOWN] | CCDC3 | -2,076 | 1,700 | 0,020 | 0,526 | |
| [UP] | CCDC85B | 3,800 | 1,366 | 0,043 | 0,673 | Dev. Proc. |
| [DOWN] | CCDC88B | -2,933 | 1,633 | 0,023 | 0,546 | Reg. of cell prolif. |
| [UP] | CCKBR | 3,912 | 1,414 | 0,039 | 0,653 | Dev. Proc. + Reg. of cell prolif. |
| [UP] | CCM2L | 3,767 | 1,365 | 0,043 | 0,673 | Dev. Proc. |
| [UP] | CCNB1 | 1,627 | 3,196 | 0,001 | 0,057 | Dev. Proc. + Reg. of cell prolif. |
| [UP] | CCNG1 | 2,112 | 1,325 | 0,047 | 0,680 | Dev. Proc. |
| [UP] | CCNJL | 3,904 | 1,448 | 0,036 | 0,637 | |
| [DOWN] | CD96 | -9,241 | 3,046 | 0,001 | 0,077 | |
| [DOWN] | CDC42EP1 | -1,680 | 1,748 | 0,018 | 0,501 | |
| [UP] | CDCA3 | 1,211 | 1,460 | 0,035 | 0,637 | |
| [DOWN] | CDH2 | -3,307 | 1,742 | 0,018 | 0,502 | Dev. Proc. + Reg. of cell prolif. |
| [UP] | CDHR5 | 4,507 | 1,629 | 0,023 | 0,546 | Dev. Proc. |
| [UP] | CDK2AP1 | 1,472 | 1,440 | 0,036 | 0,643 | Dev. Proc. |
| [DOWN] | CDR2L | -6,834 | 4,845 | 0,000 | 0,002 | |
| [DOWN] | CELF4 | -3,370 | 1,310 | 0,049 | 0,680 | Dev. Proc. |
| [UP] | CENPH | 1,267 | 2,127 | 0,007 | 0,336 | |
| [UP] | CENPW | 1,531 | 1,520 | 0,030 | 0,597 | |
| [DOWN] | CER1 | -4,256 | 1,918 | 0,012 | 0,416 | Dev. Proc. + Reg. of cell prolif. |
| [UP] | CFAP58 | 5,181 | 2,451 | 0,004 | 0,216 | Dev. Proc. |
| [UP] | CFAP73 | 4,358 | 1,554 | 0,028 | 0,589 | Dev. Proc. |
| [DOWN] | CFAP99 | -9,053 | 2,664 | 0,002 | 0,155 | |
| [DOWN] | CHAC1 | -2,463 | 1,786 | 0,016 | 0,482 | Dev. Proc. |
| [DOWN] | CHRNA1 | -4,020 | 1,739 | 0,018 | 0,502 | Dev. Proc. |
| [DOWN] | CHTF18 | -1,239 | 1,916 | 0,012 | 0,416 | Dev. Proc. |
| [DOWN] | CIZ1 | -5,407 | 5,088 | 0,000 | 0,001 | |
| [UP] | CKAP2 | 0,890 | 1,332 | 0,047 | 0,680 | |

| | | | | | | |
|---|---|---|---|---|---|---|
| [UP] | CKS2 | 1,207 | 1,437 | 0,037 | 0,645 | Dev. Proc. |
| [DOWN] | CLDN10 | -3,370 | 1,310 | 0,049 | 0,680 | |
| [DOWN] | CLPTM1L | -3,099 | 1,949 | 0,011 | 0,416 | |
| [DOWN] | CMYA5 | -4,668 | 1,895 | 0,013 | 0,423 | |
| [DOWN] | CNMD | -3,794 | 1,565 | 0,027 | 0,587 | Dev. Proc. + Reg. of cell prolif. |
| [DOWN] | CNNM2 | -1,939 | 1,535 | 0,029 | 0,589 | |
| [DOWN] | CNOT2 | -2,273 | 1,669 | 0,021 | 0,532 | Dev. Proc. |
| [DOWN] | CNTN1 | -2,313 | 2,338 | 0,005 | 0,258 | Dev. Proc. |
| [DOWN] | CNTN4 | -4,350 | 3,016 | 0,001 | 0,081 | Dev. Proc. |
| [DOWN] | CNTN5 | -1,316 | 1,670 | 0,021 | 0,532 | Dev. Proc. |
| [DOWN] | CNTNAP2 | -3,855 | 1,596 | 0,025 | 0,568 | Dev. Proc. |
| [DOWN] | COG4 | -4,229 | 3,550 | 0,000 | 0,029 | |
| [DOWN] | COL20A1 | -3,050 | 1,635 | 0,023 | 0,546 | |
| [DOWN] | COL21A1 | -2,623 | 5,260 | 0,000 | 0,001 | |
| [DOWN] | COL4A2 | -1,111 | 1,883 | 0,013 | 0,429 | Dev. Proc. |
| [DOWN] | COQ5 | -1,154 | 1,329 | 0,047 | 0,680 | |
| [DOWN] | CORIN | -1,631 | 2,090 | 0,008 | 0,343 | Dev. Proc. |
| [UP] | CRYGN | 3,956 | 1,417 | 0,038 | 0,653 | Dev. Proc. |
| [DOWN] | CSF1R | -1,335 | 1,888 | 0,013 | 0,427 | Dev. Proc. + Reg. of cell prolif. |
| [UP] | CSRP2 | 0,989 | 1,521 | 0,030 | 0,597 | Dev. Proc. |
| [DOWN] | CTDNEP1 | -1,354 | 1,970 | 0,011 | 0,408 | Dev. Proc. |
| [UP] | CUNH11orf65 | 2,511 | 2,210 | 0,006 | 0,304 | |
| [UP] | CUNH11orf70 | 4,538 | 1,810 | 0,015 | 0,470 | |
| [UP] | CUNH11orf88 | 2,666 | 1,483 | 0,033 | 0,623 | |
| [UP] | CUNH12orf57 | 1,233 | 1,460 | 0,035 | 0,637 | |
| [DOWN] | CUNH14orf132 | -3,976 | 1,661 | 0,022 | 0,532 | |
| [DOWN] | CUNH14orf180 | -3,745 | 2,224 | 0,006 | 0,299 | |
| [DOWN] | CUNH16orf86 | -11,845 | 8,462 | 0,000 | 0,000 | |
| [DOWN] | CUNH17orf67 | -5,060 | 2,616 | 0,002 | 0,166 | |
| [UP] | CUNH18orf32 | 1,175 | 1,661 | 0,022 | 0,532 | |
| [UP] | CUNH21orf58 | 2,320 | 1,410 | 0,039 | 0,653 | |
| [UP] | CUNH2orf81 | 2,649 | 1,547 | 0,028 | 0,589 | |
| [DOWN] | CUNH4orf22 | -3,916 | 1,629 | 0,024 | 0,546 | |
| [UP] | CUNH5orf63 | 2,145 | 2,973 | 0,001 | 0,088 | |
| [UP] | CUNH8orf59 | 1,517 | 2,024 | 0,009 | 0,374 | |
| [UP] | CUNH8orf89 | 5,090 | 2,303 | 0,005 | 0,275 | |
| [DOWN] | CUNHXorf38 | -1,454 | 1,771 | 0,017 | 0,485 | |
| [DOWN] | CUNHXorf65 | -3,370 | 1,310 | 0,049 | 0,680 | |
| [DOWN] | CYGB | -0,892 | 1,401 | 0,040 | 0,657 | |
| [DOWN] | CYTIP | -2,553 | 1,791 | 0,016 | 0,479 | |
| [UP] | DALRD3 | 1,209 | 1,310 | 0,049 | 0,680 | |
| [UP] | DAPL1 | 1,138 | 1,746 | 0,018 | 0,501 | Dev. Proc. |
| [DOWN] | DCUN1D3 | -1,354 | 1,915 | 0,012 | 0,416 | |

| | | | | | | | |
|---|---|---|---|---|---|---|---|
| [UP] | DDIAS | 1,078 | 1,551 | 0,028 | 0,589 | Dev. Proc. | |
| [DOWN] | DDX4 | -3,139 | 1,503 | 0,031 | 0,608 | Dev. Proc. | |
| [DOWN] | DDX58 | -2,691 | 3,591 | 0,000 | 0,027 | | |
| [UP] | DECR1 | 1,729 | 3,606 | 0,000 | 0,027 | | |
| [UP] | DENND5B | 0,940 | 1,379 | 0,042 | 0,673 | | |
| [DOWN] | DGKE | -3,100 | 3,036 | 0,001 | 0,078 | | |
| [DOWN] | DGKZ | -0,917 | 1,373 | 0,042 | 0,673 | | |
| [UP] | DHCR24 | 0,951 | 1,441 | 0,036 | 0,642 | Dev. Proc. + Reg. of cell prolif. | |
| [UP] | DHCR7 | 1,057 | 1,483 | 0,033 | 0,623 | Dev. Proc. + Reg. of cell prolif. | |
| [DOWN] | DHX58 | -2,914 | 4,789 | 0,000 | 0,002 | | |
| [DOWN] | DIO3 | -3,583 | 1,848 | 0,014 | 0,448 | Dev. Proc. | |
| [DOWN] | DIXDC1 | -2,522 | 5,754 | 0,000 | 0,000 | Dev. Proc. | |
| [DOWN] | DKK2 | -0,901 | 1,431 | 0,037 | 0,650 | Dev. Proc. | |
| [DOWN] | DKKL1 | -3,383 | 1,752 | 0,018 | 0,499 | | |
| [DOWN] | DLC1 | -0,997 | 1,492 | 0,032 | 0,616 | Dev. Proc. + Reg. of cell prolif. | |
| [DOWN] | DLK1 | -2,022 | 1,314 | 0,049 | 0,680 | Dev. Proc. + Reg. of cell prolif. | |
| [UP] | DLX6 | 2,751 | 1,924 | 0,012 | 0,416 | Dev. Proc. + Reg. of cell prolif. | X |
| [UP] | DMC1 | 3,942 | 1,454 | 0,035 | 0,637 | Dev. Proc. | |
| [DOWN] | DNAH11 | -4,611 | 1,487 | 0,033 | 0,621 | Dev. Proc. | |
| [DOWN] | DNAJB4 | -2,768 | 2,587 | 0,003 | 0,173 | | |
| [DOWN] | DNAJC6 | -11,348 | 7,715 | 0,000 | 0,000 | | |
| [DOWN] | DNHD1 | -7,979 | 5,607 | 0,000 | 0,000 | Dev. Proc. | |
| [UP] | DNTT | 2,768 | 1,585 | 0,026 | 0,577 | | |
| [UP] | DOK2 | 4,067 | 1,535 | 0,029 | 0,589 | | |
| [UP] | DPH2 | 1,444 | 1,335 | 0,046 | 0,680 | | |
| [DOWN] | DPM3 | -3,033 | 1,749 | 0,018 | 0,501 | | |
| [UP] | DPP3 | 1,547 | 1,903 | 0,012 | 0,420 | | |
| [UP] | DPYS | 3,930 | 1,361 | 0,044 | 0,677 | | |
| [DOWN] | DSCAM | -3,023 | 1,407 | 0,039 | 0,653 | Dev. Proc. | |
| [UP] | DUSP11 | 1,104 | 1,448 | 0,036 | 0,637 | | |
| [UP] | DUSP27 | 3,573 | 2,168 | 0,007 | 0,318 | | |
| [DOWN] | DUSP5 | -2,675 | 1,651 | 0,022 | 0,539 | Dev. Proc. | |
| [DOWN] | ECT2L | -8,661 | 6,380 | 0,000 | 0,000 | | |
| [DOWN] | EDEM3 | -4,371 | 2,773 | 0,002 | 0,130 | | |
| [DOWN] | EFHC2 | -3,370 | 1,310 | 0,049 | 0,680 | | |
| [UP] | EGFL6 | 3,800 | 1,366 | 0,043 | 0,673 | Dev. Proc. | |
| [DOWN] | EGFL7 | -2,482 | 1,512 | 0,031 | 0,602 | Dev. Proc. + Reg. of cell prolif. | |
| [UP] | EGR1 | 1,032 | 1,427 | 0,037 | 0,653 | Dev. Proc. + Reg. of cell prolif. | |
| [DOWN] | ELL | -1,194 | 1,428 | 0,037 | 0,653 | Dev. Proc. | |
| [DOWN] | ELN | -1,030 | 1,642 | 0,023 | 0,545 | Dev. Proc. | |
| [UP] | ELOVL7 | 2,290 | 1,372 | 0,042 | 0,673 | | |
| [UP] | EMC2 | 1,058 | 1,626 | 0,024 | 0,548 | | |
| [DOWN] | EMILIN1 | -1,104 | 1,383 | 0,041 | 0,673 | Dev. Proc. + Reg. of cell prolif. | |

| | | | | | | |
|---|---|---|---|---|---|---|
| [DOWN] | ENPP3 | -1,572 | 1,450 | 0,035 | 0,637 | Reg. of cell prolif. |
| [UP] | ENPP5 | 0,991 | 1,553 | 0,028 | 0,589 | |
| [UP] | ENTPD1 | 2,549 | 1,339 | 0,046 | 0,680 | |
| [DOWN] | ERFE | -3,583 | 1,406 | 0,039 | 0,653 | |
| [DOWN] | ERICH2 | -6,687 | 5,144 | 0,000 | 0,001 | |
| [DOWN] | ERICH5 | -5,313 | 2,877 | 0,001 | 0,106 | |
| [UP] | ESPL1 | 1,741 | 1,724 | 0,019 | 0,511 | |
| [UP] | ETFBKMT | 1,616 | 1,361 | 0,044 | 0,677 | |
| [DOWN] | EVL | -4,675 | 3,833 | 0,000 | 0,018 | Dev. Proc. |
| [DOWN] | EXO1 | -1,380 | 1,332 | 0,047 | 0,680 | Dev. Proc. |
| [DOWN] | EXOC3L2 | -4,127 | 1,802 | 0,016 | 0,474 | Dev. Proc. |
| [DOWN] | F8 | -5,162 | 1,603 | 0,025 | 0,568 | |
| [UP] | FAAP24 | 1,712 | 2,509 | 0,003 | 0,196 | |
| [DOWN] | FAIM2 | -3,583 | 1,406 | 0,039 | 0,653 | Dev. Proc. |
| [UP] | FAM104A | 1,315 | 1,694 | 0,020 | 0,530 | |
| [DOWN] | FAM110D | -3,733 | 1,534 | 0,029 | 0,589 | |
| [UP] | FAM124A | 2,968 | 1,412 | 0,039 | 0,653 | |
| [DOWN] | FAM124B | -4,232 | 1,868 | 0,014 | 0,435 | |
| [UP] | FAM129A | 1,181 | 1,945 | 0,011 | 0,416 | |
| [DOWN] | FAM160A1 | -3,189 | 1,877 | 0,013 | 0,432 | |
| [DOWN] | FAM161A | -8,375 | 5,795 | 0,000 | 0,000 | |
| [DOWN] | FAM161B | -3,706 | 4,031 | 0,000 | 0,012 | |
| [UP] | FAM167A | 1,063 | 1,508 | 0,031 | 0,604 | |
| [UP] | FAM169A | 1,764 | 1,303 | 0,050 | 0,686 | |
| [UP] | FAM173B | 1,287 | 1,699 | 0,020 | 0,526 | |
| [DOWN] | FAM196A | -1,232 | 1,343 | 0,045 | 0,680 | |
| [DOWN] | FAM217A | -3,733 | 1,534 | 0,029 | 0,589 | |
| [DOWN] | FAM3D | -3,312 | 1,805 | 0,016 | 0,474 | |
| [DOWN] | FAM46C | -8,641 | 6,311 | 0,000 | 0,000 | |
| [UP] | FAM53A | 1,459 | 2,548 | 0,003 | 0,183 | |
| [DOWN] | FAM69B | -4,074 | 1,770 | 0,017 | 0,485 | |
| [UP] | FAM83D | 1,651 | 1,713 | 0,019 | 0,517 | Dev. Proc. |
| [DOWN] | FAM89A | -1,212 | 1,660 | 0,022 | 0,532 | |
| [DOWN] | FAN1 | -3,841 | 1,821 | 0,015 | 0,464 | |
| [DOWN] | FBXO43 | -8,946 | 3,765 | 0,000 | 0,020 | |
| [UP] | FDXACB1 | 1,740 | 1,646 | 0,023 | 0,543 | |
| [DOWN] | FERMT2 | -0,928 | 1,458 | 0,035 | 0,637 | Dev. Proc. + Reg. of cell prolif. |
| [UP] | FGF13 | 3,767 | 1,365 | 0,043 | 0,673 | Dev. Proc. |
| [DOWN] | FGF18 | -2,888 | 1,734 | 0,018 | 0,504 | Dev. Proc. + Reg. of cell prolif. |
| [DOWN] | FGF9 | -1,180 | 1,471 | 0,034 | 0,632 | Dev. Proc. + Reg. of cell prolif. |
| [UP] | FGG | 3,904 | 1,448 | 0,036 | 0,637 | |
| [UP] | FGGY | 1,149 | 1,963 | 0,011 | 0,412 | |
| [UP] | FHL1 | 0,906 | 1,360 | 0,044 | 0,677 | Dev. Proc. |

| | | | | | | | |
|---|---|---|---|---|---|---|---|
| [UP] | FHL3 | 1,696 | 1,334 | 0,046 | 0,680 | | |
| [DOWN] | FIBIN | -4,463 | 2,646 | 0,002 | 0,158 | | |
| [DOWN] | FKBP6 | -3,441 | 1,341 | 0,046 | 0,680 | Dev. Proc. | |
| [DOWN] | FMOD | -1,722 | 2,220 | 0,006 | 0,300 | | |
| [DOWN] | FNDC7 | -3,916 | 1,629 | 0,024 | 0,546 | | |
| [DOWN] | FOXA1 | -4,256 | 1,918 | 0,012 | 0,416 | Dev. Proc. | |
| [DOWN] | FOXI2 | -2,332 | 1,651 | 0,022 | 0,539 | Dev. Proc. | |
| [UP] | FOXRED2 | 1,389 | 1,736 | 0,018 | 0,504 | | |
| [UP] | FPGT | 1,082 | 1,633 | 0,023 | 0,546 | | |
| [DOWN] | FST | -0,923 | 1,354 | 0,044 | 0,680 | Dev. Proc. | X |
| [DOWN] | FTCD | -13,245 | 9,453 | 0,000 | 0,000 | | |
| [UP] | FUCA2 | 1,155 | 1,602 | 0,025 | 0,568 | | |
| [DOWN] | FUT9 | -1,124 | 1,922 | 0,012 | 0,416 | Dev. Proc. | |
| [DOWN] | GABRG1 | -5,601 | 2,221 | 0,006 | 0,300 | Dev. Proc. | |
| [DOWN] | GABRP | -3,370 | 1,310 | 0,049 | 0,680 | | |
| [UP] | GALK1 | 2,115 | 1,819 | 0,015 | 0,465 | | |
| [DOWN] | GALNT15 | -3,092 | 1,467 | 0,034 | 0,635 | Dev. Proc. | |
| [DOWN] | GALNT6 | -2,759 | 1,678 | 0,021 | 0,532 | | |
| [DOWN] | GATA2 | -4,352 | 1,980 | 0,010 | 0,399 | Dev. Proc. + Reg. of cell prolif. | |
| [UP] | GATA3 | 4,323 | 1,655 | 0,022 | 0,536 | Dev. Proc. + Reg. of cell prolif. | |
| [DOWN] | GCK | -3,976 | 1,661 | 0,022 | 0,532 | | |
| [DOWN] | GCNT7 | -4,433 | 4,382 | 0,000 | 0,006 | | |
| [DOWN] | GDNF | -3,794 | 1,565 | 0,027 | 0,587 | Dev. Proc. + Reg. of cell prolif. | X |
| [DOWN] | GDPD3 | -5,198 | 2,771 | 0,002 | 0,130 | | |
| [UP] | GLB1L | 2,115 | 1,335 | 0,046 | 0,680 | | |
| [UP] | GLE1 | 1,688 | 1,721 | 0,019 | 0,511 | | |
| [DOWN] | GLIS3 | -2,026 | 1,390 | 0,041 | 0,668 | | |
| [DOWN] | GNAO1 | -3,321 | 1,688 | 0,021 | 0,532 | | |
| [DOWN] | GP1BA | -4,217 | 1,463 | 0,034 | 0,637 | Dev. Proc. | |
| [DOWN] | GPA33 | -7,751 | 7,675 | 0,000 | 0,000 | | |
| [DOWN] | GPBAR1 | -4,256 | 1,918 | 0,012 | 0,416 | Regulation of cell proliferation | |
| [DOWN] | GPM6A | -1,048 | 1,664 | 0,022 | 0,532 | Dev. Proc. | |
| [DOWN] | GPNMB | -2,173 | 3,056 | 0,001 | 0,076 | Regulation of cell proliferation | |
| [UP] | GPR137 | 3,322 | 1,690 | 0,020 | 0,532 | | |
| [DOWN] | GPR149 | -3,733 | 1,534 | 0,029 | 0,589 | Dev. Proc. | |
| [DOWN] | GRB10 | -4,133 | 2,624 | 0,002 | 0,164 | | |
| [DOWN] | GREM2 | -1,379 | 2,043 | 0,009 | 0,362 | Dev. Proc. | X |
| [DOWN] | GRK5 | -3,460 | 5,618 | 0,000 | 0,000 | Dev. Proc. + Reg. of cell prolif. | |
| [DOWN] | GRM8 | -3,733 | 1,534 | 0,029 | 0,589 | | |
| [DOWN] | HACD1 | -3,442 | 1,783 | 0,016 | 0,482 | Dev. Proc. | |
| [UP] | HACD3 | 0,908 | 1,324 | 0,047 | 0,680 | | |
| [DOWN] | HAL | -13,415 | 9,332 | 0,000 | 0,000 | | |
| [UP] | HAS3 | 4,473 | 1,788 | 0,016 | 0,481 | | |

| | | | | | | |
|---|---|---|---|---|---|---|
| [DOWN] | HCN1 | -9,458 | 9,105 | 0,000 | 0,000 | Dev. Proc. |
| [DOWN] | HEATR9 | -12,953 | 9,671 | 0,000 | 0,000 | Dev. Proc. |
| [UP] | HEBP1 | 1,362 | 2,075 | 0,008 | 0,344 | |
| [DOWN] | HELZ2 | -1,280 | 2,028 | 0,009 | 0,371 | |
| [DOWN] | HHIPL1 | -3,142 | 4,226 | 0,000 | 0,008 | |
| [UP] | HMGCS1 | 1,068 | 1,521 | 0,030 | 0,597 | |
| [DOWN] | HOXD3 | -3,370 | 1,310 | 0,049 | 0,680 | Dev. Proc. |
| [DOWN] | HOXD8 | -3,733 | 1,534 | 0,029 | 0,589 | Dev. Proc. |
| [UP] | HS3ST3B1 | 5,645 | 2,752 | 0,002 | 0,134 | Dev. Proc. |
| [DOWN] | HTR4 | -2,637 | 3,783 | 0,000 | 0,020 | |
| [UP] | IDH3G | 1,004 | 1,537 | 0,029 | 0,589 | |
| [DOWN] | IFIH1 | -2,044 | 1,370 | 0,043 | 0,673 | |
| [UP] | IFT140 | 0,999 | 1,468 | 0,034 | 0,635 | Dev. Proc. |
| [DOWN] | IFT88 | -2,772 | 4,831 | 0,000 | 0,002 | Dev. Proc. + Reg. of cell prolif. |
| [DOWN] | IGFN1 | -8,659 | 6,579 | 0,000 | 0,000 | |
| [DOWN] | IGSF10 | -1,267 | 1,914 | 0,012 | 0,417 | Dev. Proc. |
| [DOWN] | IGSF11 | -3,794 | 1,565 | 0,027 | 0,587 | |
| [UP] | IL10RB | 1,683 | 1,805 | 0,016 | 0,474 | |
| [DOWN] | IL17D | -5,068 | 2,604 | 0,002 | 0,169 | |
| [UP] | IL18R1 | 5,288 | 2,455 | 0,004 | 0,215 | Dev. Proc. |
| [DOWN] | IL1RAPL1 | -3,960 | 2,651 | 0,002 | 0,158 | Dev. Proc. |
| [UP] | IL6R | 3,800 | 1,366 | 0,043 | 0,673 | Dev. Proc. + Reg. of cell prolif. |
| [DOWN] | IMPG1 | -3,441 | 1,341 | 0,046 | 0,680 | |
| [UP] | ING4 | 1,164 | 1,325 | 0,047 | 0,680 | Reg. of cell prolif. |
| [DOWN] | INPP5B | -2,421 | 1,516 | 0,030 | 0,598 | Dev. Proc. |
| [DOWN] | INPP5D | -1,878 | 2,459 | 0,003 | 0,213 | Reg. of cell prolif. |
| [UP] | IQCA1 | 3,904 | 1,448 | 0,036 | 0,637 | |
| [UP] | IRAK4 | 1,448 | 2,152 | 0,007 | 0,325 | Reg. of cell prolif. |
| [UP] | IRX6 | 3,767 | 1,365 | 0,043 | 0,673 | Dev. Proc. |
| [DOWN] | ISL2 | -4,457 | 2,081 | 0,008 | 0,343 | Dev. Proc. |
| [DOWN] | ITGA4 | -2,266 | 4,595 | 0,000 | 0,004 | Dev. Proc. + Reg. of cell prolif. |
| [DOWN] | ITGB2 | -6,798 | 4,399 | 0,000 | 0,005 | |
| [DOWN] | ITM2C | -3,253 | 1,567 | 0,027 | 0,587 | Dev. Proc. |
| [DOWN] | JCHAIN | -3,794 | 1,565 | 0,027 | 0,587 | |
| [DOWN] | JOSD2 | -5,657 | 4,035 | 0,000 | 0,012 | |
| [DOWN] | KCNA4 | -4,937 | 2,471 | 0,003 | 0,209 | |
| [UP] | KCNAB1 | 1,944 | 1,703 | 0,020 | 0,525 | |
| [UP] | KCNB1 | 3,904 | 1,448 | 0,036 | 0,637 | Dev. Proc. |
| [DOWN] | KCND2 | -12,272 | 9,387 | 0,000 | 0,000 | |
| [DOWN] | KCNE5 | -4,283 | 1,901 | 0,013 | 0,420 | |
| [UP] | KCNG1 | 3,483 | 1,951 | 0,011 | 0,416 | |
| [DOWN] | KCNG4 | -4,217 | 1,463 | 0,034 | 0,637 | |
| [UP] | KCNH7 | 4,358 | 1,554 | 0,028 | 0,589 | |

| | | | | | | | |
|---|---|---|---|---|---|---|---|
| [DOWN] | KCNJ15 | -1,759 | 1,455 | 0,035 | 0,637 | | |
| [DOWN] | KCNK15 | -1,836 | 1,563 | 0,027 | 0,589 | | |
| [DOWN] | KCNS3 | -2,583 | 1,307 | 0,049 | 0,684 | | |
| [UP] | KCNT2 | 4,027 | 1,466 | 0,034 | 0,635 | | |
| [DOWN] | KCTD12 | -1,175 | 1,838 | 0,015 | 0,456 | | |
| [DOWN] | KCTD17 | -1,976 | 1,844 | 0,014 | 0,452 | | |
| [UP] | KCTD8 | 3,982 | 1,459 | 0,035 | 0,637 | | |
| [DOWN] | KDM7A | -4,110 | 2,585 | 0,003 | 0,174 | Dev. Proc. | |
| [DOWN] | KEL | -2,443 | 2,545 | 0,003 | 0,183 | Dev. Proc. | |
| [DOWN] | KIAA1324L | -4,927 | 2,503 | 0,003 | 0,198 | | |
| [DOWN] | KIAA1644 | -1,507 | 2,361 | 0,004 | 0,251 | | |
| [DOWN] | KIAA1755 | -0,938 | 1,359 | 0,044 | 0,678 | | |
| [DOWN] | KIAA1958 | -1,520 | 1,542 | 0,029 | 0,589 | | |
| [UP] | KIAA2022 | 2,019 | 2,654 | 0,002 | 0,158 | | |
| [DOWN] | KIF21A | -2,955 | 2,011 | 0,010 | 0,380 | | |
| [UP] | KIF23 | 0,991 | 1,450 | 0,035 | 0,637 | | |
| [DOWN] | KIN | -7,886 | 5,407 | 0,000 | 0,001 | | |
| [DOWN] | KL | -3,512 | 1,373 | 0,042 | 0,673 | | |
| [DOWN] | KLF10 | -2,425 | 1,663 | 0,022 | 0,532 | Dev. Proc. | |
| [DOWN] | KLF15 | -1,267 | 1,866 | 0,014 | 0,435 | Dev. Proc. | |
| [DOWN] | KLHL14 | -1,794 | 1,784 | 0,016 | 0,482 | | |
| [DOWN] | KLHL4 | -5,991 | 4,023 | 0,000 | 0,012 | | |
| [DOWN] | KMT2A | -4,101 | 3,745 | 0,000 | 0,021 | Dev. Proc. + Reg. of cell prolif. | |
| [DOWN] | KPNA1 | -2,734 | 1,655 | 0,022 | 0,536 | Dev. Proc. | |
| [DOWN] | KRT18 | -1,073 | 1,782 | 0,017 | 0,482 | | |
| [DOWN] | LASP1 | -3,472 | 1,927 | 0,012 | 0,416 | | |
| [UP] | LCTL | 1,566 | 2,547 | 0,003 | 0,183 | Dev. Proc. | |
| [DOWN] | LDLRAD2 | -3,583 | 1,406 | 0,039 | 0,653 | | |
| [DOWN] | LEPR | -1,594 | 2,687 | 0,002 | 0,149 | Dev. Proc. | |
| [DOWN] | LGR5 | -1,357 | 1,919 | 0,012 | 0,416 | Dev. Proc. + Reg. of cell prolif. | |
| [DOWN] | LHX8 | -1,228 | 1,830 | 0,015 | 0,461 | Dev. Proc. | X |
| [UP] | LIMCH1 | 1,412 | 2,613 | 0,002 | 0,167 | | |
| [DOWN] | LIN7A | -6,320 | 6,770 | 0,000 | 0,000 | Dev. Proc. | |
| [DOWN] | LIPI | -8,088 | 5,420 | 0,000 | 0,001 | | |
| [UP] | LMCD1 | 1,146 | 1,621 | 0,024 | 0,553 | | |
| [UP] | LMF2 | 1,210 | 1,537 | 0,029 | 0,589 | | |
| [DOWN] | LMOD1 | -2,592 | 2,337 | 0,005 | 0,258 | Dev. Proc. | |
| [DOWN] | LMX1B | -3,733 | 1,534 | 0,029 | 0,589 | Dev. Proc. | |
| [DOWN] | LPL | -1,109 | 1,678 | 0,021 | 0,532 | | |
| [UP] | LRRC39 | 4,425 | 1,798 | 0,016 | 0,476 | | |
| [DOWN] | LRRC71 | -3,370 | 1,310 | 0,049 | 0,680 | | |
| [DOWN] | LRRC74B | -4,711 | 2,266 | 0,005 | 0,283 | | |
| [UP] | LRRC8D | 1,247 | 1,586 | 0,026 | 0,577 | | |

| | | | | | | | |
|---|---|---|---|---|---|---|---|
| [DOWN] | LRRIQ3 | -8,698 | 2,888 | 0,001 | 0,104 | | |
| [UP] | LSM7 | 1,009 | 1,602 | 0,025 | 0,568 | | |
| [DOWN] | LVRN | -1,217 | 1,715 | 0,019 | 0,516 | | |
| [DOWN] | LYVE1 | -2,088 | 1,727 | 0,019 | 0,510 | | |
| [DOWN] | MAEL | -3,438 | 1,967 | 0,011 | 0,410 | Dev. Proc. | |
| [DOWN] | MAJIN | -3,916 | 1,629 | 0,024 | 0,546 | Dev. Proc. | |
| [DOWN] | MAL | -3,115 | 1,391 | 0,041 | 0,668 | Dev. Proc. + Reg. of cell prolif. | |
| [DOWN] | MAN1A1 | -4,911 | 2,974 | 0,001 | 0,088 | | |
| [DOWN] | MAN2A1 | -3,696 | 2,057 | 0,009 | 0,353 | Dev. Proc. | |
| [DOWN] | MANSC4 | -4,587 | 2,173 | 0,007 | 0,318 | Dev. Proc. | |
| [DOWN] | MAP2K5 | -5,584 | 3,666 | 0,000 | 0,024 | Dev. Proc. + Reg. of cell prolif. | |
| [UP] | MAPK1IP1L | 0,995 | 1,414 | 0,039 | 0,653 | | |
| [UP] | MAPK8 | 0,883 | 1,313 | 0,049 | 0,680 | Dev. Proc. + Reg. of cell prolif. | |
| [DOWN] | MARCH1 | -6,413 | 8,401 | 0,000 | 0,000 | | |
| [DOWN] | MARCH11 | -3,512 | 1,373 | 0,042 | 0,673 | | |
| [UP] | MARK4 | 1,355 | 1,437 | 0,037 | 0,645 | Dev. Proc. | |
| [DOWN] | MB21D1 | -1,955 | 2,635 | 0,002 | 0,161 | | |
| [UP] | MED26 | 2,304 | 1,701 | 0,020 | 0,526 | | |
| [DOWN] | MEIS3 | -3,370 | 1,310 | 0,049 | 0,680 | Dev. Proc. + Reg. of cell prolif. | |
| [UP] | MEST | 1,283 | 1,470 | 0,034 | 0,633 | | |
| [DOWN] | METTL8 | -1,298 | 1,536 | 0,029 | 0,589 | Dev. Proc. | |
| [DOWN] | MFAP4 | -1,050 | 1,711 | 0,019 | 0,517 | | |
| [UP] | MFGE8 | 0,935 | 1,341 | 0,046 | 0,680 | Dev. Proc. | |
| [DOWN] | MGAT1 | -5,592 | 3,628 | 0,000 | 0,026 | Dev. Proc. | |
| [DOWN] | MIB2 | -4,556 | 7,496 | 0,000 | 0,000 | | |
| [DOWN] | MICAL1 | -1,305 | 1,665 | 0,022 | 0,532 | | |
| [DOWN] | MIS18BP1 | -3,602 | 1,520 | 0,030 | 0,597 | | |
| [DOWN] | MLH1 | -1,742 | 2,558 | 0,003 | 0,181 | Dev. Proc. | |
| [DOWN] | MMD | -3,377 | 2,441 | 0,004 | 0,219 | | |
| [UP] | MMP20 | 7,094 | 1,732 | 0,019 | 0,505 | Dev. Proc. | X |
| [DOWN] | MMRN2 | -8,327 | 5,547 | 0,000 | 0,001 | Dev. Proc. + Reg. of cell prolif. | |
| [UP] | MOCOS | 1,657 | 1,670 | 0,021 | 0,532 | | |
| [DOWN] | MORN3 | -10,338 | 9,920 | 0,000 | 0,000 | | |
| [UP] | MPG | 4,899 | 2,128 | 0,007 | 0,336 | | |
| [UP] | MPI | 1,604 | 2,384 | 0,004 | 0,240 | | |
| [UP] | MPPED1 | 1,504 | 2,839 | 0,001 | 0,115 | | |
| [UP] | MRPL21 | 1,042 | 1,404 | 0,039 | 0,655 | | |
| [UP] | MRPL38 | 0,898 | 1,311 | 0,049 | 0,680 | | |
| [UP] | MRPL54 | 1,278 | 2,086 | 0,008 | 0,343 | | |
| [UP] | MRPL57 | 0,908 | 1,335 | 0,046 | 0,680 | | |
| [UP] | MRPS6 | 1,093 | 1,567 | 0,027 | 0,587 | | |
| [DOWN] | MTERF3 | -4,799 | 4,699 | 0,000 | 0,003 | | |
| [DOWN] | MTUS2 | -6,243 | 9,206 | 0,000 | 0,000 | | |

| | | | | | | |
|---|---|---|---|---|---|---|
| [DOWN] | MUC6 | -4,713 | 1,840 | 0,014 | 0,455 | |
| [DOWN] | MXI1 | -0,943 | 1,425 | 0,038 | 0,653 | Dev. Proc. |
| [UP] | MYBPC3 | 4,971 | 2,095 | 0,008 | 0,343 | Dev. Proc. |
| [UP] | MYLK4 | 1,632 | 1,433 | 0,037 | 0,648 | |
| [DOWN] | MYO15B | -3,441 | 1,341 | 0,046 | 0,680 | |
| [UP] | MYO5B | 1,831 | 3,623 | 0,000 | 0,026 | Dev. Proc. |
| [DOWN] | MYO5C | -1,367 | 1,600 | 0,025 | 0,568 | |
| [DOWN] | MYRFL | -4,304 | 1,949 | 0,011 | 0,416 | |
| [UP] | NADK | 0,994 | 1,535 | 0,029 | 0,589 | |
| [UP] | NALCN | 3,464 | 1,867 | 0,014 | 0,435 | |
| [DOWN] | NAV3 | -1,127 | 1,898 | 0,013 | 0,422 | Dev. Proc. |
| [DOWN] | NBEAL2 | -1,643 | 2,646 | 0,002 | 0,158 | Dev. Proc. |
| [UP] | NCAPG2 | 1,021 | 1,346 | 0,045 | 0,680 | Dev. Proc. |
| [DOWN] | NCBP2 | -1,288 | 1,765 | 0,017 | 0,488 | |
| [UP] | NDUFA6 | 1,140 | 1,356 | 0,044 | 0,680 | |
| [UP] | NDUFB1 | 1,103 | 1,605 | 0,025 | 0,568 | |
| [UP] | NDUFV3 | 1,205 | 1,981 | 0,010 | 0,399 | |
| [UP] | NEIL3 | 1,547 | 1,571 | 0,027 | 0,587 | |
| [UP] | NEK2 | 1,244 | 1,497 | 0,032 | 0,614 | Dev. Proc. |
| [DOWN] | NELL2 | -11,486 | 13,311 | 0,000 | 0,000 | Dev. Proc. |
| [DOWN] | NEURL1 | -3,916 | 1,629 | 0,024 | 0,546 | Dev. Proc. + Reg. of cell prolif. |
| [DOWN] | NFE2 | -4,751 | 2,321 | 0,005 | 0,266 | Dev. Proc. + Reg. of cell prolif. |
| [DOWN] | NFKB2 | -2,240 | 3,091 | 0,001 | 0,071 | Dev. Proc. |
| [DOWN] | NGF | -4,861 | 2,430 | 0,004 | 0,222 | Dev. Proc. + Reg. of cell prolif. |
| [UP] | NICN1 | 1,246 | 1,377 | 0,042 | 0,673 | |
| [DOWN] | NLGN3 | -4,457 | 2,081 | 0,008 | 0,343 | Dev. Proc. |
| [DOWN] | NMBR | -3,855 | 1,596 | 0,025 | 0,568 | Reg. of cell prolif. |
| [UP] | NME7 | 2,109 | 2,125 | 0,007 | 0,336 | Dev. Proc. |
| [UP] | NOP10 | 1,192 | 1,320 | 0,048 | 0,680 | |
| [UP] | NOP9 | 1,587 | 1,641 | 0,023 | 0,545 | |
| [UP] | NOS3 | 4,211 | 1,485 | 0,033 | 0,622 | Dev. Proc. + Reg. of cell prolif. |
| [UP] | NOSIP | 1,130 | 1,756 | 0,018 | 0,496 | Dev. Proc. |
| [DOWN] | NOTUM | -2,365 | 2,498 | 0,003 | 0,200 | Dev. Proc. |
| [DOWN] | NPEPPS | -4,690 | 2,743 | 0,002 | 0,136 | |
| [DOWN] | NPLOC4 | -3,066 | 1,821 | 0,015 | 0,464 | |
| [DOWN] | NR0B1 | -3,370 | 1,310 | 0,049 | 0,680 | Dev. Proc. |
| [DOWN] | NR2C2AP | -2,278 | 1,403 | 0,040 | 0,655 | |
| [DOWN] | NR4A2 | -4,232 | 1,868 | 0,014 | 0,435 | Dev. Proc. |
| [DOWN] | NT5DC4 | -3,583 | 1,406 | 0,039 | 0,653 | |
| [DOWN] | NT5E | -1,865 | 2,270 | 0,005 | 0,283 | |
| [DOWN] | NTN4 | -10,814 | 7,606 | 0,000 | 0,000 | Dev. Proc. |
| [DOWN] | NTNG1 | -0,932 | 1,319 | 0,048 | 0,680 | Dev. Proc. |
| [DOWN] | NUP160 | -1,926 | 1,434 | 0,037 | 0,647 | Dev. Proc. |

| | | | | | | |
|---|---|---|---|---|---|---|
| [UP] | NUP37 | 1,345 | 1,495 | 0,032 | 0,615 | |
| [DOWN] | NXNL1 | -10,135 | 1,304 | 0,050 | 0,686 | |
| [DOWN] | NXPH2 | -2,826 | 1,797 | 0,016 | 0,476 | |
| [DOWN] | OBSL1 | -10,477 | 9,451 | 0,000 | 0,000 | |
| [UP] | OCIAD2 | 1,119 | 1,515 | 0,031 | 0,599 | |
| [DOWN] | OCSTAMP | -4,537 | 2,108 | 0,008 | 0,340 | Dev. Proc. + Reg. of cell prolif. |
| [DOWN] | OLIG1 | -2,986 | 1,557 | 0,028 | 0,589 | Dev. Proc. |
| [DOWN] | OMG | -4,446 | 2,044 | 0,009 | 0,362 | Dev. Proc. |
| [UP] | OSBPL6 | 1,217 | 1,546 | 0,028 | 0,589 | |
| [DOWN] | OSBPL8 | -4,198 | 2,459 | 0,003 | 0,213 | Dev. Proc. |
| [UP] | OST4 | 3,942 | 1,454 | 0,035 | 0,637 | |
| [DOWN] | OTOG | -9,997 | 7,201 | 0,000 | 0,000 | Dev. Proc. |
| [DOWN] | OTUD7A | -2,698 | 1,325 | 0,047 | 0,680 | |
| [DOWN] | P2RX1 | -4,632 | 2,232 | 0,006 | 0,296 | |
| [UP] | PACRGL | 2,502 | 1,826 | 0,015 | 0,463 | |
| [DOWN] | PARG | -4,224 | 2,549 | 0,003 | 0,183 | |
| [DOWN] | PARP4 | -3,339 | 2,148 | 0,007 | 0,327 | |
| [DOWN] | PATL2 | -3,976 | 2,367 | 0,004 | 0,248 | |
| [DOWN] | PAXBP1 | -2,085 | 1,485 | 0,033 | 0,622 | Dev. Proc. + Reg. of cell prolif. |
| [DOWN] | PCDH20 | -1,947 | 1,664 | 0,022 | 0,532 | |
| [DOWN] | PCP4 | -2,656 | 2,247 | 0,006 | 0,290 | |
| [DOWN] | PDE10A | -2,410 | 2,910 | 0,001 | 0,101 | |
| [DOWN] | PDE3A | -1,179 | 1,428 | 0,037 | 0,653 | Dev. Proc. |
| [DOWN] | PDGFRB | -5,233 | 3,395 | 0,000 | 0,038 | Dev. Proc. + Reg. of cell prolif. |
| [DOWN] | PDZD4 | -4,457 | 2,081 | 0,008 | 0,343 | |
| [DOWN] | PHC1 | -3,770 | 2,700 | 0,002 | 0,146 | Dev. Proc. |
| [DOWN] | PHF10 | -4,069 | 2,571 | 0,003 | 0,178 | Dev. Proc. |
| [DOWN] | PHF14 | -4,329 | 2,577 | 0,003 | 0,176 | Dev. Proc. + Reg. of cell prolif. |
| [DOWN] | PHF21B | -1,876 | 1,495 | 0,032 | 0,615 | |
| [UP] | PHLDA3 | 0,947 | 1,436 | 0,037 | 0,645 | |
| [DOWN] | PHLPP1 | -1,502 | 2,178 | 0,007 | 0,317 | |
| [DOWN] | PHYHIPL | -12,057 | 12,189 | 0,000 | 0,000 | |
| [DOWN] | PIAS3 | -1,123 | 1,378 | 0,042 | 0,673 | |
| [DOWN] | PLA2R1 | -2,754 | 2,744 | 0,002 | 0,136 | |
| [UP] | PLAUR | 3,767 | 1,365 | 0,043 | 0,673 | Dev. Proc. |
| [DOWN] | PLEKHG4B | -2,726 | 1,455 | 0,035 | 0,637 | |
| [DOWN] | PLEKHG7 | -0,950 | 1,421 | 0,038 | 0,653 | |
| [DOWN] | PLEKHO2 | -6,801 | 6,617 | 0,000 | 0,000 | |
| [UP] | PNCK | 1,787 | 1,604 | 0,025 | 0,568 | |
| [UP] | PNO1 | 1,069 | 1,709 | 0,020 | 0,519 | |
| [DOWN] | POC1A | -6,765 | 5,213 | 0,000 | 0,001 | Dev. Proc. |
| [DOWN] | PODXL | -2,708 | 1,356 | 0,044 | 0,680 | |
| [DOWN] | POLG | -1,239 | 1,342 | 0,046 | 0,680 | |

| | | | | | | | |
|---|---|---|---|---|---|---|---|
| [DOWN] | POMC | -3,295 | 1,649 | 0,022 | 0,540 | | |
| [UP] | POP7 | 1,763 | 1,718 | 0,019 | 0,514 | | |
| [DOWN] | PPEF1 | -4,256 | 1,918 | 0,012 | 0,416 | | |
| [UP] | PPFIA3 | 1,635 | 1,630 | 0,023 | 0,546 | | |
| [UP] | PPIB | 1,052 | 1,668 | 0,021 | 0,532 | Dev. Proc. | |
| [DOWN] | PPIL6 | -4,632 | 2,232 | 0,006 | 0,296 | | |
| [UP] | PPP1R15B | 1,150 | 1,660 | 0,022 | 0,532 | | |
| [DOWN] | PPP1R16B | -4,491 | 2,076 | 0,008 | 0,344 | Dev. Proc. + Reg. of cell prolif. | |
| [UP] | PREB | 2,915 | 2,686 | 0,002 | 0,149 | | |
| [DOWN] | PRG4 | -8,687 | 8,056 | 0,000 | 0,000 | Dev. Proc. + Reg. of cell prolif. | |
| [DOWN] | PRKAG3 | -1,168 | 1,308 | 0,049 | 0,683 | | |
| [DOWN] | PRKCG | -6,058 | 3,329 | 0,000 | 0,043 | Dev. Proc. | |
| [DOWN] | PRRT1 | -4,020 | 1,739 | 0,018 | 0,502 | | |
| [DOWN] | PRSS57 | -4,304 | 1,949 | 0,011 | 0,416 | | |
| [UP] | PSAT1 | 1,138 | 1,667 | 0,022 | 0,532 | | |
| [UP] | PSMA2 | 1,018 | 1,615 | 0,024 | 0,560 | | |
| [DOWN] | PTER | -1,748 | 1,677 | 0,021 | 0,532 | | |
| [DOWN] | PTHLH | -1,093 | 1,489 | 0,032 | 0,618 | Dev. Proc. + Reg. of cell prolif. | X |
| [DOWN] | PTPN21 | -1,118 | 1,517 | 0,030 | 0,598 | | |
| [DOWN] | PTPRC | -3,747 | 4,200 | 0,000 | 0,008 | Dev. Proc. + Reg. of cell prolif. | |
| [UP] | PTPRJ | 1,323 | 1,324 | 0,047 | 0,680 | Dev. Proc. + Reg. of cell prolif. | |
| [DOWN] | PTPRN2 | -1,463 | 1,322 | 0,048 | 0,680 | | |
| [UP] | PTRH2 | 4,768 | 2,049 | 0,009 | 0,358 | | |
| [DOWN] | QRICH2 | -3,063 | 1,425 | 0,038 | 0,653 | | |
| [DOWN] | RAB26 | -4,711 | 2,266 | 0,005 | 0,283 | Dev. Proc. | |
| [DOWN] | RAB38 | -3,583 | 1,406 | 0,039 | 0,653 | Dev. Proc. | |
| [UP] | RAB9B | 3,942 | 1,454 | 0,035 | 0,637 | | |
| [DOWN] | RABGAP1L | -2,194 | 1,458 | 0,035 | 0,637 | Dev. Proc. | |
| [UP] | RACGAP1 | 1,323 | 1,988 | 0,010 | 0,398 | Dev. Proc. | |
| [UP] | RAD51C | 1,149 | 1,726 | 0,019 | 0,510 | Dev. Proc. | |
| [DOWN] | RAD9A | -3,872 | 6,473 | 0,000 | 0,000 | | |
| [UP] | RAG1 | 3,837 | 1,366 | 0,043 | 0,673 | Dev. Proc. | |
| [UP] | RANGAP1 | 1,088 | 1,776 | 0,017 | 0,485 | | |
| [DOWN] | RAPGEF4 | -6,839 | 4,883 | 0,000 | 0,002 | | |
| [UP] | RARRES2 | 2,296 | 1,347 | 0,045 | 0,680 | Dev. Proc. | |
| [DOWN] | RASA4B | -2,018 | 1,849 | 0,014 | 0,448 | | |
| [UP] | RASAL1 | 1,999 | 1,398 | 0,040 | 0,661 | Dev. Proc. | |
| [DOWN] | RASGRF2 | -4,352 | 1,980 | 0,010 | 0,399 | | |
| [DOWN] | RB1CC1 | -2,661 | 1,443 | 0,036 | 0,641 | Dev. Proc. + Reg. of cell prolif. | |
| [UP] | REEP2 | 1,289 | 1,398 | 0,040 | 0,661 | | |
| [DOWN] | RERG | -6,567 | 1,614 | 0,024 | 0,561 | Reg. of cell prolif. | |
| [UP] | REXO4 | 1,112 | 1,680 | 0,021 | 0,532 | | |
| [UP] | RGL3 | 4,332 | 1,678 | 0,021 | 0,532 | | |

| | | | | | | |
|---|---|---|---|---|---|---|
| [DOWN] | RGS17 | -4,544 | 2,142 | 0,007 | 0,329 | |
| [UP] | RGS22 | 3,794 | 1,313 | 0,049 | 0,680 | |
| [UP] | RGS6 | 4,887 | 1,934 | 0,012 | 0,416 | |
| [DOWN] | RGS7 | -3,099 | 1,459 | 0,035 | 0,637 | |
| [DOWN] | RHBDD1 | -5,063 | 3,980 | 0,000 | 0,013 | Dev. Proc. |
| [DOWN] | RHPN1 | -5,238 | 3,554 | 0,000 | 0,029 | Dev. Proc. |
| [DOWN] | RIMS4 | -3,625 | 1,913 | 0,012 | 0,417 | |
| [UP] | RIPK3 | 2,245 | 1,632 | 0,023 | 0,546 | Dev. Proc. + Reg. of cell prolif. |
| [UP] | RMI1 | 2,442 | 1,752 | 0,018 | 0,499 | Dev. Proc. |
| [DOWN] | RNF144B | -3,758 | 3,492 | 0,000 | 0,032 | |
| [UP] | RNF212 | 4,880 | 2,098 | 0,008 | 0,343 | |
| [DOWN] | RNF222 | -3,158 | 3,003 | 0,001 | 0,083 | |
| [DOWN] | ROBO2 | -1,115 | 1,606 | 0,025 | 0,568 | Dev. Proc. |
| [UP] | ROBO4 | 4,111 | 1,545 | 0,029 | 0,589 | Dev. Proc. |
| [UP] | RPF1 | 0,953 | 1,431 | 0,037 | 0,650 | |
| [UP] | RPGRIP1 | 4,710 | 1,938 | 0,012 | 0,416 | Dev. Proc. |
| [DOWN] | RPL3L | -1,607 | 1,460 | 0,035 | 0,637 | |
| [DOWN] | RPRM | -5,156 | 2,700 | 0,002 | 0,146 | |
| [DOWN] | RRBP1 | -1,819 | 3,526 | 0,000 | 0,030 | |
| [DOWN] | RSAD2 | -4,457 | 2,081 | 0,008 | 0,343 | Dev. Proc. |
| [DOWN] | RSPO1 | -3,094 | 1,956 | 0,011 | 0,416 | Reg. of cell prolif. |
| [DOWN] | RSPO3 | -3,441 | 1,341 | 0,046 | 0,680 | Dev. Proc. |
| [DOWN] | RTN4IP1 | -1,464 | 1,374 | 0,042 | 0,673 | Dev. Proc. |
| [DOWN] | SACS | -2,594 | 4,146 | 0,000 | 0,009 | |
| [DOWN] | SAMD14 | -3,370 | 1,310 | 0,049 | 0,680 | Dev. Proc. |
| [DOWN] | SAP130 | -3,139 | 2,206 | 0,006 | 0,305 | |
| [UP] | SARM1 | 3,837 | 1,366 | 0,043 | 0,673 | Dev. Proc. |
| [UP] | SART3 | 0,903 | 1,312 | 0,049 | 0,680 | Dev. Proc. |
| [DOWN] | SCFD1 | -1,995 | 1,729 | 0,019 | 0,508 | |
| [DOWN] | SCFD2 | -1,184 | 1,498 | 0,032 | 0,613 | |
| [UP] | SCGN | 5,021 | 2,272 | 0,005 | 0,283 | |
| [DOWN] | SCN1A | -3,360 | 2,389 | 0,004 | 0,238 | Dev. Proc. |
| [DOWN] | SCNN1A | -3,370 | 1,310 | 0,049 | 0,680 | |
| [UP] | SEC23B | 1,221 | 1,964 | 0,011 | 0,412 | |
| [DOWN] | SEC62 | -4,948 | 2,926 | 0,001 | 0,098 | |
| [DOWN] | SEPT1 | -4,352 | 1,980 | 0,010 | 0,399 | |
| [DOWN] | SEZ6L2 | -3,512 | 1,373 | 0,042 | 0,673 | Dev. Proc. |
| [DOWN] | SFMBT2 | -1,535 | 1,377 | 0,042 | 0,673 | |
| [DOWN] | SFRP4 | -1,406 | 1,540 | 0,029 | 0,589 | Dev. Proc. + Reg. of cell prolif. |
| [UP] | SGPP1 | 1,088 | 1,721 | 0,019 | 0,511 | |
| [UP] | SH3BP2 | 1,422 | 1,534 | 0,029 | 0,589 | |
| [UP] | SH3BP5 | 1,317 | 1,781 | 0,017 | 0,483 | |
| [UP] | SHC1 | 1,197 | 1,482 | 0,033 | 0,624 | Dev. Proc. + Reg. of cell prolif. |

| | | | | | | | |
|---|---|---|---|---|---|---|---|
| [DOWN] | SHE | -3,512 | 1,373 | 0,042 | 0,673 | | |
| [UP] | SHH | 6,221 | 3,442 | 0,000 | 0,035 | Dev. Proc. + Reg. of cell prolif. | X |
| [DOWN] | SIK1 | -1,389 | 2,659 | 0,002 | 0,156 | Dev. Proc. | |
| [UP] | SKA1 | 1,909 | 1,326 | 0,047 | 0,680 | | |
| [DOWN] | SLC10A7 | -4,392 | 3,169 | 0,001 | 0,061 | Dev. Proc. | |
| [DOWN] | SLC15A1 | -13,664 | 9,383 | 0,000 | 0,000 | | |
| [DOWN] | SLC17A6 | -12,147 | 8,606 | 0,000 | 0,000 | Dev. Proc. | |
| [DOWN] | SLC18A2 | -3,689 | 2,122 | 0,008 | 0,336 | Dev. Proc. | |
| [DOWN] | SLC19A1 | -1,069 | 1,352 | 0,044 | 0,680 | | |
| [DOWN] | SLC19A2 | -7,773 | 5,356 | 0,000 | 0,001 | Dev. Proc. | |
| [UP] | SLC1A3 | 5,529 | 2,705 | 0,002 | 0,146 | Dev. Proc. | |
| [DOWN] | SLC22A3 | -1,073 | 1,611 | 0,024 | 0,563 | | |
| [UP] | SLC25A18 | 1,020 | 1,581 | 0,026 | 0,581 | | |
| [DOWN] | SLC25A19 | -4,854 | 4,499 | 0,000 | 0,004 | | |
| [UP] | SLC25A26 | 1,840 | 1,371 | 0,043 | 0,673 | | |
| [UP] | SLC25A30 | 3,156 | 2,194 | 0,006 | 0,311 | | |
| [UP] | SLC25A40 | 3,767 | 1,365 | 0,043 | 0,673 | Dev. Proc. | |
| [UP] | SLC26A1 | 7,047 | 3,217 | 0,001 | 0,055 | | |
| [UP] | SLC27A4 | 1,361 | 1,792 | 0,016 | 0,479 | Dev. Proc. | |
| [UP] | SLC38A1 | 1,015 | 1,319 | 0,048 | 0,680 | | |
| [UP] | SLC38A3 | 3,661 | 2,106 | 0,008 | 0,340 | | |
| [DOWN] | SLC39A4 | -1,166 | 1,302 | 0,050 | 0,686 | | |
| [DOWN] | SLC3A1 | -1,988 | 1,553 | 0,028 | 0,589 | | |
| [UP] | SLC4A1 | 2,227 | 2,034 | 0,009 | 0,368 | Dev. Proc. + Reg. of cell prolif. | |
| [DOWN] | SLC4A3 | -1,334 | 1,392 | 0,041 | 0,668 | | |
| [DOWN] | SLC5A11 | -4,283 | 1,901 | 0,013 | 0,420 | | |
| [DOWN] | SLC6A2 | -4,256 | 1,918 | 0,012 | 0,416 | | |
| [UP] | SLC7A7 | 3,354 | 1,783 | 0,017 | 0,482 | | |
| [UP] | SLC7A9 | 3,755 | 1,317 | 0,048 | 0,680 | | |
| [DOWN] | SLC9A3 | -2,802 | 2,105 | 0,008 | 0,340 | | |
| [DOWN] | SLCO1C1 | -13,842 | 9,314 | 0,000 | 0,000 | | |
| [UP] | SLITRK5 | 2,104 | 1,394 | 0,040 | 0,665 | Dev. Proc. | |
| [UP] | SMARCD1 | 0,952 | 1,451 | 0,035 | 0,637 | Dev. Proc. + Reg. of cell prolif. | |
| [DOWN] | SMARCD3 | -2,542 | 1,802 | 0,016 | 0,474 | Dev. Proc. + Reg. of cell prolif. | |
| [UP] | SMIM22 | 1,257 | 1,311 | 0,049 | 0,680 | Reg. of cell prolif. | |
| [DOWN] | SMTNL2 | -3,583 | 1,406 | 0,039 | 0,653 | | |
| [DOWN] | SNCB | -7,590 | 7,467 | 0,000 | 0,000 | | |
| [DOWN] | SNED1 | -2,783 | 3,378 | 0,000 | 0,040 | | |
| [UP] | SNRPA | 1,555 | 1,473 | 0,034 | 0,632 | | |
| [UP] | SNRPC | 1,301 | 2,228 | 0,006 | 0,297 | | |
| [UP] | SNX1 | 0,909 | 1,334 | 0,046 | 0,680 | Dev. Proc. | |
| [DOWN] | SNX4 | -3,726 | 2,198 | 0,006 | 0,309 | | |
| [DOWN] | SOD3 | -3,017 | 2,178 | 0,007 | 0,317 | | |

| | | | | | | |
|---|---|---|---|---|---|---|
| [DOWN] | SORL1 | -2,947 | 7,378 | 0,000 | 0,000 | Dev. Proc. |
| [DOWN] | SP2 | -4,213 | 3,440 | 0,000 | 0,035 | Dev. Proc. |
| [UP] | SPA17 | 4,977 | 2,207 | 0,006 | 0,305 | |
| [DOWN] | SPAG17 | -2,991 | 1,384 | 0,041 | 0,673 | Dev. Proc. |
| [DOWN] | SPATA13 | -6,069 | 9,609 | 0,000 | 0,000 | |
| [DOWN] | SPATA7 | -5,604 | 3,310 | 0,000 | 0,045 | |
| [DOWN] | SPG21 | -3,549 | 2,340 | 0,005 | 0,258 | Dev. Proc. |
| [DOWN] | SPI1 | -4,610 | 1,692 | 0,020 | 0,531 | Dev. Proc. |
| [UP] | SPINT2 | 1,092 | 1,813 | 0,015 | 0,469 | Dev. Proc. + Reg. of cell prolif. |
| [DOWN] | SPINT4 | -1,536 | 1,867 | 0,014 | 0,435 | |
| [UP] | SRFBP1 | 0,952 | 1,389 | 0,041 | 0,669 | |
| [DOWN] | SRMS | -3,370 | 1,310 | 0,049 | 0,680 | Dev. Proc. |
| [DOWN] | ST18 | -4,537 | 2,108 | 0,008 | 0,340 | Reg. of cell prolif. |
| [DOWN] | ST6GALNAC5 | -2,730 | 2,138 | 0,007 | 0,332 | |
| [DOWN] | STAB2 | -1,484 | 2,156 | 0,007 | 0,323 | |
| [DOWN] | STAP2 | -3,733 | 1,534 | 0,029 | 0,589 | |
| [UP] | STARD10 | 1,826 | 1,723 | 0,019 | 0,511 | |
| [DOWN] | STAT1 | -0,980 | 1,546 | 0,028 | 0,589 | Dev. Proc. + Reg. of cell prolif. |
| [DOWN] | STAT4 | -4,711 | 2,266 | 0,005 | 0,283 | Dev. Proc. + Reg. of cell prolif. |
| [DOWN] | STKLD1 | -4,619 | 3,725 | 0,000 | 0,021 | |
| [DOWN] | STOX1 | -0,900 | 1,387 | 0,041 | 0,672 | Dev. Proc. |
| [UP] | STRA6 | 3,904 | 1,448 | 0,036 | 0,637 | Dev. Proc. |
| [DOWN] | STX11 | -3,289 | 1,594 | 0,025 | 0,570 | |
| [DOWN] | STXBP5L | -4,283 | 1,901 | 0,013 | 0,420 | |
| [DOWN] | SULT2A1 | -3,916 | 1,629 | 0,024 | 0,546 | |
| [DOWN] | SULT4A1 | -3,583 | 1,406 | 0,039 | 0,653 | Dev. Proc. |
| [UP] | SUV39H1 | 1,652 | 1,519 | 0,030 | 0,597 | Dev. Proc. |
| [DOWN] | SV2B | -10,135 | 1,368 | 0,043 | 0,673 | |
| [UP] | SWI5 | 1,468 | 1,474 | 0,034 | 0,631 | |
| [DOWN] | SYCP1 | -11,970 | 6,412 | 0,000 | 0,000 | Dev. Proc. |
| [UP] | SYNPO2 | 1,112 | 1,456 | 0,035 | 0,637 | |
| [DOWN] | SYT6 | -4,180 | 1,835 | 0,015 | 0,457 | |
| [DOWN] | SYT7 | -7,654 | 7,249 | 0,000 | 0,000 | |
| [DOWN] | SZT2 | -2,666 | 1,797 | 0,016 | 0,476 | Dev. Proc. |
| [DOWN] | TACR2 | -3,370 | 1,310 | 0,049 | 0,680 | |
| [DOWN] | TARSL2 | -7,444 | 4,866 | 0,000 | 0,002 | |
| [DOWN] | TBX15 | -4,283 | 1,901 | 0,013 | 0,420 | Dev. Proc. |
| [DOWN] | TC2N | -1,261 | 1,367 | 0,043 | 0,673 | |
| [DOWN] | TCP11 | -12,774 | 9,774 | 0,000 | 0,000 | Dev. Proc. |
| [DOWN] | TDRKH | -3,794 | 1,565 | 0,027 | 0,587 | Dev. Proc. |
| [UP] | TECTA | 4,560 | 1,856 | 0,014 | 0,442 | Dev. Proc. |
| [DOWN] | TEKT2 | -4,751 | 2,297 | 0,005 | 0,275 | |
| [DOWN] | TEKT4 | -3,916 | 1,629 | 0,024 | 0,546 | |

| | | | | | | |
|---|---|---|---|---|---|---|
| [DOWN] | TENM2 | -2,555 | 1,649 | 0,022 | 0,540 | Dev. Proc. |
| [UP] | TESC | 3,389 | 2,353 | 0,004 | 0,253 | Dev. Proc. + Reg. of cell prolif. |
| [UP] | TGDS | 1,356 | 1,501 | 0,032 | 0,610 | |
| [DOWN] | TGFB3 | -1,077 | 1,325 | 0,047 | 0,680 | Dev. Proc. + Reg. of cell prolif. |
| [DOWN] | TGM4 | -1,636 | 2,170 | 0,007 | 0,318 | |
| [DOWN] | THEMIS2 | -2,887 | 1,307 | 0,049 | 0,684 | |
| [UP] | THNSL2 | 1,240 | 1,312 | 0,049 | 0,680 | |
| [UP] | THOC7 | 1,217 | 2,065 | 0,009 | 0,348 | |
| [UP] | THSD7B | 3,924 | 1,367 | 0,043 | 0,673 | |
| [DOWN] | THY1 | -6,964 | 9,634 | 0,000 | 0,000 | Dev. Proc. |
| [DOWN] | TIMM10B | -3,512 | 1,373 | 0,042 | 0,673 | |
| [UP] | TIMM17B | 1,234 | 1,685 | 0,021 | 0,532 | |
| [DOWN] | TLL1 | -9,859 | 8,204 | 0,000 | 0,000 | Dev. Proc. |
| [DOWN] | TM4SF19 | -4,937 | 2,471 | 0,003 | 0,209 | |
| [DOWN] | TMC2 | -3,370 | 1,310 | 0,049 | 0,680 | |
| [DOWN] | TMCC1 | -3,142 | 4,882 | 0,000 | 0,002 | |
| [DOWN] | TMEM117 | -1,809 | 1,483 | 0,033 | 0,623 | |
| [DOWN] | TMEM131 | -3,480 | 1,939 | 0,012 | 0,416 | |
| [DOWN] | TMEM145 | -10,662 | 1,311 | 0,049 | 0,680 | |
| [DOWN] | TMEM151B | -0,997 | 1,314 | 0,049 | 0,680 | |
| [UP] | TMEM168 | 1,121 | 1,793 | 0,016 | 0,479 | |
| [DOWN] | TMEM211 | -2,360 | 2,063 | 0,009 | 0,348 | |
| [UP] | TMEM237 | 1,430 | 1,354 | 0,044 | 0,680 | |
| [DOWN] | TMEM26 | -4,531 | 3,352 | 0,000 | 0,041 | |
| [DOWN] | TMEM40 | -4,302 | 1,634 | 0,023 | 0,546 | |
| [UP] | TMEM72 | 4,507 | 1,629 | 0,023 | 0,546 | |
| [DOWN] | TMEM88 | -4,399 | 2,012 | 0,010 | 0,380 | Dev. Proc. |
| [UP] | TMPRSS4 | 4,285 | 1,683 | 0,021 | 0,532 | |
| [DOWN] | TMPRSS7 | -4,074 | 1,770 | 0,017 | 0,485 | |
| [UP] | TMPRSS9 | 4,285 | 1,683 | 0,021 | 0,532 | |
| [DOWN] | TNC | -1,142 | 1,849 | 0,014 | 0,448 | Dev. Proc. + Reg. of cell prolif. |
| [DOWN] | TNFAIP8 | -0,887 | 1,323 | 0,048 | 0,680 | |
| [UP] | TNFRSF18 | 4,242 | 1,668 | 0,021 | 0,532 | |
| [UP] | TNFRSF6B | 1,483 | 1,572 | 0,027 | 0,587 | |
| [DOWN] | TOLLIP | -2,828 | 2,686 | 0,002 | 0,149 | |
| [DOWN] | TOX4 | -5,039 | 6,322 | 0,000 | 0,000 | |
| [DOWN] | TPD52L1 | -1,727 | 1,583 | 0,026 | 0,579 | |
| [UP] | TPH1 | 4,358 | 1,554 | 0,028 | 0,589 | Dev. Proc. |
| [UP] | TPMT | 1,499 | 1,709 | 0,020 | 0,519 | |
| [DOWN] | TPPP3 | -2,811 | 1,597 | 0,025 | 0,568 | Dev. Proc. |
| [DOWN] | TRH | -3,512 | 1,373 | 0,042 | 0,673 | |
| [DOWN] | TRIM25 | -1,432 | 2,329 | 0,005 | 0,261 | |
| [DOWN] | TRIM56 | -3,370 | 1,310 | 0,049 | 0,680 | |

| | | | | | | |
|---|---|---|---|---|---|---|
| [UP] | TRMT10C | 1,171 | 1,947 | 0,011 | 0,416 | |
| [UP] | TRMT2A | 0,969 | 1,438 | 0,036 | 0,645 | |
| [DOWN] | TRNAC-GCA | -3,512 | 1,373 | 0,042 | 0,673 | |
| [UP] | TRNAK-CUU | 3,094 | 1,435 | 0,037 | 0,646 | |
| [UP] | TRNAL-AAG | 5,378 | 2,565 | 0,003 | 0,179 | |
| [UP] | TRNAL-CAA | 3,836 | 2,524 | 0,003 | 0,191 | |
| [UP] | TRNAP-CGG | 4,425 | 1,769 | 0,017 | 0,485 | |
| [UP] | TRNAS-GCU | 3,110 | 1,532 | 0,029 | 0,590 | |
| [UP] | TRNAV-CAC | 3,305 | 1,675 | 0,021 | 0,532 | |
| [DOWN] | TRNAV-UAC | -2,874 | 1,452 | 0,035 | 0,637 | |
| [DOWN] | TRPA1 | -9,941 | 9,199 | 0,000 | 0,000 | |
| [DOWN] | TRPC4 | -4,304 | 1,949 | 0,011 | 0,416 | Dev. Proc. |
| [DOWN] | TRPV3 | -2,271 | 1,480 | 0,033 | 0,624 | |
| [UP] | TRPV4 | 4,506 | 1,711 | 0,019 | 0,517 | Dev. Proc. |
| [DOWN] | TSC1 | -1,006 | 1,658 | 0,022 | 0,534 | Dev. Proc. + Reg. of cell prolif. |
| [DOWN] | TSHR | -5,198 | 2,771 | 0,002 | 0,130 | Dev. Proc. + Reg. of cell prolif. |
| [DOWN] | TSPAN1 | -1,562 | 1,335 | 0,046 | 0,680 | |
| [UP] | TSPEAR | 6,221 | 3,442 | 0,000 | 0,035 | Dev. Proc. |
| [DOWN] | TSTD1 | -4,457 | 2,081 | 0,008 | 0,343 | |
| [DOWN] | TTC13 | -5,861 | 4,067 | 0,000 | 0,011 | |
| [DOWN] | TTC16 | -3,916 | 1,629 | 0,024 | 0,546 | |
| [DOWN] | TTC29 | -5,141 | 1,747 | 0,018 | 0,501 | |
| [DOWN] | TTI2 | -7,114 | 4,962 | 0,000 | 0,002 | |
| [UP] | TUBB4B | 0,953 | 1,441 | 0,036 | 0,642 | |
| [DOWN] | TULP1 | -4,256 | 1,918 | 0,012 | 0,416 | Dev. Proc. |
| [DOWN] | TULP2 | -5,856 | 3,745 | 0,000 | 0,021 | |
| [DOWN] | TYMP | -4,932 | 2,488 | 0,003 | 0,203 | Dev. Proc. |
| [UP] | TYROBP | 2,028 | 1,456 | 0,035 | 0,637 | Dev. Proc. + Reg. of cell prolif. |
| [DOWN] | TYRP1 | -4,256 | 1,918 | 0,012 | 0,416 | Dev. Proc. |
| [DOWN] | UBE2B | -5,505 | 3,658 | 0,000 | 0,024 | Dev. Proc. |
| [UP] | UBE2Q1 | 0,905 | 1,315 | 0,048 | 0,680 | Dev. Proc. |
| [DOWN] | UBXN11 | -4,256 | 1,918 | 0,012 | 0,416 | |
| [UP] | UCHL3 | 1,228 | 1,916 | 0,012 | 0,416 | Dev. Proc. |
| [UP] | UNC79 | 4,027 | 1,526 | 0,030 | 0,595 | Dev. Proc. |
| [DOWN] | USH2A | -4,996 | 2,093 | 0,008 | 0,343 | Dev. Proc. |
| [DOWN] | USP18 | -5,050 | 4,927 | 0,000 | 0,002 | |
| [DOWN] | USP33 | -4,105 | 2,556 | 0,003 | 0,181 | Dev. Proc. |
| [DOWN] | USP43 | -4,232 | 1,868 | 0,014 | 0,435 | |
| [UP] | VAV2 | 1,317 | 1,667 | 0,022 | 0,532 | Dev. Proc. |
| [UP] | VWA2 | 2,170 | 2,392 | 0,004 | 0,238 | |
| [UP] | VWA7 | 4,973 | 2,158 | 0,007 | 0,323 | |
| [DOWN] | VWF | -1,514 | 3,139 | 0,001 | 0,064 | Dev. Proc. |
| [DOWN] | WASHC4 | -2,589 | 1,821 | 0,015 | 0,464 | |

| | | | | | | |
|---|---|---|---|---|---|---|
| [UP] | WBP2 | 1,136 | 1,876 | 0,013 | 0,432 | |
| [DOWN] | WDR49 | -5,784 | 3,640 | 0,000 | 0,025 | |
| [DOWN] | WDR97 | -4,180 | 1,835 | 0,015 | 0,457 | |
| [UP] | WNT7A | 3,904 | 1,448 | 0,036 | 0,637 | Dev. Proc. + Reg. of cell prolif. |
| [UP] | WNT9A | 3,460 | 1,998 | 0,010 | 0,389 | Dev. Proc. + Reg. of cell prolif. |
| [DOWN] | XBP1 | -1,953 | 2,231 | 0,006 | 0,296 | Dev. Proc. + Reg. of cell prolif. |
| [DOWN] | XIAP | -2,033 | 1,390 | 0,041 | 0,668 | |
| [DOWN] | XKR8 | -1,125 | 1,329 | 0,047 | 0,680 | Dev. Proc. |
| [UP] | XRCC2 | 2,134 | 1,342 | 0,046 | 0,680 | Dev. Proc. |
| [UP] | XRCC3 | 4,181 | 3,044 | 0,001 | 0,077 | |
| [UP] | YBX2 | 4,285 | 1,683 | 0,021 | 0,532 | Dev. Proc. |
| [DOWN] | YEATS2 | -4,408 | 2,730 | 0,002 | 0,138 | |
| [DOWN] | ZBTB24 | -3,090 | 3,685 | 0,000 | 0,023 | Dev. Proc. |
| [UP] | ZBTB8B | 2,119 | 1,338 | 0,046 | 0,680 | |
| [UP] | ZC3H18 | 1,456 | 2,083 | 0,008 | 0,343 | |
| [DOWN] | ZDHHC23 | -7,664 | 5,329 | 0,000 | 0,001 | |
| [UP] | ZDHHC6 | 0,995 | 1,507 | 0,031 | 0,605 | |
| [DOWN] | ZFP36 | -1,061 | 1,445 | 0,036 | 0,639 | Dev. Proc. + Reg. of cell prolif. |
| [UP] | ZNF276 | 1,431 | 1,316 | 0,048 | 0,680 | |
| [DOWN] | ZNF385C | -1,121 | 1,420 | 0,038 | 0,653 | |
| [UP] | ZNF410 | 0,980 | 1,351 | 0,045 | 0,680 | |
| [UP] | ZNF503 | 2,996 | 1,418 | 0,038 | 0,653 | Dev. Proc. + Reg. of cell prolif. |
| [UP] | ZNHIT2 | 4,027 | 1,466 | 0,034 | 0,635 | |

**Table S3. Tooth width of teeth in *A. carolinensis* hatchlings (in μm).**

| | | Right lower jaw | Left lower jaw |
|---|---|---|---|
| Sp, | Tooth | Width | Width |
| AC120 | anterior_T2 | 104,76 | 101,01 |
| | anterior_T3 | 110,12 | 121,25 |
| | anterior_T4 | 107,11 | - |
| | anterior_T5 | 121,68 | 140,50 |
| | anterior_T6 | 125,12 | 134,21 |
| | anterior_T7 | 57,08 | 106,18 |
| | middle_T1 | 131,43 | 147,55 |
| | middle_T2 | 154,29 | 161,42 |
| | middle_T3 | 149,86 | 163,98 |
| | middle_T4 | 189,71 | 195,79 |
| | middle_T5 | 156,79 | 156,18 |
| | posterior_T1 | 211,74 | 201,54 |
| | posterior_T2 | 204,74 | 213,71 |
| | posterior_T3 | 237,39 | 247,62 |
| AC171 | anterior_T1 | 53,84 | 79,81 |
| | anterior_T2 | 113,57 | 114,75 |
| | anterior_T3 | 118,47 | 118,72 |
| | anterior_T5 | 121,37 | 132,37 |
| | anterior_T7 | 107,36 | 111,34 |
| | middle_T1 | 149,26 | 144,57 |
| | middle_T3 | 155,04 | 165,58 |
| | middle_T4 | 197,47 | 209,22 |
| | middle_T5 | 169,29 | 174,71 |
| | posterior_T1 | 222,23 | - |
| | posterior_T2 | 230,44 | 230,80 |
| | posterior_T3 | 255,02 | 267,99 |
| AC64 | anterior_T1 | 81,80 | - |
| | anterior_T2 | 93,24 | 92,17 |
| | anterior_T3 | 80,33 | 106,96 |
| | anterior_T4 | 104,28 | 135,13 |
| | anterior_T5 | 78,58 | 121,59 |
| | anterior_T6 | 110,83 | 135,85 |
| | anterior_T7 | 57,56 | 74,25 |
| | middle_T1 | 133,07 | 132,97 |
| | middle_T2 | 129,02 | 152,01 |
| | middle_T3 | 141,05 | 148,24 |
| | middle_T4 | 177,48 | 189,32 |

| | | | |
|---|---|---|---|
| | middle_T5 | 158,83 | 166,92 |
| | posterior_T1 | 195,36 | 187,58 |
| | posterior_T2 | 197,07 | 194,93 |
| | posterior_T3 | 219,13 | 228,22 |
| AC93 | anterior_T1 | 68,45 | 48,65 |
| | anterior_T2 | 99,58 | 81,78 |
| | anterior_T3 | 113,27 | 98,40 |
| | anterior_T5 | 110,97 | 118,42 |
| | anterior_T6 | 119,85 | 130,34 |
| | anterior_T7 | 73,16 | 31,46 |
| | middle_T1 | 140,90 | 142,72 |
| | middle_T3 | 161,17 | 159,62 |
| | middle_T4 | 177,32 | 184,76 |
| | middle_T5 | 142,11 | 150,95 |
| | posterior_T2 | 195,05 | 204,02 |
| | posterior_T3 | 206,09 | 215,73 |
| AC105 | anterior_T2 | 111,15 | 100,54 |
| | anterior_T3 | 120,75 | 117,79 |
| | anterior_T5 | 137,61 | 133,54 |
| | anterior_T7 | 103,21 | 77,48 |
| | middle_T1 | 156,22 | 151,96 |
| | middle_T3 | 184,48 | 175,96 |
| | middle_T4 | 211,23 | 220,36 |
| | middle_T5 | 166,19 | 182,77 |
| | posterior_T2 | 239,72 | 233,02 |
| | posterior_T3 | 268,18 | 270,05 |
| AC201 | anterior_T1 | 49,39 | 54,99 |
| | anterior_T2 | 74,55 | 88,23 |
| | anterior_T3 | 93,73 | 105,53 |
| | anterior_T5 | 103,23 | 106,78 |
| | anterior_T6 | 113,64 | 120,34 |
| | anterior_T7 | 51,43 | 82,14 |
| | middle_T1 | 131,41 | 125,50 |
| | middle_T3 | 147,60 | 157,54 |
| | middle_T4 | 181,30 | 183,49 |
| | middle_T5 | 143,37 | 153,11 |
| | posterior_T2 | 182,26 | 188,32 |
| | posterior_T3 | 227,36 | 228,72 |

**Table S4.** Volumes of tooth buds and *SHH* expression areas of embryonic uni- and tricuspid teeth in *A. carolinensis* at early and late cap stages (in µm³), and the calculated *SHH*/tooth bud ratio. dpo, days postoviposition; UJ, upper jaw; LJ, lower jaw.

| Sample ID | Jaw | Tooth N | dpo | uni/tri | Stage | tooth bud volume | *SHH* volume | *SHH*/tooth bud |
|---|---|---|---|---|---|---|---|---|
| AC253R | UJ | 2 | 15 | uni | Early cap | 69,06 | 8,61 | 0,12 |
| GDAc60.1 | LJ | 5 | 16 | uni | Early cap | 100,86 | 10,15 | 0,10 |
| AC253R | UJ | 9 | 15 | uni | Early cap | 126,60 | 25,70 | 0,20 |
| GDAc58.2 | LJ | 2 | 15 | uni | Early cap | 91,03 | 16,52 | 0,18 |
| AC253R | LJ | 2 | 15 | uni | Early cap | 83,27 | 25,61 | 0,31 |
| GDAc58.2 | UJ | 2 | 15 | uni | Early cap | 149,18 | 37,97 | 0,25 |
| GDAc58.2 | LJ | 4 | 15 | uni | Early cap | 96,23 | 19,59 | 0,20 |
| AC253R | LJ | 4 | 15 | uni | Late cap | 160,87 | 41,75 | 0,26 |
| GDAc60.1 | UJ | 2 | 16 | uni | Late cap | 162,12 | 32,51 | 0,20 |
| AC263L | LJ | 4 | 16 | uni | Late cap | 204,01 | 56,96 | 0,28 |
| GDAc32.2 | UJ | 2 | 19 | uni | Late cap | 222,09 | 60,24 | 0,27 |
| GDAc58.2 | LJ | 6 | 15 | uni | Late cap | 166,34 | 48,50 | 0,29 |
| GDAc60.1 | LJ | 2 | 16 | uni | Late cap | 123,85 | 32,92 | 0,27 |
| GDAc60.2 | LJ | 2 | 16 | uni | Late cap | 100,34 | 26,19 | 0,26 |
| AC263L | LJ | 8 | 16 | uni | Late cap | 207,70 | 66,68 | 0,32 |
| GDAc34.2 | UJ | 20 | 18 | tri | Early cap | 145,27 | 26,48 | 0,18 |
| GDAc58.2 | LJ | 16 | 15 | tri | Early cap | 224,10 | 68,52 | 0,31 |
| GDAc58.2 | UJ | 13 | 15 | tri | Early cap | 183,08 | 42,20 | 0,23 |
| GDAc32.2 | UJ | 13 | 19 | tri | Early cap | 328,79 | 63,74 | 0,19 |
| AC263L | UJ | 15 | 16 | tri | Early cap | 233,18 | 56,20 | 0,24 |
| GDAc34.2 | UJ | 15 | 18 | tri | Early cap | 283,47 | 47,60 | 0,17 |
| GDAc60.1 | LJ | 18 | 16 | tri | Early cap | 147,47 | 24,62 | 0,17 |
| GDAc60.2 | LJ | 18 | 16 | tri | Late cap | 319,60 | 108,73 | 0,34 |
| GDAc60.1 | LJ | 21 | 16 | tri | Late cap | 251,04 | 91,27 | 0,36 |
| GDAc32.2 | LJ | 21 | 19 | tri | Late cap | 329,95 | 113,01 | 0,34 |
| AC263L | LJ | 21 | 16 | tri | Late cap | 455,08 | 139,96 | 0,31 |
| GDAc32.2 | UJ | 18 | 19 | tri | Late cap | 419,40 | 200,04 | 0,48 |
| GDAc60.2 | LJ | 21 | 16 | tri | Late cap | 363,48 | 133,44 | 0,37 |
| GDAc60.1 | UJ | 18 | 16 | tri | Late cap | 321,02 | 130,21 | 0,41 |
| GDAc32.2 | LJ | 16 | 19 | tri | Late cap | 329,80 | 115,99 | 0,35 |

**Table S5.** Perimeters of IEE cells of embryonic uni- and tricuspid teeth in A. *carolinensis* (in µm²).

| Specimen | uni/tri | Tooth N | Stage | Cell perim. | Specimen | uni/tri | Tooth N | Stage | Cell perim. |
|---|---|---|---|---|---|---|---|---|---|
| AC270L | tri | UJ T16 | Late cap | 46,79 | AC270L | uni | UJT7 | Late cap | 40,98 |
| AC270L | tri | UJ T16 | Late cap | 44,15 | AC270L | uni | UJT7 | Late cap | 43,2 |
| AC270L | tri | UJ T16 | Late cap | 43,1 | AC270L | uni | UJT7 | Late cap | 33,69 |
| AC270L | tri | UJ T16 | Late cap | 38,66 | AC270L | uni | UJT7 | Late cap | 24,4 |
| AC270L | tri | UJ T16 | Late cap | 35,81 | AC270L | uni | UJT7 | Late cap | 36,76 |
| AC270L | tri | UJ T16 | Late cap | 35,49 | AC270L | uni | UJT7 | Late cap | 13,84 |
| AC270L | tri | UJ T16 | Late cap | 35,49 | AC270L | uni | UJT7 | Late cap | 23,87 |
| AC270L | tri | UJ T16 | Late cap | 26,09 | AC270L | uni | UJT7 | Late cap | 24,29 |
| AC270L | tri | UJ T16 | Late cap | 24,08 | AC270L | uni | UJT7 | Late cap | 25,98 |
| AC270L | tri | UJ T16 | Late cap | 23,66 | AC270L | uni | UJT7 | Late cap | 22,92 |
| AC270L | tri | UJ T16 | Late cap | 21,76 | AC270L | uni | UJT7 | Late cap | 12,57 |
| AC270L | tri | UJ T16 | Late cap | 21,23 | AC270L | uni | UJT7 | Late cap | 32,64 |
| AC270L | tri | UJ T16 | Late cap | 16,58 | AC270L | uni | UJT7 | Late cap | 32,43 |
| AC270L | tri | UJ T16 | Late cap | 15,95 | AC270L | uni | LJT6 | Late cap | 37,71 |
| AC270L | tri | UJ T16 | Late cap | 41,19 | AC270L | uni | LJT6 | Late cap | 17,46 |
| AC270L | tri | UJ T16 | Late cap | 45,42 | AC270L | uni | LJT6 | Late cap | 39,4 |
| AC270L | tri | UJ T16 | Late cap | 23,03 | AC270L | uni | LJT6 | Late cap | 40,46 |
| AC270L | tri | UJ T16 | Late cap | 21,86 | AC270L | uni | LJT6 | Late cap | 26,72 |
| AC270L | tri | UJ T16 | Late cap | 21,86 | AC270L | uni | LJT6 | Late cap | 30,1 |
| AC270L | tri | UJ T16 | Late cap | 30,74 | AC270L | uni | LJT6 | Late cap | 40,14 |
| AC270L | tri | LJT19 | Early cap | 37,39 | AC270L | uni | UJT5 | Early cap | 15,15 |
| AC270L | tri | LJT19 | Early cap | 45,95 | AC270L | uni | UJT5 | Early cap | 28,84 |
| AC270L | tri | LJT19 | Early cap | 42,99 | AC270L | uni | UJT5 | Early cap | 36,76 |
| AC270L | tri | LJT19 | Early cap | 37,5 | AC270L | uni | UJT5 | Early cap | 20,6 |
| AC270L | tri | LJT19 | Early cap | 25,25 | AC270L | uni | UJT5 | Early cap | 15,32 |
| AC270L | tri | LJT19 | Early cap | 17,11 | AC270L | uni | UJT5 | Early cap | 22,84 |
| AC270L | tri | LJT19 | Early cap | 41,41 | AC270L | uni | UJT5 | Early cap | 30,53 |
| AC270L | tri | LJT19 | Early cap | 34,33 | AC270L | uni | UJT5 | Early cap | 33,17 |
| AC270L | tri | LJT19 | Early cap | 22,82 | AC247L | uni | LJT4 | Early cap | 22,5 |
| AC270L | tri | LJT19 | Early cap | 21,55 | AC247L | uni | LJT4 | Early cap | 33,48 |
| AC270L | tri | LJT19 | Early cap | 15,32 | AC270L | uni | LJT1/2 | Early cap | 37,29 |
| AC270L | tri | LJT19 | Early cap | 21,55 | AC270L | uni | LJT1/2 | Early cap | 24,72 |
| AC270L | tri | LJT19 | Early cap | 8,56 | AC270L | uni | LJT1/2 | Early cap | 32,96 |
| AC270L | tri | LJT19 | Early cap | 8,98 | AC270L | uni | LJT4 | Early cap | 43,52 |
| AC270L | tri | LJT19 | Early cap | 18,38 | AC270L | uni | LJT4 | Early cap | 28,52 |
| AC270L | tri | LJT19 | Early cap | 26,2 | AC270L | uni | LJT4 | Early cap | 33,7 |
| AC270L | tri | LJT19 | Early cap | 29,15 | AC270L | uni | LJT4 | Early cap | 23,34 |
| AC270L | tri | LJT19 | Early cap | 30,32 | AC270L | uni | LJT4 | Early cap | 37,92 |
| AC270L | tri | LJT19 | Early cap | 40,56 | AC270L | uni | LJT11 | Early cap | 31,69 |
| AC270L | tri | LJT19 | Early cap | 32,01 | AC270L | uni | LJT11 | Early cap | 20,21 |

**Table S6.** Model parameter values corresponding to the representative *in silico* tooth models shown in Fig. 4 for each of the six species. GS, *Gerrhosaurus skoogi*; AA, *Ameiva ameiva*; TS, *Takydromus sexlineatus*; CC, *Chamaeleo calyptratus*; PV, *Pogona vitticeps*; AC Tric, *Anolis carolinensis, tricuspid tooth*; AC Unic, *Anolis carolinensis, unicuspid tooth.*

| Parameters | Par | GS | AA | TS | CC | PV | AC Tric | AC Unic |
|---|---|---|---|---|---|---|---|---|
| Epithelial proliferation rate | Egr | 0,027 | 0,045 | 0,028 | 0,042 | 0,031 | 0,028 | 0,025 |
| Mesenchymal proliferation rate | Mgr | 5140 | 6153 | 8163 | 12961 | 19248 | 16545 | 16545 |
| Young's modulus (stiffness) | Rep | 7,94 | 8,43 | 4,46 | 2,69 | 1,13 | 8,01 | 8,01 |
| Distance from 0 where the borders are defined | Swi | 1,08 | 0,740 | 1,441 | 1,939 | 1,11 | 0,264 | 0,264 |
| Traction between neighbours | Adh | 0,0074 | 0,0120 | 0,0153 | 0,0133 | 0,0110 | 0,0165 | 0,0165 |
| Activator auto-activation | Act | 0,87 | 1,060 | 1,341 | 1,133 | 2,60 | 0,378 | 0,378 |
| Inhibition of activator | Inh | 497 | 955 | 920 | 536 | 375 | 799 | 713 |
| Growth factor secretion rate | Sec | 0,056 | 0,220 | 0,143 | 0,218 | 0,055 | 0,136 | 0,136 |
| Activator diffusion rate | Da | 0,40 | 1,90 | 1,32 | 1,39 | 0,12 | 1,69 | 1,69 |
| Inhibitor diffusion rate | Di | 0,83 | 1,23 | 0,73 | 1,38 | 1,42 | 0,86 | 0,86 |
| Growth factor diffusion rate | Ds | 0,450 | 0,690 | 0,662 | 1,518 | 0,057 | 0,760 | 0,760 |
| Initial inhibitor threshold | Int | 0,770 | 0,580 | 0,176 | 0,487 | 0,068 | 0,930 | 0,930 |
| Growth factor threshold | Set | 0,78 | 0,700 | 0,632 | 0,863 | 0,79 | 0,834 | 0,834 |
| Mesenchyme mechanic resistance | Boy | 0,66 | 0,680 | 0,750 | 0,709 | 0,082 | 0,920 | 0,920 |
| Differentiation rate | Dff | 0,00036 | 0,00033 | 0,00002 | 0,00032 | 0,00023 | 0,00014 | 0,00014 |
| Border growth, amount of mesenchyme in anterior-posterior | Bgr | 1,270 | 1,220 | 1,419 | 1,141 | 0,810 | 1,035 | 1,035 |
| Anterior bias | Abi | 6,67 | 20,00 | 16,17 | 2,91 | 3,72 | 2,84 | 2,84 |
| Posterior bias | Pbi | 2,09 | 20,00 | 16,17 | 2,91 | 3,93 | 2,84 | 2,84 |
| Lingual bias | Lbi | 0,72 | 0,32 | 0,79 | 0,04 | 1,39 | 0,26 | 0,26 |
| Buccal bias | Bbi | 0,30 | 0,32 | 0,79 | 0,04 | 0,06 | 0,26 | 0,26 |
| Radius of initial conditions | Rad | 2 | 2 | 2 | 2 | 3 | 2 | 2 |
| Protein degradation rate | Deg | 0,11 | 0,29 | 0,24 | 0,06 | 0,11 | 0,20 | 0,20 |
| Downward vector of growth | Dgr | 28274 | 4415 | 14532 | 15367 | 12727 | 2025 | 2025 |
| Mechanical traction from the borders to the nucleus | Ntr | 0,000074 | 0,000110 | 0,000019 | 0,000149 | 0,000065 | 0,000007 | 0,000007 |
| Width of border | Bwi | 0,50 | 0,58 | 0,38 | 0,55 | 4,73 | 3,36 | 3,36 |
| Initial activator concentration | Ina | 5,66 | 5,50 | 27,95 | 18,15 | 4,16 | 33,26 | 33,26 |
| Basal mesenchymal proliferation rate | uMgr | 77 | 216 | 475 | 401 | 318,00 | 192 | 192 |

**Table S7.** Normalized parameter variability values for all model parameters across the six analyzed species. For each parameter and species, variability was calculated as the range (maximum minus minimum) of normalized parameter values obtained from three independent *in silico* modeling runs in BITES. Sp., species; Pr., parameters; Par. Range, range of parameter values; Raw, raw parameter values from the run; Normalized, normalized values to a [0,1] scale using min-max scaling; Range of norm. val., range of normalized values.

| Sp. | Par. | Par. Range | | Raw | | | Normalized | | | Range of norm. val. |
|---|---|---|---|---|---|---|---|---|---|---|
| | | MIN | MAX | 1 | 2 | 3 | 1 | 2 | 3 | Max-Min |
| *Gerrhosaurus skoogi* | Egr | 0 | 0,05 | 0,042 | 0,038 | 0,045 | 0,80 | 0,70 | 0,88 | 0,18 |
| | Mgr | 0 | 20000 | 14260 | 11491 | 19726 | 0,71 | 0,57 | 0,99 | 0,41 |
| | Rep | 0 | 10 | 0,29 | 4,96 | 3,31 | 0,03 | 0,50 | 0,33 | 0,47 |
| | Swi | 0 | 2 | 0,98 | 1,73 | 1,35 | 0,49 | 0,87 | 0,68 | 0,38 |
| | Adh | 0 | 0,02000 | 0,0021 | 0,0034 | 0,0044 | 0,11 | 0,17 | 0,22 | 0,12 |
| | Act | 0 | 3 | 1,43 | 2,11 | 2,15 | 0,47 | 0,70 | 0,72 | 0,24 |
| | Inh | 1 | 1000 | 328 | 388 | 294 | 0,33 | 0,39 | 0,29 | 0,09 |
| | Sec | 0 | 0,30 | 0,26 | 0,28 | 0,24 | 0,87 | 0,93 | 0,80 | 0,13 |
| | Da | 0 | 2 | 1,89 | 2 | 1,55 | 0,95 | 1,00 | 0,78 | 0,23 |
| | Di | 0 | 2 | 1,8 | 1,13 | 1,95 | 0,90 | 0,57 | 0,98 | 0,41 |
| | Ds | 0 | 2 | 1,75 | 1,12 | 1,59 | 0,88 | 0,56 | 0,80 | 0,32 |
| | Int | 0 | 1 | 0,96 | 0,43 | 0,71 | 0,96 | 0,43 | 0,71 | 0,53 |
| | Set | 0,5 | 1 | 0,6 | 0,53 | 0,5 | 0,20 | 0,06 | 0,00 | 0,20 |
| | Boy | 0 | 1 | 0,29 | 0,27 | 0,34 | 0,29 | 0,27 | 0,34 | 0,07 |
| | Dff | 0 | 0,00100 | 0,000075 | 0,00023 | 0,00033 | 0,08 | 0,23 | 0,33 | 0,26 |
| | Bgr | 0 | 2 | 0,95 | 1,17 | 1,47 | 0,48 | 0,59 | 0,74 | 0,26 |
| | Abi | 0 | 20 | 4,1 | 4,31 | 4,13 | 0,21 | 0,22 | 0,21 | 0,01 |
| | Pbi | 0 | 20 | 4,1 | 4,31 | 4,13 | 0,21 | 0,22 | 0,21 | 0,01 |
| | Lbi | 0 | 3 | 0,57 | 0,073 | 0,84 | 0,19 | 0,02 | 0,28 | 0,26 |
| | Bbi | 0 | 3 | 0,57 | 0,073 | 0,84 | 0,19 | 0,02 | 0,28 | 0,26 |
| | Rad | 2 | 4 | 2 | 2 | 2 | 0,00 | 0,00 | 0,00 | 0,00 |
| | Deg | 0 | 0,3 | 0,019 | 0,29 | 0,14 | 0,06 | 0,97 | 0,47 | 0,90 |
| | Dgr | 0 | 30000 | 3722 | 11947 | 5420 | 0,12 | 0,40 | 0,18 | 0,27 |
| | Ntr | 0 | 0,00020 | 0,000024 | 0,000017 | 0,00016 | 0,12 | 0,09 | 0,80 | 0,72 |
| | Bwi | 0 | 5 | 4,36 | 4,46 | 4,4 | 0,87 | 0,89 | 0,88 | 0,02 |
| | Ina | 0 | 100 | 8,92 | 50 | 6,27 | 0,09 | 0,50 | 0,06 | 0,44 |
| | uMgr | 0 | 500 | 335 | 173 | 217 | 0,67 | 0,35 | 0,43 | 0,32 |
| *Ameiva ameiva* | Egr | 0 | 0,05 | 0,045 | 0,038 | 0,04 | 0,88 | 0,70 | 0,75 | 0,18 |
| | Mgr | 0 | 20000 | 6153 | 7768 | 4433 | 0,31 | 0,39 | 0,22 | 0,17 |
| | Rep | 0 | 10 | 8,43 | 5,77 | 4,62 | 0,84 | 0,58 | 0,46 | 0,38 |
| | Swi | 0 | 2 | 0,74 | 0,74 | 0,82 | 0,37 | 0,37 | 0,41 | 0,04 |
| | Adh | 0 | 0,02000 | 0,012 | 0,011 | 0,0039 | 0,60 | 0,55 | 0,20 | 0,41 |
| | Act | 0 | 3 | 1,06 | 2,52 | 2,57 | 0,35 | 0,84 | 0,86 | 0,51 |

| | | | | | | | | | | |
|---|---|---|---|---|---|---|---|---|---|---|
| | Inh | 1 | 1000 | 955 | 878 | 801 | 0,95 | 0,88 | 0,80 | 0,15 |
| | Sec | 0 | 0,30 | 0,22 | 0,26 | 0,25 | 0,73 | 0,87 | 0,83 | 0,13 |
| | Da | 0 | 2 | 1,9 | 1,85 | 1,89 | 0,95 | 0,93 | 0,95 | 0,02 |
| | Di | 0 | 2 | 1,23 | 0,92 | 1,23 | 0,62 | 0,46 | 0,62 | 0,16 |
| | Ds | 0 | 2 | 0,69 | 1,38 | 1,51 | 0,35 | 0,69 | 0,76 | 0,41 |
| | Int | 0 | 1 | 0,58 | 0,3 | 0,45 | 0,58 | 0,30 | 0,45 | 0,28 |
| | Set | 0,5 | 1 | 0,7 | 0,78 | 0,69 | 0,40 | 0,56 | 0,38 | 0,18 |
| | Boy | 0 | 1 | 0,68 | 0,69 | 0,53 | 0,68 | 0,69 | 0,53 | 0,16 |
| | Dff | 0 | 0,00100 | 0,00033 | 0,00012 | 0,00019 | 0,33 | 0,12 | 0,19 | 0,21 |
| | Bgr | 0 | 2 | 1,22 | 1,38 | 1,37 | 0,61 | 0,69 | 0,69 | 0,08 |
| | Abi | 0 | 20 | 20 | 9,7 | 8,33 | 1,00 | 0,49 | 0,42 | 0,58 |
| | Pbi | 0 | 20 | 20 | 9,7 | 8,33 | 1,00 | 0,49 | 0,42 | 0,58 |
| | Lbi | 0 | 3 | 0,32 | 0,9 | 0,66 | 0,11 | 0,30 | 0,22 | 0,19 |
| | Bbi | 0 | 3 | 0,32 | 0,9 | 0,66 | 0,11 | 0,30 | 0,22 | 0,19 |
| | Rad | 2 | 4 | 2 | 2 | 2 | 0,00 | 0,00 | 0,00 | 0,00 |
| | Deg | 0 | 0,3 | 0,29 | 0,25 | 0,24 | 0,97 | 0,83 | 0,80 | 0,17 |
| | Dgr | 0 | 30000 | 4415 | 27865 | 4246 | 0,15 | 0,93 | 0,14 | 0,79 |
| | Ntr | 0 | 0,00020 | 0,00011 | 0,00013 | 0,00014 | 0,55 | 0,65 | 0,70 | 0,15 |
| | Bwi | 0 | 5 | 0,58 | 0,28 | 0,17 | 0,12 | 0,06 | 0,03 | 0,08 |
| | Ina | 0 | 100 | 5,5 | 61 | 59 | 0,06 | 0,61 | 0,59 | 0,56 |
| | uMgr | 0 | 500 | 216 | 377 | 385 | 0,43 | 0,75 | 0,77 | 0,34 |
| ***Takydromus sexlineatus*** | Egr | 0 | 0,05 | 0,039 | 0,027 | 0,036 | 0,73 | 0,43 | 0,65 | 0,30 |
| | Mgr | 0 | 20000 | 7911 | 17513 | 9730 | 0,40 | 0,88 | 0,49 | 0,48 |
| | Rep | 0 | 10 | 1,44 | 6,21 | 5,85 | 0,14 | 0,62 | 0,59 | 0,48 |
| | Swi | 0 | 2 | 0,21 | 1,69 | 1 | 0,11 | 0,85 | 0,50 | 0,74 |
| | Adh | 0 | 0,02000 | 0,019 | 0,015 | 0,015 | 0,95 | 0,75 | 0,75 | 0,20 |
| | Act | 0 | 3 | 0,78 | 1,42 | 2,59 | 0,26 | 0,47 | 0,86 | 0,61 |
| | Inh | 1 | 1000 | 220 | 383 | 353 | 0,22 | 0,38 | 0,35 | 0,16 |
| | Sec | 0 | 0,30 | 0,19 | 0,29 | 0,21 | 0,63 | 0,97 | 0,70 | 0,33 |
| | Da | 0 | 2 | 1,39 | 1,74 | 1,36 | 0,70 | 0,87 | 0,68 | 0,19 |
| | Di | 0 | 2 | 1,95 | 0,67 | 0,75 | 0,98 | 0,34 | 0,38 | 0,64 |
| | Ds | 0 | 2 | 1,31 | 1,75 | 1,33 | 0,66 | 0,88 | 0,67 | 0,22 |
| | Int | 0 | 1 | 0,46 | 0,97 | 0,42 | 0,46 | 0,97 | 0,42 | 0,55 |
| | Set | 0,5 | 1 | 0,97 | 0,72 | 0,88 | 0,94 | 0,44 | 0,76 | 0,50 |
| | Boy | 0 | 1 | 0,74 | 0,7 | 0,8 | 0,74 | 0,70 | 0,80 | 0,10 |
| | Dff | 0 | 0,00100 | 0,00018 | 0,00037 | 0,00019 | 0,18 | 0,37 | 0,19 | 0,19 |
| | Bgr | 0 | 2 | 1,36 | 0,96 | 1,19 | 0,68 | 0,48 | 0,60 | 0,20 |
| | Abi | 0 | 20 | 8,8 | 4,61 | 17,04 | 0,44 | 0,23 | 0,85 | 0,62 |
| | Pbi | 0 | 20 | 8,8 | 4,61 | 17,04 | 0,44 | 0,23 | 0,85 | 0,62 |
| | Lbi | 0 | 3 | 0,9 | 0,3 | 0,43 | 0,30 | 0,10 | 0,14 | 0,20 |

| | | | | | | | | | | |
|---|---|---|---|---|---|---|---|---|---|---|
| | Bbi | 0 | 3 | 0,9 | 0,3 | 0,43 | 0,30 | 0,10 | 0,14 | 0,20 |
| | Rad | 2 | 4 | 2 | 2 | 2 | 0,00 | 0,00 | 0,00 | 0,00 |
| | Deg | 0 | 0,3 | 0,04 | 0,11 | 0,11 | 0,13 | 0,37 | 0,37 | 0,23 |
| | Dgr | 0 | 30000 | 6176 | 27300 | 29948 | 0,21 | 0,91 | 1,00 | 0,79 |
| | Ntr | 0 | 0,00020 | 0,00016 | 0,00012 | 0,000026 | 0,80 | 0,60 | 0,13 | 0,67 |
| | Bwi | 0 | 5 | 0,37 | 0,44 | 0,33 | 0,07 | 0,09 | 0,07 | 0,02 |
| | Ina | 0 | 100 | 32,69 | 32,34 | 41,21 | 0,33 | 0,32 | 0,41 | 0,09 |
| | uMgr | 0 | 500 | 203 | 225 | 195 | 0,41 | 0,45 | 0,39 | 0,06 |
| *Chamaeleo calyptratus* | Egr | 0 | 0,05 | 0,0378 | 0,0424 | 0,0338 | 0,70 | 0,81 | 0,60 | 0,22 |
| | Mgr | 0 | 20000 | 5832 | 12961 | 13457 | 0,29 | 0,65 | 0,67 | 0,38 |
| | Rep | 0 | 10 | 9,21 | 2,69 | 7,4 | 0,92 | 0,27 | 0,74 | 0,65 |
| | Swi | 0 | 2 | 0,71 | 1,94 | 0,31 | 0,36 | 0,97 | 0,16 | 0,82 |
| | Adh | 0 | 0,02000 | 0,0197 | 0,0133 | 0,00478 | 0,99 | 0,67 | 0,24 | 0,75 |
| | Act | 0 | 3 | 0,43 | 1,13 | 1,31 | 0,14 | 0,37 | 0,43 | 0,29 |
| | Inh | 1 | 1000 | 676 | 536 | 529 | 0,68 | 0,54 | 0,53 | 0,15 |
| | Sec | 0 | 0,30 | 0,0279 | 0,22 | 0,13 | 0,09 | 0,73 | 0,43 | 0,64 |
| | Da | 0 | 2 | 1,42 | 1,39 | 1,67 | 0,71 | 0,70 | 0,84 | 0,14 |
| | Di | 0 | 2 | 2,57 | 1,38 | 0,64 | 1,29 | 0,69 | 0,32 | 0,97 |
| | Ds | 0 | 2 | 0,47 | 1,52 | 1,56 | 0,24 | 0,76 | 0,78 | 0,55 |
| | Int | 0 | 1 | 0,1 | 0,49 | 0,65 | 0,10 | 0,49 | 0,65 | 0,55 |
| | Set | 0,5 | 1 | 0,73 | 0,86 | 0,74 | 0,46 | 0,72 | 0,48 | 0,26 |
| | Boy | 0 | 1 | 0,69 | 0,71 | 0,76 | 0,69 | 0,71 | 0,76 | 0,07 |
| | Dff | 0 | 0,00100 | 0,000476 | 0,000312 | 0,000432 | 0,48 | 0,31 | 0,43 | 0,16 |
| | Bgr | 0 | 2 | 0,96 | 1,14 | 1,01 | 0,48 | 0,57 | 0,51 | 0,09 |
| | Abi | 0 | 20 | 2,45 | 2,91 | 2,91 | 0,12 | 0,15 | 0,15 | 0,02 |
| | Pbi | 0 | 20 | 2,45 | 2,91 | 2,91 | 0,12 | 0,15 | 0,15 | 0,02 |
| | Lbi | 0 | 3 | 0,44 | 0,044 | 0,64 | 0,15 | 0,01 | 0,21 | 0,20 |
| | Bbi | 0 | 3 | 0,44 | 0,044 | 0,64 | 0,15 | 0,01 | 0,21 | 0,20 |
| | Rad | 2 | 4 | 2 | 2 | 2 | 0,00 | 0,00 | 0,00 | 0,00 |
| | Deg | 0 | 0,3 | 0,27 | 0,063 | 0,25 | 0,90 | 0,21 | 0,83 | 0,69 |
| | Dgr | 0 | 30000 | 876 | 15367 | 3062 | 0,03 | 0,51 | 0,10 | 0,48 |
| | Ntr | 0 | 0,00020 | 0,0000551 | 0,000149 | 0,0001 | 0,28 | 0,75 | 0,50 | 0,47 |
| | Bwi | 0 | 5 | 4,31 | 0,55 | 0,57 | 0,86 | 0,11 | 0,11 | 0,75 |
| | Ina | 0 | 100 | 11,07 | 18,15 | 22,65 | 0,11 | 0,18 | 0,23 | 0,12 |
| | uMgr | 0 | 500 | 337 | 401 | 122 | 0,67 | 0,80 | 0,24 | 0,56 |
| ***Pogona vitticeps* (acrodont)** | Egr | 0 | 0,05 | 0,031 | 0,035 | 0,039 | 0,53 | 0,63 | 0,73 | 0,20 |
| | Mgr | 0 | 20000 | 12065 | 15579 | 12723 | 0,60 | 0,78 | 0,64 | 0,18 |
| | Rep | 0 | 10 | 0,012 | 2,54 | 1,07 | 0,00 | 0,25 | 0,11 | 0,25 |
| | Swi | 0 | 2 | 0,12 | 0,078 | 0,4 | 0,06 | 0,04 | 0,20 | 0,16 |
| | Adh | 0 | 0,02000 | 0,0088 | 0,011 | 0,0042 | 0,44 | 0,55 | 0,21 | 0,34 |

| | | | | | | | | | | |
|---|---|---|---|---|---|---|---|---|---|---|
| | Act | 0 | 3 | 1,49 | 2,39 | 2,47 | 0,49 | 0,80 | 0,82 | 0,33 |
| | Inh | 1 | 1000 | 586 | 589 | 579 | 0,59 | 0,59 | 0,58 | 0,01 |
| | Sec | 0 | 0,30 | 0,01 | 0,052 | 0,064 | 0,03 | 0,17 | 0,21 | 0,18 |
| | Da | 0 | 2 | 0,12 | 1,17 | 1,31 | 0,06 | 0,59 | 0,66 | 0,60 |
| | Di | 0 | 2 | 1,84 | 0,072 | 1,49 | 0,92 | 0,04 | 0,75 | 0,88 |
| | Ds | 0 | 2 | 0,97 | 0,11 | 1,07 | 0,49 | 0,06 | 0,54 | 0,48 |
| | Int | 0 | 1 | 0,88 | 0,47 | 0,69 | 0,88 | 0,47 | 0,69 | 0,41 |
| | Set | 0,5 | 1 | 0,83 | 0,77 | 0,54 | 0,66 | 0,54 | 0,08 | 0,58 |
| | Boy | 0 | 1 | 0,4 | 0,07 | 0,058 | 0,40 | 0,07 | 0,06 | 0,34 |
| | Dff | 0 | 0,00100 | 0,000046 | 0,00066 | 0,000084 | 0,05 | 0,66 | 0,08 | 0,61 |
| | Bgr | 0 | 2 | 1,35 | 0,91 | 1,58 | 0,68 | 0,46 | 0,79 | 0,34 |
| | Abi | 0 | 20 | 7,39 | 5,33 | 4,89 | 0,37 | 0,27 | 0,24 | 0,13 |
| | Pbi | 0 | 20 | 7,39 | 5,33 | 4,89 | 0,37 | 0,27 | 0,24 | 0,13 |
| | Lbi | 0 | 3 | 2,3 | 0,33 | 1,81 | 0,77 | 0,11 | 0,60 | 0,66 |
| | Bbi | 0 | 3 | 2,3 | 0,33 | 1,81 | 0,77 | 0,11 | 0,60 | 0,66 |
| | Rad | 2 | 4 | 4 | 4 | 4 | 1,00 | 1,00 | 1,00 | 0,00 |
| | Deg | 0 | 0,3 | 0,2 | 0,14 | 0,16 | 0,67 | 0,47 | 0,53 | 0,20 |
| | Dgr | 0 | 30000 | 29031 | 4755 | 4680 | 0,97 | 0,16 | 0,16 | 0,81 |
| | Ntr | 0 | 0,00020 | 0,00018 | 0,000066 | 0,0000055 | 0,90 | 0,33 | 0,03 | 0,87 |
| | Bwi | 0 | 5 | 3,65 | 3,42 | 4,81 | 0,73 | 0,68 | 0,96 | 0,28 |
| | Ina | 0 | 100 | 46 | 96 | 77 | 0,46 | 0,96 | 0,77 | 0,50 |
| | uMgr | 0 | 500 | 146 | 127 | 474 | 0,29 | 0,25 | 0,95 | 0,69 |
| ***Anolis carolinensis* (tricuspid)** | Egr | 0 | 0,05 | 0,035 | 0,028 | 0,027 | 0,63 | 0,45 | 0,43 | 0,20 |
| | Mgr | 0 | 20000 | 2999 | 16545 | 18318 | 0,15 | 0,83 | 0,92 | 0,77 |
| | Rep | 0 | 10 | 1,64 | 8,01 | 9,54 | 0,16 | 0,80 | 0,95 | 0,79 |
| | Swi | 0 | 2 | 0,86 | 0,26 | 0,43 | 0,43 | 0,13 | 0,22 | 0,30 |
| | Adh | 0 | 0,02000 | 0,00236 | 0,01700 | 0,01300 | 0,12 | 0,85 | 0,65 | 0,73 |
| | Act | 0 | 3 | 2,18 | 0,38 | 0,43 | 0,73 | 0,12 | 0,14 | 0,60 |
| | Inh | 1 | 1000 | 932 | 799 | 892 | 0,93 | 0,80 | 0,89 | 0,13 |
| | Sec | 0 | 0,30 | 0,084 | 0,140 | 0,140 | 0,28 | 0,47 | 0,47 | 0,19 |
| | Da | 0 | 2 | 1,41 | 1,69 | 1,03 | 0,71 | 0,85 | 0,52 | 0,33 |
| | Di | 0 | 2 | 1,44 | 0,86 | 1,19 | 0,72 | 0,43 | 0,60 | 0,29 |
| | Ds | 0 | 2 | 1,44 | 0,76 | 0,87 | 0,72 | 0,38 | 0,44 | 0,34 |
| | Int | 0 | 1 | 0,75 | 0,93 | 0,90 | 0,75 | 0,93 | 0,90 | 0,18 |
| | Set | 0,5 | 1 | 0,80 | 0,83 | 0,84 | 0,60 | 0,66 | 0,68 | 0,08 |
| | Boy | 0 | 1 | 0,85 | 0,92 | 0,98 | 0,85 | 0,92 | 0,98 | 0,13 |
| | Dff | 0 | 0,00100 | 0,00030 | 0,00014 | 0,00021 | 0,30 | 0,14 | 0,21 | 0,16 |
| | Bgr | 0 | 2 | 1,06 | 1,03 | 0,91 | 0,53 | 0,52 | 0,46 | 0,08 |
| | Abi | 0 | 20 | 3,36 | 2,84 | 2,60 | 0,17 | 0,14 | 0,13 | 0,04 |
| | Pbi | 0 | 20 | 3,36 | 2,84 | 2,60 | 0,17 | 0,14 | 0,13 | 0,04 |

| | | | | | | | | | | |
|---|---|---|---|---|---|---|---|---|---|---|
| | Lbi | 0 | 3 | 0,31 | 0,26 | 0,84 | 0,10 | 0,09 | 0,28 | 0,19 |
| | Bbi | 0 | 3 | 0,31 | 0,26 | 0,84 | 0,10 | 0,09 | 0,28 | 0,19 |
| | Rad | 2 | 4 | 2 | 2 | 2 | 0,00 | 0,00 | 0,00 | 0,00 |
| | Deg | 0 | 0,3 | 0,29 | 0,20 | 0,28 | 0,97 | 0,67 | 0,93 | 0,30 |
| | Dgr | 0 | 30000 | 28354 | 2025 | 2610 | 0,95 | 0,07 | 0,09 | 0,88 |
| | Ntr | 0 | 0,00020 | 0,0000550 | 0,0000074 | 0,0000190 | 0,28 | 0,04 | 0,10 | 0,24 |
| | Bwi | 0 | 5 | 0,97 | 3,36 | 4,79 | 0,19 | 0,67 | 0,96 | 0,76 |
| | Ina | 0 | 100 | 3,16 | 33 | 34 | 0,03 | 0,33 | 0,34 | 0,31 |
| | uMgr | 0 | 500 | 461 | 192 | 120 | 0,92 | 0,38 | 0,24 | 0,68 |

**SI References**


1. I. Salazar-Ciudad, J. Jernvall, A computational model of teeth and the developmental origins of morphological variation. *Nature* **464**, 583–586 (2010).
2. O. Hallikas, *et al.*, System - level analyses of keystone genes required for mammalian tooth development. *J. Exp. Zool. Part B Mol. Dev. Evol.* **336**, 7–17 (2021).
